# A novel mouse home cage lickometer system reveals sex- and housing-based influences on alcohol drinking

**DOI:** 10.1101/2024.05.22.595186

**Authors:** Nicholas Petersen, Danielle N. Adank, Yizhen Quan, Caitlyn M. Edwards, Anne Taylor, Danny G. Winder, Marie A. Doyle

**Affiliations:** Vanderbilt Brain Institute, Vanderbilt University School of Medicine, Nashville, TN, 37232; Vanderbilt Center for Addiction Research, Vanderbilt University School of Medicine, Nashville, TN, 37232; Department of Molecular Physiology and Biophysics, Vanderbilt University School of Medicine, Nashville, TN, 37232; Department of Neurobiology, UMass Chan Medical School, Worcester, MA, 01655

## Abstract

Alcohol use disorder (AUD) is a significant global health issue. Despite historically higher rates among men, AUD prevalence and negative alcohol-related outcomes in women are rising. Loneliness in humans has been associated with increased alcohol use, and traditional rodent drinking models involve single housing, presenting challenges for studying social enrichment. We developed LIQ PARTI (Lick Instance Quantifier with Poly-Animal RFID Tracking Integration), an open-source tool to examine home cage continuous access two-bottle choice drinking behavior in a group-housed setting, investigating the influence of sex and social isolation on ethanol consumption and bout microstructure in C57Bl/6J mice. LIQ PARTI, based on our previously developed single-housed LIQ HD system, accurately tracks drinking behavior using capacitive-based sensors and RFID technology. Group-housed female mice exhibited higher ethanol preference than males, while males displayed a unique undulating pattern of ethanol preference linked to cage changes, suggesting a potential stress-related response. Chronic ethanol intake distinctly altered bout microstructure between male and female mice, highlighting sex and social environmental influences on drinking behavior. Social isolation with the LIQ HD system amplified fluid intake and ethanol preference in both sexes, accompanied by sex- and fluid-dependent changes in bout microstructure. However, these effects largely reversed upon resocialization, indicating the plasticity of these behaviors in response to social context. Utilizing a novel group-housed home cage lickometer device, our findings illustrate the critical interplay of sex and housing conditions in voluntary alcohol drinking behaviors in C57Bl/6J mice, facilitating nuanced insights into the potential contributions to AUD etiology.

**Significance Statement:** Loneliness has been associated with increased alcohol use, and traditional rodent drinking models involve single housing, presenting challenges for studying social enrichment. Here we developed LIQ PARTI (Lick Instance Quantifier with Poly-Animal RFID Tracking Integration), an open-source group-housed lickometer system to investigate how social housing, isolation, and sex influence alcohol consumption patterns in C57Bl/6J mice. LIQ PARTI accurately identifies mouse drinking events and uncovered significant housing- and fluid-dependent differences in ethanol consumption and bout microstructure between male and female mice. Social isolation-induced changes to ethanol drinking behavior and microstructure were highly plastic, as resocialization generally reversed these changes. These findings expand on the complex interplay between sex, social isolation, and alcohol use.

## Introduction

Alcohol use disorder (AUD) is highly prevalent in the United States and worldwide (Grant et al., 2015). There are significant sex differences in alcohol use, with a lifetime AUD prevalence of 36% in men, compared to 22.7% in women (Flores-Bonilla & Richardson, 2020; Grant et al., 2015). However, the gap is narrowing due to an increase in alcohol use in women (White, 2020). Additionally, alcohol-related negative outcomes have increased overall for US adults but have increased more in women compared to men (White, 2020). In rodents, female mice and rats tend to exhibit increased ethanol consumption compared to males, especially in the commonly used C57Bl/6J mouse strain (M. A. Caruso et al., 2021; Centanni et al., 2019a; Flores-Bonilla & Richardson, 2020; Lopez & Laber, 2015; Middaugh et al., 1999; Rivera-Irizarry et al., 2023; Sneddon et al., 2019; Yu et al., 2019).

Social isolation has significant impacts on behavior and neurophysiology, including the effects of alcohol and alcohol consumption (Butler et al., 2016). Prolonged social isolation potentiates the locomotor-inducing effects of ethanol injection in mice (Päivärinta, 1990), and studies utilizing various models of ethanol consumption report that social isolation prior to ethanol exposure increases later ethanol consumption (Butler, Ariwodola, et al., 2014; Cortés-Patiño et al., 2016; Cullins & Chester, 2024; Evans et al., 2020; Lopez et al., 2011; Lopez & Laber, 2015; Panksepp et al., 2017; Sanna et al., 2011). Given that most rodent drinking models require animals to be singly housed for accurate individual intake measurements, the effects of sex and social environments on continuous access ethanol consumption, preference, and drinking patterns in ethanol-experienced mice are currently poorly understood.

The assessment of drinking in group-housed rodents has previously required the use of drinking models where ethanol consumption is recorded in discrete, limited sessions where animals are singly housed (Butler, Ariwodola, et al., 2014; Butler, Carter, et al., 2014; Cortés-Patiño et al., 2016; Cullins & Chester, 2024; Lopez et al., 2011; Lopez & Laber, 2015; Sanna et al., 2011), separated by a physical barrier (Evans et al., 2020; McKenzie-Quirk & Miczek, 2008), or recorded as an average of all animals in a cage (Panksepp et al., 2017). Specialized housing environments, such as the IntelliCage (Holgate et al., 2017) and HM2 cage (Fulenwider et al., 2021), have been valuable tools in studying the effects of social housing and enrichment on consummatory behavior. However, these systems are costly and require specialized housing. Tracking multiple animals in a cage can be performed with implanted radio-frequency identification (RFID) tags, which have been utilized in commercially available systems and by recently developed open-source behavioral devices to identify group-housed mice with video-assisted tracking in an arena (Peleh et al., 2019), mice in specialized housing (Wong et al., 2023), and rats during home cage drinking (Frie & Khokhar, 2024).

We previously developed the open-source tool LIQ HD (Lick Instance Quantifier Home cage Device) (Petersen et al., 2023), a home cage lickometer system that utilizes capacitive sensor-based technology to record undisturbed two-bottle choice mouse drinking behavior and lick microstructure in individually housed mice. Here we present LIQ PARTI (Lick Instance Quantifier with Poly-Animal RFID Tracking Integration), an open-source system designed and built based on the LIQ HD technology with the addition of RFID tags and readers for group-housed mice. Unlike currently available systems for mice, LIQ PARTI is compatible with normal vivarium home cages and can be modified to fit other cage models. Like LIQ HD, this system accurately tracks undisturbed licks and bout microstructure over at least a week, is inexpensive, and is easy to assemble with the use of 3D-printed parts and commercially available electronic components.

Using LIQ PARTI and LIQ HD, we investigated the role of sex and social environments on continuous access two-bottle choice ethanol consumption and bout microstructure in male and female C57Bl/6J mice. We find that male and female mice display significant differences in ethanol preference, drinking patterns, and drink bout microstructure in a group-housed environment that differ from those observed in an isolated setting. Group-housed females have a significantly higher preference for ethanol, while males display an undulating pattern of ethanol preference that aligns with cage bedding changes. Both male and female mice increase their ethanol preference and total consumption when transitioned into the socially isolated LIQ HD system and show significant sex- and bottle-dependent changes to bout microstructure. Drinking changes due to social isolation generally normalize following reintroduction to a group-housed environment. Altogether, we identified sex- and malleable housing-based influences on alcohol drinking using a novel open-source group-housed drinking device.

## Materials and Methods

### Animals

Male and female C57BL/6J mice (8 weeks of age) were purchased from Jackson Laboratory (#000664). Mice were allowed to habituate to the animal facility for at least 7 days before the start of experimentation. All mice were housed in Lab Products Super Mouse 750 Ventilated Cages on a standard 12hr light-dark cycle at 22-25°C with food and water available *ad libitum*. All fluid measurements conducted by experimenters took place during the light phase. All experiments were approved by the Vanderbilt University Institutional Animal Care and Use Committee (IACUC) and were carried out in accordance with the guidelines set in the Guide for the Care and Use of Laboratory Animals of the National Institutes of Health.

### LIQ PARTI Build

LIQ PARTI is built from commercially available electronic components combined with 3D-printed parts (**Supplemental Figure 1A,B**). Each cage is controlled by a single Arduino MEGA microcontroller connected to a real-time clock plus data logging shield (Adafruit), a capacitive touchscreen shield (Adafruit), a 12-channel MPR121 capacitive sensor breakout board (Adafruit), and two 125kHz RFID readers (SparkFun). Based on the previously published LIQ HD two-bottle choice system (Petersen et al., 2023), LIQ PARTI utilizes capacitive touch sensing to measure individual lick events in group-housed mice. The metal sipper for each bottle is in contact with conductive copper foil tape that is wired to the MPR121 capacitance sensor breakout board. The two RFID readers lay inside the 3D-printed housing above two small tunnels that the mice must enter to reach each sipper, which allows the device to accurately determine which mouse is actively drinking from each sipper. Both sippers can be used simultaneously without affecting data collection. Date and time are kept with the Adafruit Data Logger Shield, which also writes the cage data to a CSV file on an SD card for downstream analysis. A 2.8” Adafruit Touchscreen Shield is used to display an easy-to-use graphical user interface, allowing the user to change device settings (time, light/dark schedule, sensor sensitivity, metrics to record), mount/eject the SD card, track the total licks at each sipper throughout recordings, and start/stop/pause recordings. The screen will also prompt the user if there are any errors, such as disconnected sensors or SD card.

The Data Logger Shield and Touchscreen Shield were modified as previously described (Petersen et al., 2023). Briefly, a connection must be made between the “CS” pin to pin 7 on the Data Logger, and the connection on the “CS” solder pad must be cut. On the Touchscreen Shield, a solder bridge across the “back lite #5” pads allows for dimming of the screen during the animals’ dark cycle. Mount the Data Logger Shield and the Touchscreen Shield onto the Arduino Mega by aligning and inserting the headers. The MPR121 breakout board and RFID readers communicate with the Arduino Mega via I^2^C, which allows the microcontroller to communicate with multiple devices connected to the same pins if they have different I^2^C addresses. The MPR121 Board and RFID reader #1 (LEFT side) will remain unmodified, but to modify the I^2^C address of the RFID reader #2 (RIGHT side), create a solder bridge across the “ADR” pads. Additionally, cut the “BUZZER” jumper on both RFID reader boards and mount the ID-12LA RFID reader module onto the boards’ headers. Solder a piece of stranded wire to the “INT” pins on the RFID boards and attach them to the stripped ends of a 2-pin connector (for consistency, red wire for the LEFT RFID reader, and black wire for the RIGHT RIFD reader). Next, cut a pair of Male/Male jumper wires (Adafruit) in half and solder the stripped ends of each wire to pins 0 and 1 on the MPR121 (for consistency, red wire for 0, and black wire for 1). Using the other half of the jumper wires, solder each stripped end to a piece of 3” x ¼” conductive copper tape, secure the tape inside each sipper clip, and insert each clip into the device body (red wire for the left side and black wire for the right side). The MPR121 board and RFID readers are connected in series with Qwiic connection cables (SparkFun) and connected to the Qwiic connector on the Touchscreen Shield. Older models of the touchscreen do not have a Qwiic connector, so a Qwiic cable with breadboard jumpers can be connected directly to the Arduino Mega (blue – pin 20, yellow – pin 21, red – 5V, black – GND), secured with hot glue, and connected with an adapter (SparkFun). A ground wire is connected to the Arduino GND pin and used to ground the system during recordings. Finally, the electronics are secured in the base of the 3D-printed housing.

All 3D models were generated with Shapr3D and 3D-printed components were printed with PETG filament on an Ultimaker S5 printer as previously described (Petersen et al., 2023). PETG was chosen for its high strength, durability, chemical resistance, ease of use, and food-safe properties. Bottles were printed with translucent filament and then coated internally with food-safe epoxy resin (meets regulation requirements for repeated use under US FDA 21 CFR 177.2600) as previously described (Petersen et al., 2023). The bottles are temperature sensitive and should be washed by hand in warm water with high-quality soap (Dawn Platinum Dish Soap or Alconox Liquinox) or through a Steris Reliance 400XLS Laboratory Glassware Washer using a modified “Plastics” cycle (water temperature <50°C, no heated drying). If sterilization is required, the bottles can be sterilized via gas sterilization or UV sterilization methods. The in-cage device body was printed in pieces with black PETG and the top bottle rack was assembled to the device base with hot glue. The Arduino microcontroller case was printed with a generic PLA material.

The LIQ PARTI Arduino code was uploaded using the open-source Arduino IDE software (version 2.3.2 on MacOS). Users must first install the necessary libraries through the Arduino IDE before uploading the code. A detailed step-by-step guide, along with the Arduino code and 3D models, can be found at (https://github.com/nickpetersen93/LIQ_PARTI).

### LIQ PARTI Operation (Group-housed)

The four LIQ PARTI cages were powered by a single 12V power supply fitted with a 4-way DC power splitter. The device operates in a similar manner to LIQ HD, as previously described (Petersen et al., 2023), except that data are logged as individual time-stamped bouts (detailed below) rather than in 1-minute bins. Briefly, settings can be changed by pressing the cogwheel icon. The user should designate which side the “experimental” solution is on in the cages (e.g. sucrose, quinine, ethanol, etc.) before pressing “Start” to initiate recording. The capacitive sensor has been further optimized with improved baseline tracking and touch settings for more accurate lick detection. Default settings are pre-loaded, but users may choose to determine which sensor threshold works best for them. During recordings, users can refresh the lick display page, pause the recording, and save & quit as previously described (Petersen et al., 2023).

### RFID Tag Implant Surgery

Mice were implanted with a 125 kHz Glass Tube Radio Frequency Identification (RFID) Tag (GAO RFID Inc. Product#111001), measuring 2.12 mm in diameter and 12.0 mm in length, subcutaneously on the scruff of the neck (**Supplemental Figure 1C**). The RFID tag is specifically designed for use in animal identification with ISO 11784, ISO 11785, and EM4100 compliance and AMA 1234 and EU3456 regulation specifications. RFID tags were sterilized with a steam autoclave. For the implantation procedure, mice were induced with 3% isoflurane with oxygen and then maintained on 1.5% isoflurane for the remainder of the procedure. Mice were given a single pre-operative subcutaneous injection of 2.5mg/kg meloxicam analgesic. A small (<10mm) skin incision was made at the implantation site and the skin was carefully released from the underlying muscle using small surgical scissors. The RFID tag was inserted in parallel with the mouse’s body with either sterilized forceps or a sterilized injector syringe, and the incision was closed with a 6-0 nylon suture. Mice were allowed to recover for 7 days before the start of experimentation.

### Determination of drinking bouts

Unlike the original LIQ HD system (Petersen et al., 2023), LIQ PARTI logs each individual drinking bout rather than logging data in 1-minute bins. Thus, only licks that occur in a defined bout are recorded. The start of a drinking *bout* is defined as three licks in less than 1 second and the end of a drinking bout is defined as no licks within 1 second. *Lick number* is defined here as the number of times the animal licked the sipper during a bout, while *lick duration* is defined as the actual contact time on the sipper. *Bout duration* is defined as the time from the start of the first lick in a bout to the end of the last lick in a bout.

*Bout size* is defined as the number of licks that occur during each bout. The average *lick frequency* and *inter-lick interval* (ILI) are automatically calculated on the Arduino for each drinking bout. *Lick frequency* is defined as licks per second during bouts and calculated with the equation:

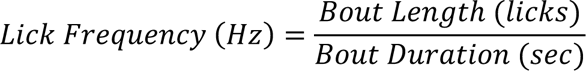

The *estimated ILI* is defined as the time between the offset of a lick and the onset of the subsequent lick and is calculated by subtracting the total bout lick duration bout duration from the bout duration and dividing by the total bout length:

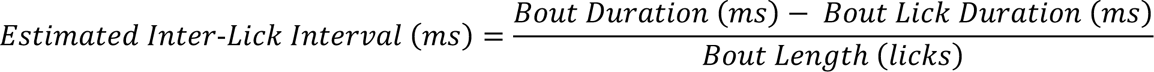

The LIQ HD systems used in this study were upgraded to log data in an identical structure to LIQ PARTI.

### LIQ HD Operation (Single-housed)

The single-housed LIQ HD system was built and operated as previously described (Petersen et al., 2023), with the exception that the capacitive sensor settings were optimized for more accurate lick detection to closely match the lick detection settings of LIQ PARTI. A detailed step-by-step guide, along with the Arduino code and 3D models, can be found at (https://github.com/nickpetersen93/LIQ_HD).

### LIQ PARTI Video Recording Validation

In a separate cohort of 10 male and 10 female mice across 4 cages, 10-minute video recordings were taken of each cage to validate RFID tracking accuracy. The tail of each mouse was marked with a different color permanent marker in order to manually score the videos. 2-3 mice per cage were randomly selected to receive a 200µL i.p. injection of hypertonic saline (2M NaCl) to induce thirst 15 minutes before video recording (Augustine et al., 2018; Pool et al., 2020). The remaining mice in each cage received an i.p. injection of 200µL normal saline (0.9% NaCl). During each video recording, both LIQ PARTI bottles contained water and the Arduino activity was simultaneously recorded using the Arduino IDE serial monitor. Videos were manually scored alongside the Arduino activity to determine the accuracy of RFID tag identification during each recorded drinking bout. A representative video was manually annotated using Premiere Rush (Adobe) (**Supplemental Video 1**). The percentage of bouts that were correctly identified was calculated by dividing the total number of recorded drinking bouts by the number of drinking bouts where the video-captured mouse (identified by tail color) matched the correct RFID number.

### Two-bottle voluntary ethanol choice task

To assess volitional ethanol intake in group-housed mice, a continuous access two-bottle choice task was used (Centanni et al., 2019b; Hodge et al., 1999; Winters et al., 2021). Specifically, 10 male and 10 female mice were group housed with LIQ PARTI devices across 4 cages with 5 mice each. Following habituation to the animal facility and RFID tag implant surgery, mice were given a 7-day LIQ PARTI habituation period, during which both bottles contained only water. No measurements were taken during this time. Mice then received 7-days of water only in both bottles, followed by 3-days of 3% ethanol and water, 7-days of 7% ethanol and water, and 4-weeks of 10% ethanol and water (Centanni et al., 2019b; Hodge et al., 1999; Winters et al., 2021). The mice were then singly housed with the LIQ HD system for 3-weeks with 10% ethanol and water. During social isolation recordings, the 20 mice were evenly divided between two separate identical LIQ HD systems to accommodate 20 cages. Lastly, the mice were returned to grouped housing with their previous cage-mates with the LIQ PARTI system for two 8-day recording periods with 10% ethanol and water. For both LIQ PARTI and LIQ HD systems, bottles and mice were manually weighed by experimenters. Cages were changed every 7 days (every Friday) throughout the study to calculate fluid intake values. Bottle placements were also swapped each time bottles were weighed to account for potential side biases.

### Statistical analysis

Data was first extracted and processed with a custom MATLAB script. Unless otherwise stated, data were binned into 1-hour bins. For the calculation of bottle preference over time, the binned data were smoothed with a moving average with a sliding window length of 6. Statistical analyses were performed with Prism 10 (GraphPad). Pearson correlation coefficients and simple linear regressions were computed for all correlation pairs. T-tests and repeated measures ANOVA with corrections for multiple comparisons were performed as indicated in the figure legends. Significance was defined as *\*p* < 0.05, *\*\*p* < 0.01, *\*\*\*p* < 0.001, and *\*\*\*\*p* < 0.0001.

### Code Accessibility

The code/software described in the paper is freely available online at (https://github.com/nickpetersen93/LIQ_PARTI). The code is available as Extended Data 1.

## Results

### LIQ PARTI accurately detects drinking bouts in a group-housed environment

We first validated the accuracy of LIQ PARTI to correctly identify RFID-tagged mice and to accurately record drinking behavior. A pilot cohort of 10 male and 10 female C57Bl/6J mice across 4 cages (5 same-sex mice each) were implanted with RFID tags under the scruff and allowed to recover and habituate to the LIQ PARTI system for at least 1 week, with both bottles containing water. These mice were then used for video-scoring validation where the tails of the mice were color-coded and a random subset of 12 mice (3 per cage) were injected with hypertonic saline to induce thirst (Augustine et al., 2018; Pool et al., 2020). The remaining mice were injected with normal saline as a control. Mice were placed in their home cage with the LIQ PARTI device, with both bottles containing water, 15 minutes after hypertonic saline injections and were video recorded alongside the Arduino IDE serial monitor to time-lock RFID scans and drinking bouts for 10 minutes. A bout is defined as a drinking event with three licks in less than 1 second and continues until no licks have occurred within 1 second. As expected, mice injected with hypertonic saline had significantly more drinking events than mice injected with normal saline, which had no drinking bouts during the recording sessions (**Supplemental Figure 2A-D**). There were 82 drinking bouts recorded throughout the 10-minute videos across the 4 cages. LIQ PARTI attributed drinking events to the correct mouse for 81/82 of drinking bouts, representing a 99% accuracy rate (**Supplemental Figure 2E**). See **Supplemental Video 1** for a fully annotated representative video recording.

To determine the accuracy of LIQ PARTI to record drinking behavior during continuous access ethanol exposure, we correlated whole-cage lickometry data with experimenter-measured bottle weights. Specifically, we correlated the total lick number, lick duration, bout number, and bout duration detected from each bottle from each recording session to the independently measured volume consumed. Lick number is defined here as the number of times the animal licked the sipper during a bout, while lick duration is defined as the actual contact time on the sipper. Bout duration is defined as the time from the start of the first lick in a bout to the end of the last lick in a bout. 10 male and 10 female mice across 4 same-sex cages were implanted with RFID tags and allowed to habituate to the LIQ PARTI devices for 1 week prior to beginning the experiment. The mice were then given 1 week of water only, 3 days of 3% ethanol and water, 1 week of 7% ethanol and water, and 4 weeks of 10% ethanol and water (Centanni et al., 2019b; Hodge et al., 1999; Winters et al., 2021). After this group-housed phase, the mice were then transferred to individual cages with the LIQ HD system for socially isolated drinking for 3 weeks with access to 10% ethanol and water before being returned to the group-housed LIQ PARTI system for 2 weeks, again with access to 10% ethanol and water (**Figure 1A**). Bottles and mice were manually weighed by experimenters weekly (except during 3% ethanol exposure when they were weighed after 3 days), and the measured volume change during group-housed recording sessions was correlated to the various LIQ parameters to determine the accuracy of whole-cage intake behavior (**Supplemental Figure 2F-M**). We observed a strong, significant correlation between total lick number and volume change (*R^2^* = 0.9357, *F*_(1,54)_ = 785.5, *p*<0.0001) (**Supplemental Figure 2F**), total lick duration and volume change (*R^2^* = 0.8724, *F*_(1,54)_ = 369.1, *p*<0.0001) (**Supplemental Figure 2G**), total bout number and volume change (*R^2^*= 0.7742, *F*_(1,54)_ = 185.2, *p*<0.0001) (**Supplemental Figure 2H**), and total bout duration and volume change (*R^2^* = 0.9446, *F*_(1,54)_ = 921.5, *p*<0.0001) (**Supplemental Figure 2I**). We observed a strong, significant correlation between the preference score calculated by the total lick number and the preference by volume change (*R^2^* = 0.9809, *F*_(1,26)_ = 1339, *p*<0.0001) (**Supplemental Figure 2J**), preference by total lick duration and preference by volume change (*R^2^*= 0.9668, *F*_(1,26)_ = 756.1, *p*<0.0001) (**Supplemental Figure 2K**), preference by total bout number and preference by volume change (*R^2^* = 0.9713, *F*_(1,26)_ = 881.2, *p*<0.0001) (**Supplemental Figure 2L**), and preference by total bout duration and preference by volume change (*R^2^* = 0.9860, *F*_(1,26)_ = 1837, *p*<0.0001) (**Supplemental Figure 2M**). The fitted regression model for the correlation between lick number and volume change is Y = 719.5*X − 3153 (slope 95% CI, 668.0-770.9). Thus, on average, the mice took 720 licks to drink 1 mL of fluid or 1.4 μL per lick. This is consistent with our previously published LIQ HD system (Petersen et al., 2023). Lick number will be used throughout this paper as a proxy for fluid consumption. Altogether, LIQ PARTI identifies RFID-tagged mice and drinking behavior with high fidelity.

**Figure 1.**
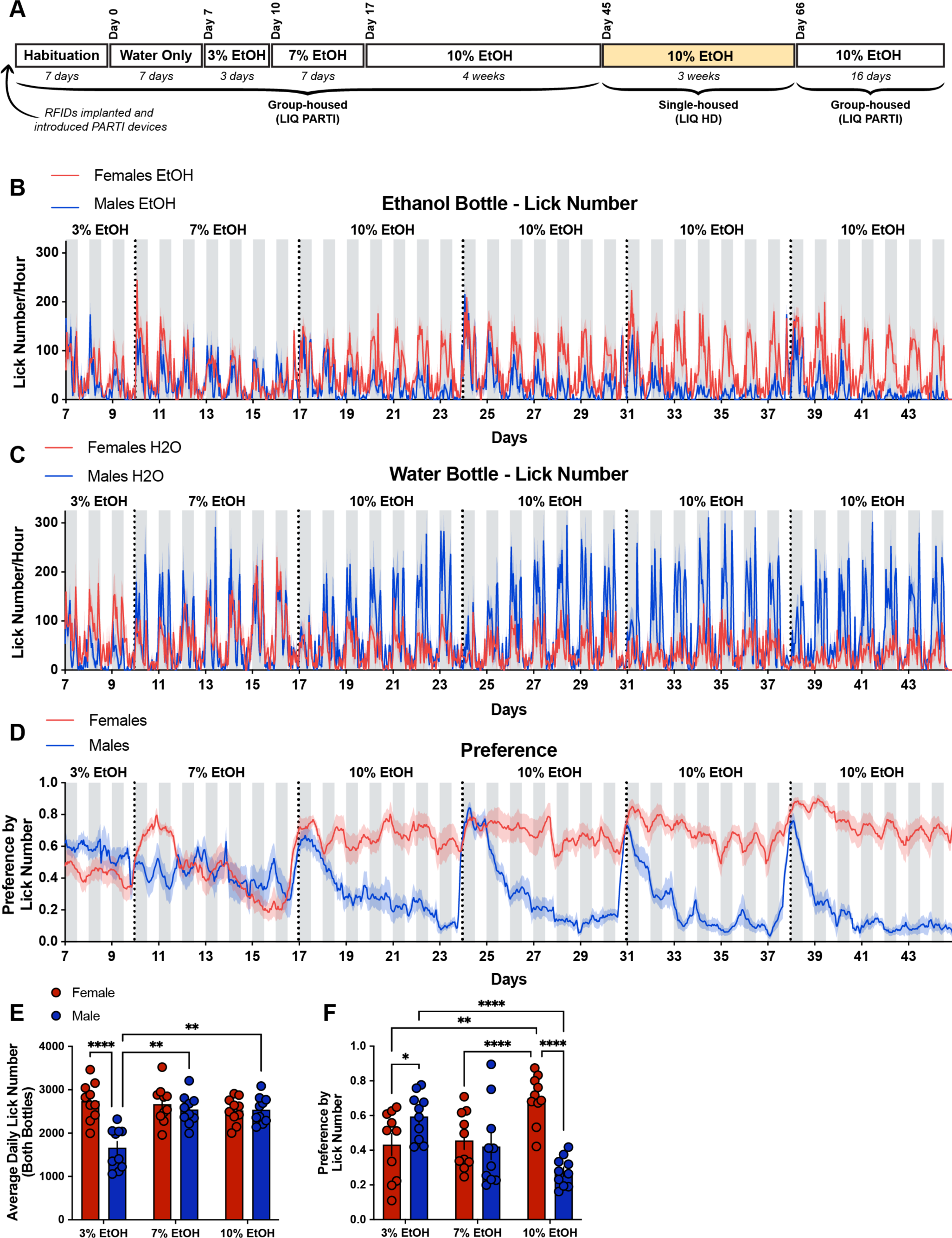
Female mice form a significantly higher preference for ethanol compared to males in a group-housed continuous access two-bottle choice task, and males display an undulating pattern of ethanol preference that aligns with cage-changing days. **A**, Timeline of experimental setup. 10 male and 10 female mice across 4 cages were implanted with RFID tags under the scruff of the neck and given 7 days to recover and habituate with the LIQ PARTI devices. Mice were then exposed to a continuous two-bottle choice paradigm with water only for 1 week, 3% ethanol and water for 3 days, 7% ethanol and water for 1 week, and 10% ethanol and water for 4 weeks. Mice were then socially isolated with the LIQ HD system for 3 weeks and then returned to group housing with LIQ PARTI for 16 days. Lick number per hour from male and female mice at the ethanol bottle (**B**) and water bottle (**C**) throughout ethanol exposure with LIQ PARTI prior to social isolation. **D**, Ethanol preference score calculated by lick number in male and female mice throughout ethanol exposure with LIQ PARTI prior to social isolation. **E**, Male and female mice show similar levels of average total lick number per day (both bottles) throughout ethanol exposure, except for during 3% ethanol and water where males displayed significantly fewer licks than females (repeated measures two-way ANOVA, with the Geisser–Greenhouse correction and Tukey’s multiple comparisons test). **F**, Male mice have a significantly higher preference for ethanol during exposure to 3% ethanol and water, but they significantly reduce their preference through exposure to 10% ethanol (repeated measures two-way ANOVA, with the Geisser–Greenhouse correction and Tukey’s multiple comparisons test). Female mice have a significantly higher preference for ethanol than males during exposure to 10% ethanol with LIQ PARTI prior to social isolation. The solid lines (**B-D**) represent the mean lick number in 1-h bins, and the red/blue shaded areas represent ±SEM. Gray shaded areas represent the dark cycle and vertical dotted lines represent when cages were cleaned and bottle positions were swapped. Error bars (**E,F**) represent ±SEM. (*n* = 10 mice per group and reported as individual mice). *\*p* < 0.05, *\*\*p* < 0.01, *\*\*\*\*p* < 0.0001.

### Male and female mice display significant diPerences in ethanol preference, drinking patterns, and microstructure in a group-housed ad libitum home cage drinking environment

In the same cohort, we next utilized LIQ PARTI to evaluate ethanol drinking behavior of individual C57Bl/6J mice during the continuous access voluntary two-bottle choice paradigm (Centanni et al., 2019b; Hodge et al., 1999; Winters et al., 2021) in a group-housed environment. As described above, following RFID implantation and habituation to the LIQ PARTI system, the mice were given access to water in both bottles for 1 week (**Supplemental Figure 3**). We did not observe any significant differences in the total number of licks (**Supplemental Figure 3A,B,D**) or bottle preference (**Supplemental Figure 3C,E**) between the male and female group-housed mice, suggesting male and female mice consume water at similar rates when given continuous access to two water bottles. Female mice displayed a significantly higher average lick duration (**Supplemental Figure 3I**) and average lick frequency (licks per second) (**Supplemental Figure 3J**) than males, with no significant differences in average daily bout number, bout size (licks per bout), bout duration, or inter-lick-interval (ILI, the time between the end of one lick to the start of another) (**Supplemental Figure 3F,G,H,K**).

The group-housed mice were then given access to 3% ethanol and water for 3 days, 7% ethanol and water for 1 week, and 10% ethanol and water for 4 weeks (**Figure 1**). Throughout this group-housed period, male and female mice had a similar number of average total licks per day (both bottles), except during access to 3% ethanol where male mice had significantly fewer licks (**Figure 1B,C,E**). However, female mice developed a significantly higher preference for ethanol during access to 10% ethanol compared to 3% and 7% ethanol and compared to males at 10% ethanol (**Figure 1D,F**). Interestingly, males showed a gradual decrease in the average preference for ethanol throughout ethanol exposure, with significantly higher ethanol preference during 3% ethanol compared to females and compared to themselves at 10% ethanol. Dirty cages were exchanged each week throughout the experiment, which corresponded to within-week fluctuations in the preference for ethanol in male mice only (**Figure 1D**). With the advantage of high temporal resolution with LIQ PARTI, we observed that male mice displayed an undulating pattern of ethanol preference with a significantly higher preference for ethanol, at similar levels to the females, at the beginning of each recording week period. These spikes in male ethanol preference occur in tandem with bedding changes and diminish over 24-48 hours.

In addition to differences in ethanol preference between male and female group-housed mice, the animals showed significant differences in bout microstructure that were ethanol exposure-, sex-, and fluid (ethanol or water)-dependent (**Figure 2**). Compared to water-only access, during exposure to 10% ethanol, male mice significantly decreased their average daily bout number at the ethanol bottle while female mice increased their average daily bout number at the ethanol bottle and decreased at the water bottle (**Figure 2A**). Compared to the male mice during 10% ethanol exposure, female mice had a significantly lower average daily bout number at the water bottle and a significantly higher bout number at the ethanol bottle (**Figure 2A**). Male mice significantly increased their bout size at the ethanol bottle and bout duration at the water and ethanol bottle compared to the water-only week (**Figure 2B,C**). Female mice did not significantly change their bout size or duration at either bottle in response to ethanol exposure and had significantly smaller and shorter bouts at both bottles compared to males during 10% ethanol (**Figure 2B,C**). Both male and female mice displayed a decrease in the average lick duration at the ethanol bottle compared to the water bottle during 10% ethanol exposure and compared to the water week (**Figure 2D**). Female mice also had a decreased average lick duration at the water bottle compared to male mice during exposure to 10% ethanol (**Figure 2D**). Male mice significantly decreased their lick frequency at the water bottle during exposure to 10% ethanol and had a significantly slow lick frequency during ethanol exposure at the water bottle compared to females (**Figure 2E**). In contrast, females significantly increased their lick frequency at the water bottle during exposure to 10% ethanol (**Figure 2E**). The ILI of male mice did not significantly change at either bottle in response to ethanol exposure, but female mice showed a significant increase in the ILI at the ethanol bottle compared to the water week and compared to the water bottle during 10% ethanol exposure (**Figure 2F**). Altogether, these data demonstrate significant changes to bout microstructure following exposure to chronic, continuous access ethanol in group-housed mice that are dependent on both sex and the fluid that is being consumed (**Figure 2G**).

**Figure 2.**
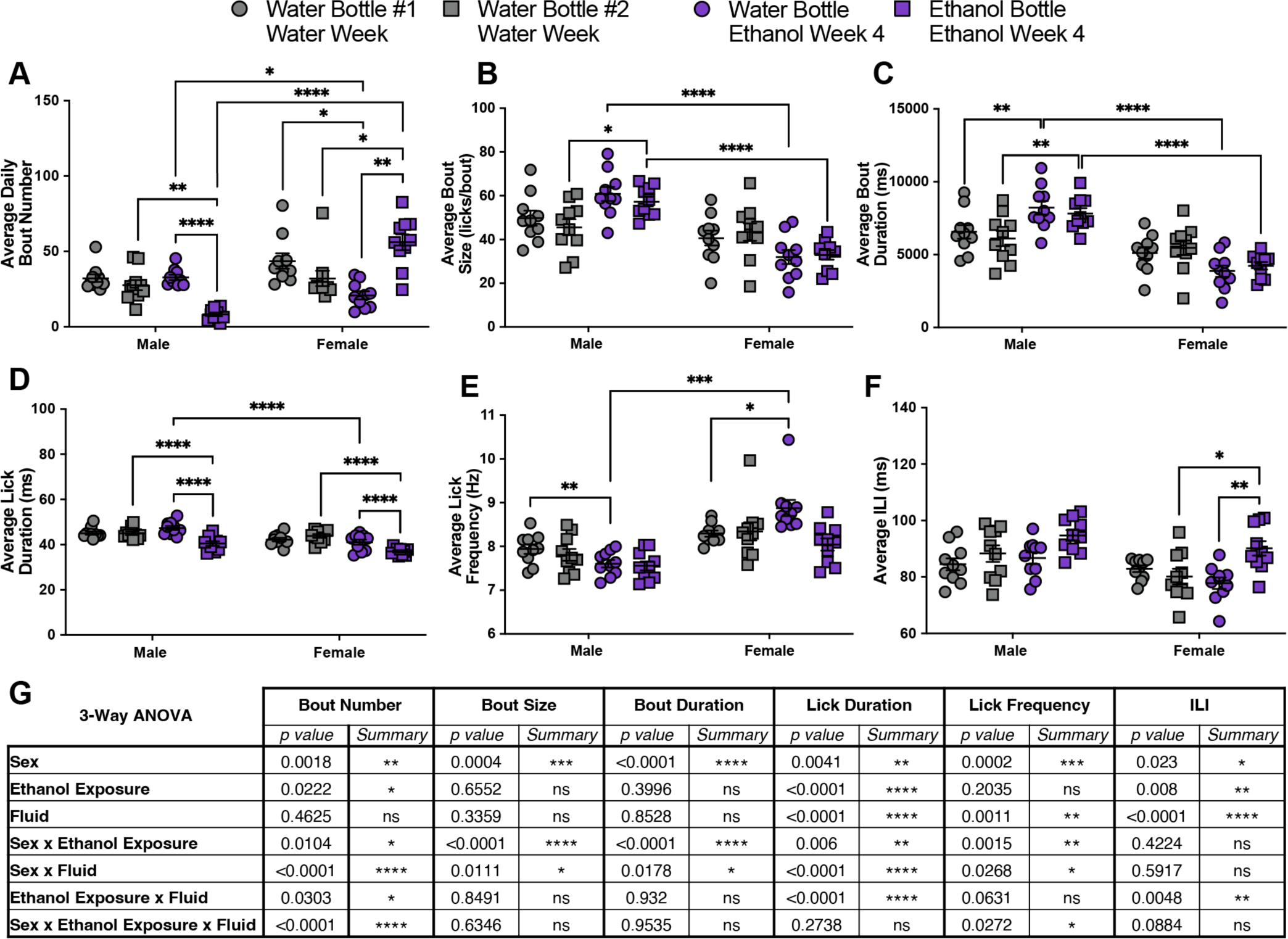
Exposure to ethanol significantly alters drinking microstructure in a sex- and bottle-specific manner in group-housed mice. Average daily bout number (**A**), bout size (**B**), bout duration (**C**), lick duration (**D**), lick frequency (**E**), and ILI (**F**) at the water and ethanol bottles for males and females during the water week (both bottles with water) and during the fourth week of 10% ethanol exposure (repeated measures three-way ANOVA, with the Geisser–Greenhouse correction and Šidák multiple comparisons test). **G**, 3-way ANOVA results for the microstructure parameters showing the main effects of sex, ethanol exposure, fluid, and interactions. Error bars represent ±SEM. (*n* = 10 mice per group and reported as individual mice). *\*p* < 0.05, *\*\*p* < 0.01, *\*\*\*p* < 0.001, *\*\*\*\*p* < 0.0001.

### Social isolation enhances ethanol preference and alters drinking microstructure in a sex- and bottle-specific manner

To investigate the effect of social isolation on alcohol-drinking behavior and bout microstructure in male and female mice, the animals were transferred into individual cages equipped with the LIQ HD system (Petersen et al., 2023). The LIQ HD code was modified to enhance the lick detection of the capacitive sensor and to match the bout detection parameters to the LIQ PARTI system. To validate these changes, we correlated the total lick number at each bottle with the experimenter measured volume change as well as the preference calculated by lick number with the preference by volume change. The data from the 3 weeks of social isolation were combined for these correlations. We observed a strong, significant correlation between total lick number and volume change (*R^2^* = 0.9509, *F*_(1,117)_ = 2268, *p*<0.0001) (**Supplemental Figure 4A**), and between preference by total lick number and preference by volume change (*R^2^*= 0.8731, *F*_(1,58)_ = 399.1, *p*<0.0001) (**Supplemental Figure 4B**). The fitted regression model for the correlation between lick number and volume change is Y = 737.7*X − 114.9 (slope 95% CI, 707.0-768.4), corresponding to an average of 738 licks per 1 mL of fluid or 1.4 μL per lick, which is consistent with our previous LIQ HD data (Petersen et al., 2023) and LIQ PARTI. The slopes of the fitted regression of total lick number versus volume change for LIQ HD and LIQ PARTI did not significantly differ (**Supplemental Figure 4C**).

Ethanol drinking behavior substantially increased in both male and female mice when transferred to social isolation (**Figure 3**). Compared to the fourth week of group-housed 10% ethanol, both male and female mice significantly increased their average total lick number per day across both the ethanol and the water bottle combined, and the average total lick number per day did not differ by sex (**Figure 3A,B,D**). Additionally, the preference for ethanol significantly increased during social isolation in both males and females, but to a greater magnitude in males (**Figure 3E**). However, this may be due to a ceiling effect in the females. Consistent with ethanol preference during group housing, females have a significantly greater preference for ethanol compared to males during social isolation (**Figure 3E**).

**Figure 3.**
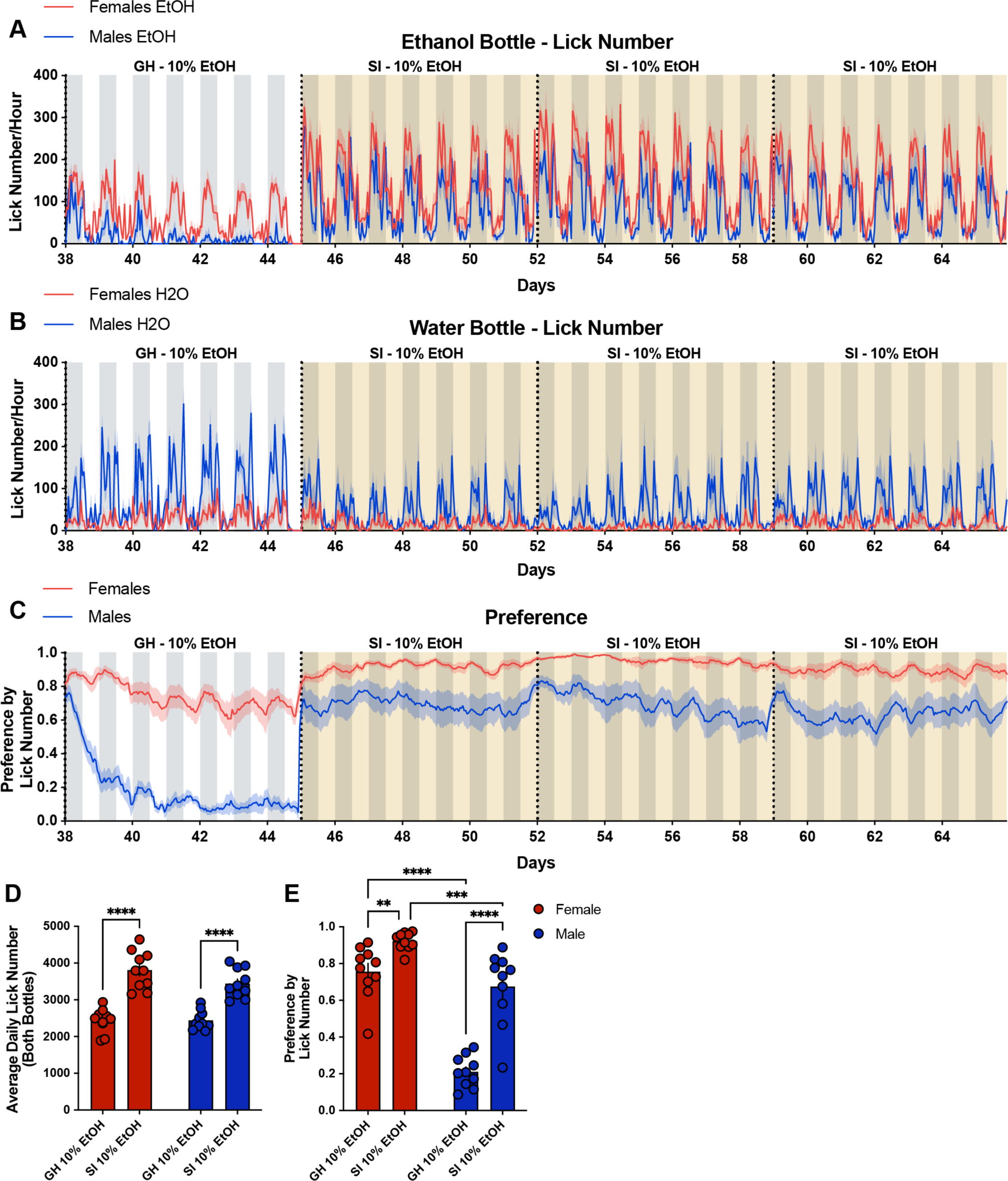
Social isolation in the LIQ HD system significantly increases ethanol preference and total consumption in male and female mice and changes the overall ethanol preference pattern over time in males. Lick number per hour from male and female mice at the ethanol bottle (**A**) and water bottle (**B**) throughout the last week of 10% ethanol exposure with the group-housed (GH) LIQ PARTI system and through social isolation (SI) with LIQ HD with 10% ethanol and water. **C**, Ethanol preference score calculated by lick number in male and female mice throughout the last week of 10% ethanol exposure with the group-housed LIQ PARTI system and through social isolation with LIQ HD with 10% ethanol and water. **D**, Both male and female mice display an increase in the average total lick number per day (both bottles) during social isolation with no significant difference between male and female mice (repeated measures two-way ANOVA, Šidák multiple comparisons test). **E**, Both male and female mice have an increased preference for ethanol during social isolation and female mice have a higher ethanol preference than males during group housing with LIQ PARTI and social isolation with LIQ HD (repeated measures two-way ANOVA, Šidák multiple comparisons test). The solid lines (**A-C**) represent the mean lick number in 1-h bins, and the red/blue shaded areas represent ±SEM. Gray shaded areas represent the dark cycle and vertical dotted lines represent when cages were cleaned and bottle positions were swapped. Yellow shaded regions indicate periods of social isolation with LIQ HD. Error bars (**D,E**) represent ±SEM. (*n* = 10 mice per group and reported as individual mice). *\*\*p* < 0.01, *\*\*\*p* < 0.001, *\*\*\*\*p* < 0.0001.

Ethanol drinking bout microstructure was significantly altered in a housing-, sex-, and fluid-dependent manner. The average daily bout number at the ethanol both significantly increased for both male and female mice during social isolation compared to grouped housing, with female mice displaying significantly more average daily bouts at the ethanol bottle compared to males during grouped housing and social isolation (**Figure 4A**) Prior to social isolation, male mice had significantly larger and longer bouts at both ethanol and water bottles compared to female mice (**Figure 4B,C**). Following social isolation, male mice significantly decreased their bout size and bout durations at both the water and ethanol bottles to comparable levels to the female mice (**Figure 4B,C**). Female mice significantly decreased their bout size and duration at the water bottle and decreased bout duration at the ethanol bottle. Both male and female mice had significantly larger bouts at the ethanol bottle compared to the water bottle during social isolation, and male mice had increased bout duration at the ethanol bottle (**Figure 4B,C**). Male and female mice also showed an increased average lick duration at both the water and ethanol bottles compared to the group-housed setting (**Figure 4D).** Additionally, both sexes had significantly longer lick duration at the water bottle compared to the ethanol bottle during both group housing and social isolation, but to a greater magnitude during isolation (**Figure 4D**). Average lick frequency and ILI did not significantly change in male mice in response to social isolation (**Figure 4E,F**). However, female mice significantly reduced their lick frequency at the water bottle and had a significantly higher lick frequency at the ethanol bottle in comparison to the water bottle during social isolation (**Figure 4F**). While female mice had a higher ILI at the ethanol bottle compared to the water bottle during group housing, the opposite effect was seen during social isolation. Female mice had an increased ILI at the water bottle and a decreased ILI at the ethanol bottle in response to social isolation (**Figure 4F**). Taken together, social isolation leads to robust changes in bout microstructure in male and female ethanol-experienced mice that vary based on sex and fluid (**Figure 4G**).

**Figure 4.**
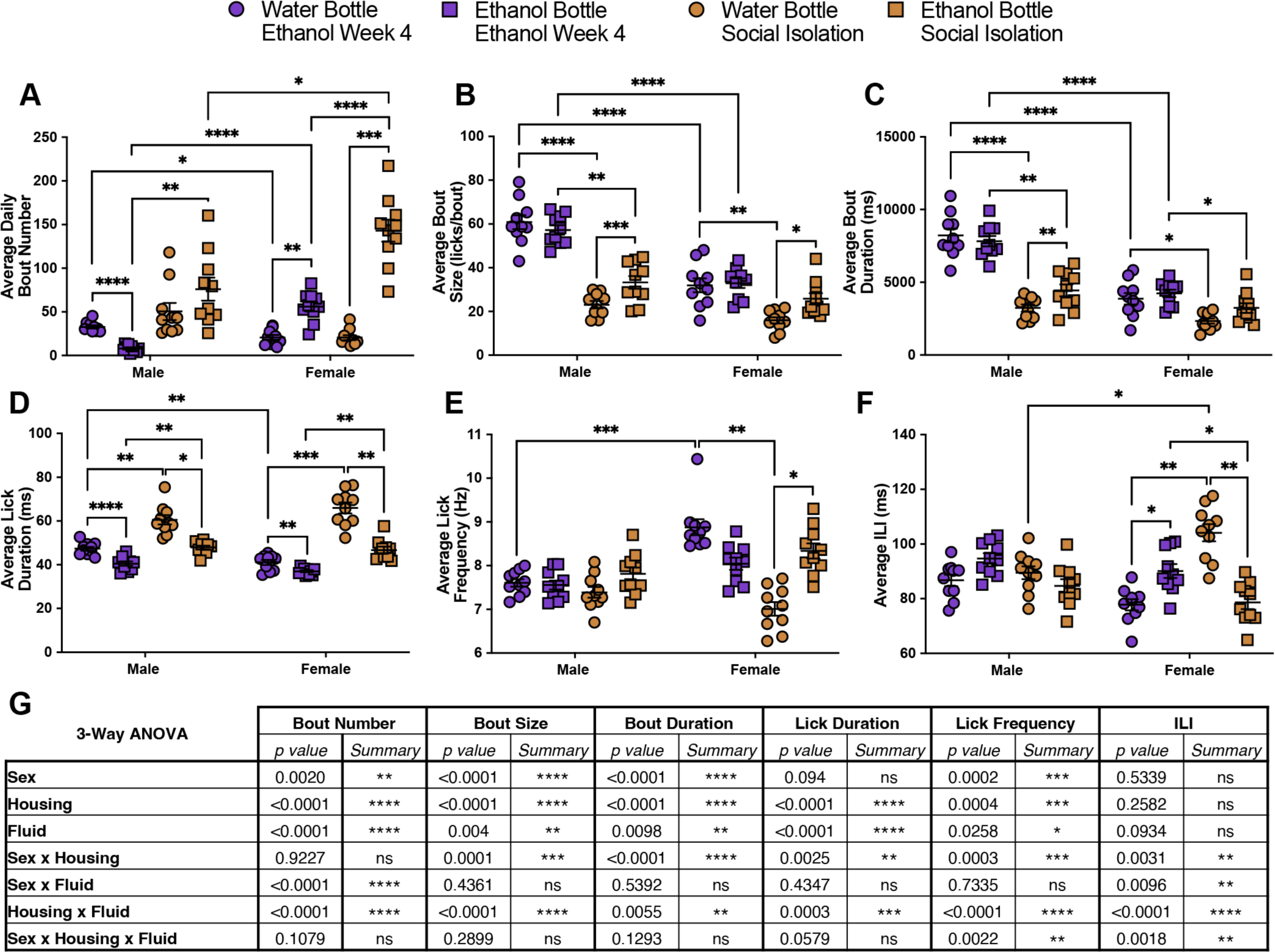
Social Isolation significantly alters drinking microstructure in a sex and bottle-specific manner. Average daily bout number (**A**), bout size (**B**), bout duration (**C**), lick duration (**D**), lick frequency (**E**), and ILI (**F**) at the water and ethanol bottles for males and females during the fourth week of group-housed 10% ethanol exposure and during social isolation with 10% ethanol (averaged across the 3 weeks) (repeated measures three-way ANOVA, with the Geisser–Greenhouse correction and Šidák multiple comparisons test). **G**, 3-way ANOVA results for the microstructure parameters showing the main effects of sex, housing, fluid, and interactions. Error bars represent ±SEM. (*n* = 10 mice per group and reported as individual mice). *\*p* < 0.05, *\*\*p* < 0.01, *\*\*\*p* < 0.001, *\*\*\*\*p* < 0.0001.

### Mice returned to a group-housed environment following social isolation displayed sex-specific changes in ethanol preference compared to pre-social isolation

Following social isolation with the LIQ HD system, the mice were returned to group housing with their original cage mates and the LIQ PARTI systems to investigate whether social isolation-induced changes to drinking behavior persist after social reintegration. The mice were group-housed for two recording periods each 8 days long. A cage of male mice was excluded from the analysis of the second recording period due to user error when starting the system. However, bottle measurement data closely recapitulate the same conclusions (**Figure 5**). Compared to the last group-housed 10% ethanol week pre-social isolation, female mice decreased their average total lick number per day (both bottles) back to pre-social isolation levels and further decreased during the second recording period of group housing post-social isolation (**Figure 6A,B,D; Figure 5E,F**). Male mice continued to show an increase in the average total licks per day during the first period post-social isolation but returned to pre-social isolation levels during the second recording period (**Figure 6A,B,D; Figure 5E,F**). Additionally, male mice showed a greater average total lick number per day compared to females during the first week of post-social isolation group housing but returned to comparable levels by the second recording period (**Figure 6A,B,D; Figure 5E,F**). Ethanol preference was significantly lower overall in male mice versus female mice pre- and post-social isolation (**Figure 6C,E**; **Figure 5A-D,G,H**), as previously reported (**Figure 1D,E**). Male mice immediately reverted to pre-social isolation ethanol preference and restarted the same undulating pattern of preference corresponding to cage changes (**Figure 6C,E; Figure 5A-D,G,H**). In contrast, female mice had a significantly lower ethanol preference post-social isolation compared to pre-social isolation but reverted to a similar level of preference by the second recording period (**Figure 6C,E**; **Figure 5A-D,G,H**).

**Figure 5.**
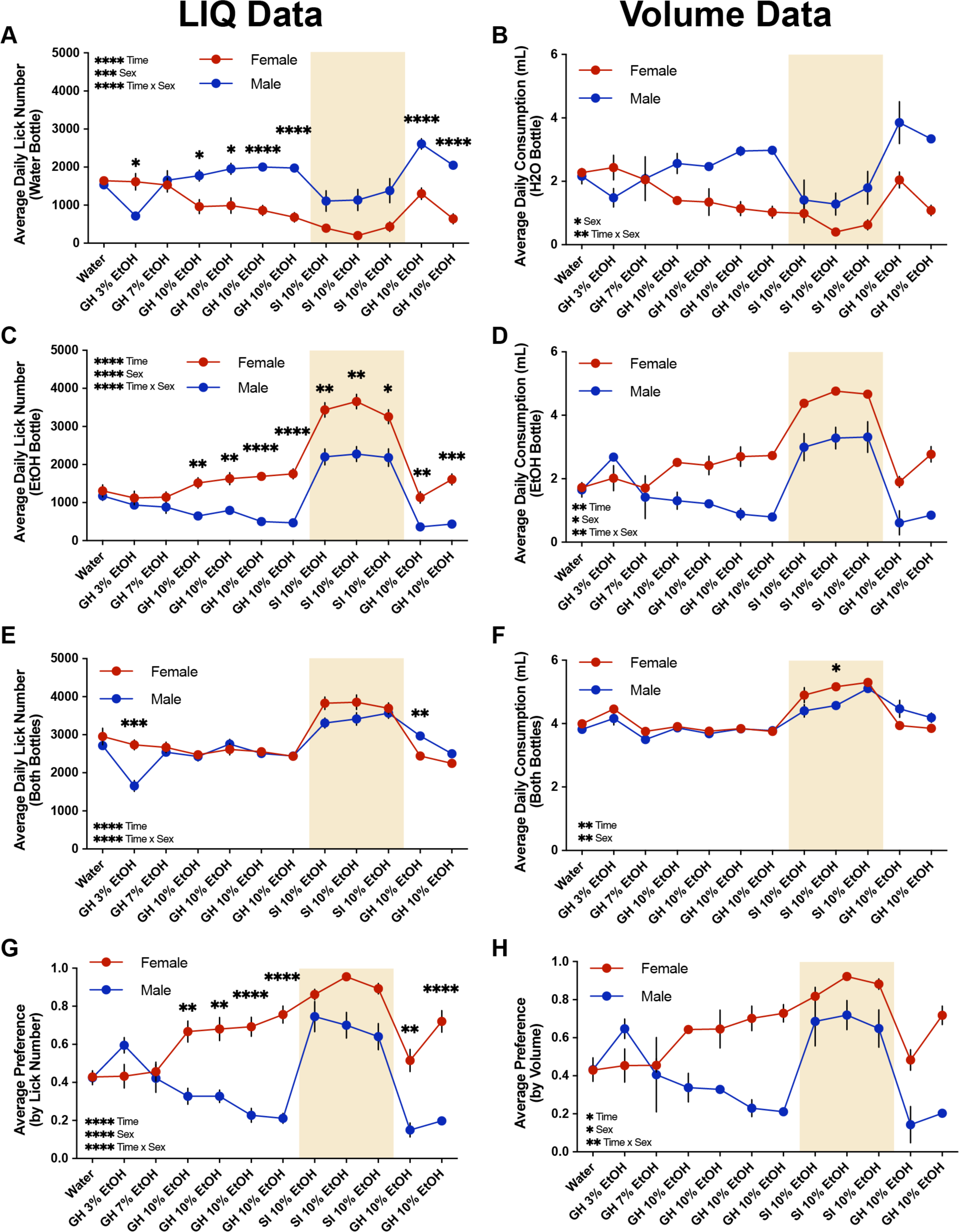
Summary of LIQ and volume measurement data throughout the two-bottle choice drinking paradigm. Average daily lick number (A) and average daily consumption (B) at the water bottle. Average daily lick number (C) and average daily consumption (D) at the ethanol bottle. Average daily lick number (E) and average daily consumption (F) at both bottles combined. Average preference by lick number (G) and average preference by average volume consumed (H). Volume measurement data was calculated as the average across all animals per cage. (LIQ data: repeated measures mixed-effects model, with the Geisser–Greenhouse correction and Šidák multiple comparisons test. Volume data: repeated measures two-way ANOVA, with the Geisser–Greenhouse correction and Šidák multiple comparisons test). Yellow shaded regions indicate periods of social isolation with LIQ HD. Error bars represent ±SEM. (LIQ data *n* = 10 mice per group, except males post-SI week 2 n = 5, and are reported as individual mouse values; Volume data *n* = 2 cages per group and are reported as whole-cage values). *\*p* < 0.05, *\*\*p* < 0.01, *\*\*\*p* < 0.001, *\*\*\*\*p* < 0.0001.

**Figure 6.**
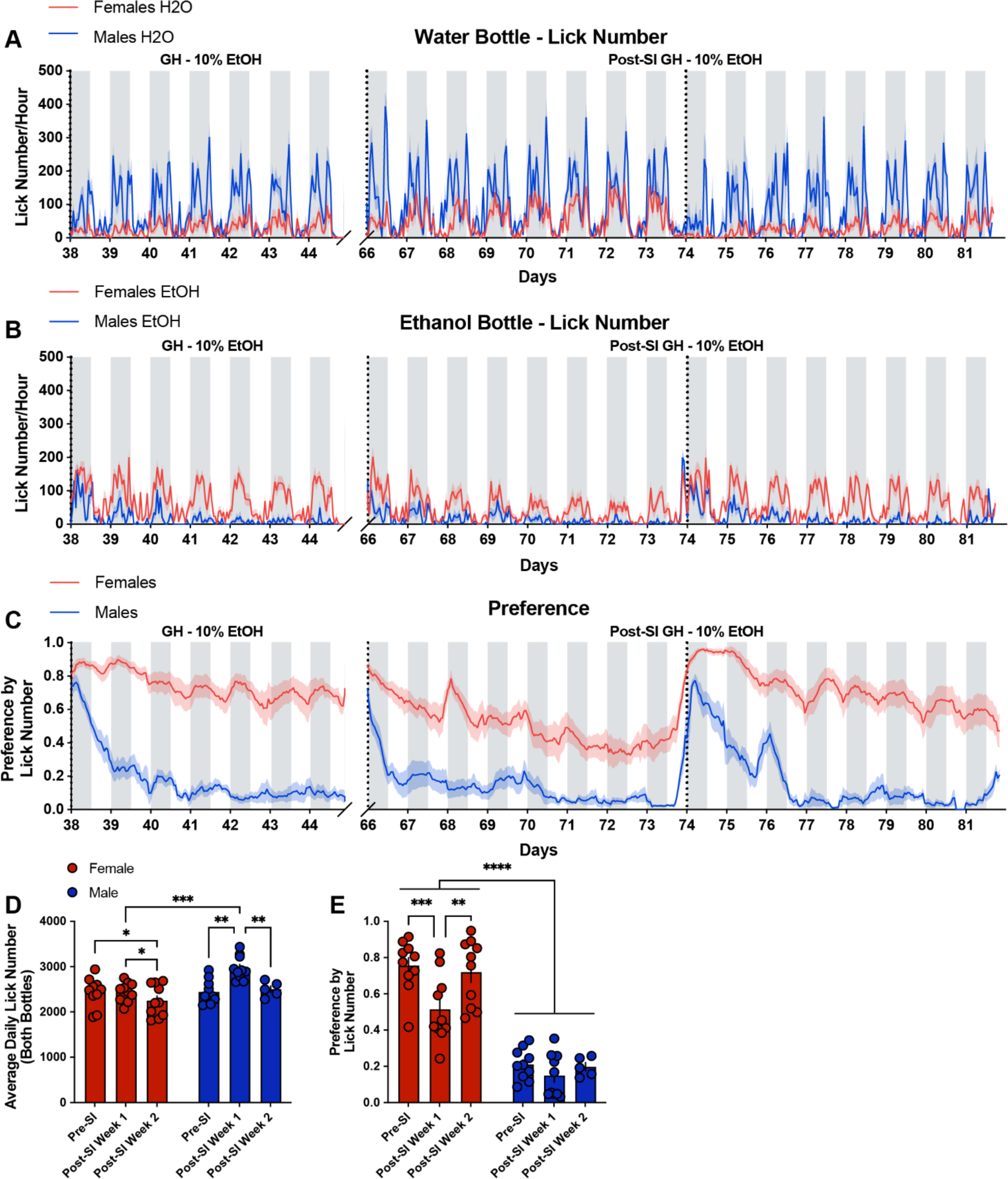
The average lick number per day modestly decreases in females and returns to baseline in males, and ethanol preference returns to baseline in both males and females during the return to a group-housed setting post-isolation compared to the last previous group-housed week. Lick number per hour from male and female mice at the ethanol bottle (**A**) and water bottle (**B**) throughout the last week of 10% ethanol exposure with the group-housed LIQ PARTI system and through post-social isolation with 10% ethanol and water. **C**, Ethanol preference score calculated by lick number in male and female mice throughout the last week of 10% ethanol exposure with the group-housed LIQ PARTI system and through post-social isolation with 10% ethanol and water. Male mice resume the previously observed pattern of undulating ethanol preference that corresponds with cage-changing days. **D**, Female mice show no significant difference in the average total licks per day (both bottles) during the first post-social isolation recording session, but modestly decrease during the second session. Male mice display a significantly greater average total lick number per day during with first recording session post-social isolation with no significant difference during the second session compared to the last week of pre-social isolation 10% ethanol. (repeated measures mixed-effects model, Tukey’s multiple comparisons test). **E**, Female mice have a significantly lower ethanol preference during the first week of post-social isolation compared to the last week of pre-social isolation with 10% ethanol. Ethanol preference returns to pre-social isolation levels in females during the second post-social isolation recording session. Male mice immediately return to pre-social isolation ethanol preference post-social (repeated measures mixed-effects model, Tukey’s multiple comparisons test). The solid lines (**A-C**) represent the mean lick number in 1-h bins, and the red/blue shaded areas represent ±SEM. Gray shaded areas represent the dark cycle and vertical dotted lines represent when cages were cleaned and bottle positions were swapped. Error bars (**D,E**) represent ±SEM. (*n* = 10 mice per group, except males post-SI week 2 n = 5, and reported as individual mice). *\*p* < 0.05, *\*\*p* < 0.01, *\*\*\*p* < 0.001, *\*\*\*\*p* < 0.0001.

Bout microstructure parameters mostly reverted to pre-social isolation values with some exceptions (**Figure 7; Supplemental Figure 5**). While significant differences between sexes remained, male mice showed no significant difference in the average daily bout number at the water or ethanol bottles post-social isolation compared to pre-social isolation (**Figure 7A; Supplemental Figure 5A**). The average daily bout number at the water bottle for females also showed no significant difference after social isolation compared to pre-social isolation, but female mice showed a moderate decrease in the average daily bout number at the ethanol bottle during post-social isolation (**Figure 7A; Supplemental Figure 5A**). Male mice displayed a significant decrease in bout size and bout duration at the ethanol bottle compared to pre-social isolation and female mice reverted to pre-social isolation values (**Figure 7B,C; Supplemental Figure 5B,C**). In contrast, male mice reverted to pre-social isolation average lick duration values at both bottles while female mice showed a small, but significant increase in the average lick duration at both the water and ethanol bottles post-social isolation (**Figure 7D; Supplemental Figure 5D**). Lick frequency and ILI remained unchanged in the male mice (**Figure 7 E,F; Supplemental Figure 5E,F**). Female mice had shown significant differences in lick frequency and ILI between the water and ethanol bottles during social isolation (**Figure 4E,F**), but showed no significant differences between the bottles during post-social isolation group housing with a significant decrease in lick frequency at the water bottle post-social isolation compared to pre-social isolation (**Figure 7E,F; Supplemental Figure 5E,F**). These data highlight that while social isolation induces substantial changes in alcohol drinking behavior in male and female mice, these behaviors are plastic and can essentially reverse following reintroduction to a social environment.

**Figure 7.**
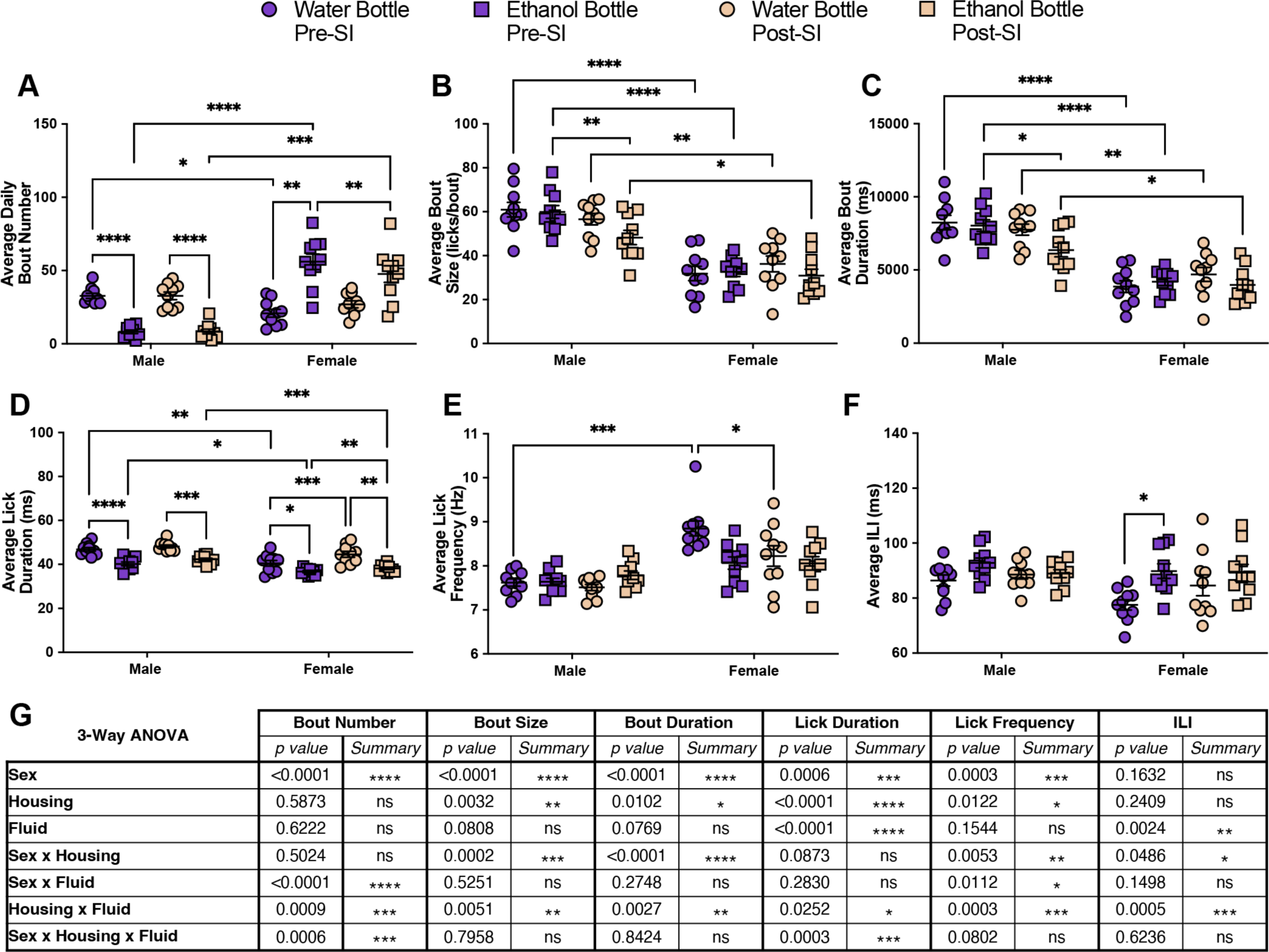
Social reintegration significantly alters drinking microstructure in a sex- and bottle-specific manner, which largely reverts to pre-social isolation values. Average daily bout number (**A**), average bout size (**B**), bout duration (**C**), lick duration (**D**), lick frequency (**E**), and ILI (**F**) at the water and ethanol bottles for males and females during fourth week of group-housed 10% ethanol exposure (pre-SI) and during post-social isolation with 10% ethanol (post-SI, averaged across the 2 recording periods) (repeated measures three-way ANOVA, with the Geisser–Greenhouse correction and Šidák multiple comparisons test). **G**, 3-way ANOVA results for the microstructure parameters showing the main effects of sex, housing, fluid, and interactions. Error bars represent ±SEM. (*n* = 10 mice per group and reported as individual mice). *\*p* < 0.05, *\*\*p* < 0.01, *\*\*\*p* < 0.001, *\*\*\*\*p* < 0.0001.

## Discussion

We utilized LIQ PARTI, a novel mouse home cage lickometer system to record two-bottle choice drinking behavior in a group-housed environment, to investigate sex differences and the effects of social environments on continuous access ethanol drinking and bout microstructure in C57Bl/6J mice. LIQ PARTI is built upon the same technology in our previously published LIQ HD system (Petersen et al., 2023) that uses capacitive-based touch sensors to detect single lick events. Each cage is controlled by a single Arduino MEGA microcontroller connected to a real-time clock plus data logging shield, a touchscreen shield, a MPR121 capacitive sensor breakout board, and two 125kHz RFID readers. The two RFID readers lay inside the 3D-printed housing above two small tunnels that the mice must enter to reach each sipper, which allows the device to read an implanted RFID tag to determine which mouse is actively drinking from each sipper with 99% accuracy. The Arduino microcontroller records each drinking bout with a timestamp and automatically logs the number of licks, the bout duration, the total lick duration (contact time), the average lick frequency, and the average ILI of each bout to an SD card. We show that in addition to the high accuracy of RFID identification, LIQ PARTI records drinking activity and bottle preference with remarkably high accuracy, as measured through a correlation of LIQ activity and manually weighed bottle measurements (**Supplemental Figure 2**). With this system, we observed that baseline measurements of water consumption between male and female group-housed mice are similar, except for a subtle, but significant increase in lick duration and frequency in female mice compared to males (**Supplemental Figure 3**).

With the use of LIQ PARTI in a continuous access two-bottle choice ethanol paradigm, we show that male and female group-housed mice develop different preferences for ethanol and modulate their bout microstructure in contrasting ways. As previously reported throughout the literature, female C57Bl/6J mice develop a significantly higher preference for ethanol than males (**Figure 1**) (M. A. Caruso et al., 2021; Centanni et al., 2019a; Flores-Bonilla & Richardson, 2020; Lopez & Laber, 2015; Middaugh et al., 1999; Rivera-Irizarry et al., 2023; Sneddon et al., 2019; Yu et al., 2019). However, with the advantages of the high temporal resolution of LIQ PARTI, we revealed that male mice display a unique pattern of ethanol consumption/preference. Male mice show a repetitive, undulating, within-week pattern of ethanol preference with the highest value occurring in concert with cage changes that diminishes over 24-48 hours. While female mice have a persistently elevated ethanol preference each week, male mice begin each recording period with a preference for ethanol at similar levels to the female mice but dip to a very low preference by the end of the recording period. Given the length of time the male mice take to diminish their ethanol preference, the sensory cues for 10% ethanol, and the absence of this pattern in females, we believe this phenomenon is unlikely to be due simply to the swapping of bottle placement. This wave-like pattern of ethanol preference in males may represent a sensitivity to cage-change stress as stress has been shown to increase ethanol consumption and preference in mice (Anderson et al., 2016; Bahi, 2013; M. J. Caruso et al., 2018; Reguilón et al., 2020). These issues will need to be further interrogated in future studies. Furthermore, bout microstructure changes in response to prolonged ethanol exposure are significantly different between male and female mice (**Figure 2**). We previously have shown that female mice display a decrease in average bout size and bout duration at both the water and ethanol bottles during chronic ethanol exposure when singly housed with the LIQ HD system (Petersen et al., 2023). We observed this same pattern in group-housed female mice, but male mice showed the opposite effect, with an increase in bout size and duration at both bottles compared to the water-only week (**Figure 2B,C**). We also showed previously in singly housed female mice that lick frequency increased and ILI decreased at the ethanol bottle compared to the water bottle during chronic ethanol exposure (Petersen et al., 2023). Interestingly, group-housed female mice displayed an increased frequency at the water bottle compared to males and compared to the water-only week, and increased ILI at the ethanol bottle compared to the water bottle and compared to the water-only week. Males only showed a decrease in lick frequency at the water bottle compared to the water-only week (**Figure 2E,F**). Together this suggests that social enrichment during the acquisition of ethanol preference and sex alters how bout microstructure is affected following chronic ethanol exposure.

Social isolation prior to ethanol exposure has been shown to significantly increase ethanol intake in mice (Butler, Ariwodola, et al., 2014; Cortés-Patiño et al., 2016; Cullins & Chester, 2024; Evans et al., 2020; Lopez et al., 2011; Lopez & Laber, 2015; Panksepp et al., 2017; Sanna et al., 2011), and here we show that social isolation in ethanol-experienced male and female mice increased ethanol preference, total fluid intake, and altered bout microstructure. We utilized LIQ HD to investigate ethanol drinking behavior during social isolation. It is important to note that LIQ PARTI and LIQ HD differ slightly in their physical structure, but both systems use identical sippers in the same vertical orientation, and the LIQ HD source code was modified to include the same lick detection optimization and bout logging method as LIQ PARTI. Furthermore, we validated that mice on average consume the same amount of liquid (∼1.4µL) per lick in both systems (**Supplemental Figure 4**). Both male and female mice significantly increased their total lick number (**Figure 3D; Figure 5E**), fluid consumption (**Figure 5F**), and ethanol preference (**Figure 3E**, **Figure 5G,H**) during social isolation. Interestingly, male mice no longer displayed the undulating pattern of ethanol preference that corresponded to cage-changing days, but instead showed persistently elevated preference throughout the recording periods similar to female mice (**Figure 3C**). This may be due to the stress of social isolation overwhelming the stress induced by cage changes. Moreover, male mice have a more rigid social dominance hierarchy than females (Karamihalev et al., 2020), and group-housed males may experience additional stress during cage changes due to the re-establishment of the social dominance hierarchy (Gray & Hurst, 1995; Hurst, 1993; Lidster et al., 2019), which would not be a factor during social isolation. In addition to male mice displaying elevated ethanol preference during social isolation at comparable levels to females in the group-housed environment, bout microstructure of the male mice, specifically bout size and duration, during social isolation closely resembled the bout microstructure of the female mice (**Figure 4**). Both male and female mice had an increased average lick duration at both bottles during social isolation, with a greater lick duration at the water bottle compared to the ethanol bottle. Interestingly, the lick frequency in female mice between the two bottles flipped during social isolation, with a greater lick frequency at the ethanol bottle compared to the water bottle, which recapitulates our previous study using LIQ HD in female mice (Petersen et al., 2023).

Several factors have been shown to contribute to changes in ethanol drinking microstructure. The literature suggests that bout size and lick frequency are directly related to the palatability of a drinking solution (Baird et al., 2005; Davis & Levine, 1977; Dwyer, 2012; Dwyer et al., 2011; Johnson, 2018).

However, although bout size and duration during social isolation are greater at the ethanol bottle compared to the water bottle, our data suggest that a decreased bout size and duration overall are associated with a significantly elevated ethanol preference. In contrast, lick frequency in female mice is significantly increased by social isolation at the ethanol bottle. Additionally, changes in lick microstructure may represent different behavioral strategies for consumption. For example, broadly, the rate of food or fluid intake has been suggested as an indicator of motivation (Johnson, 2018; Radke et al., 2021). Behavioral strategies in the context of ethanol intake have been particularly characterized in the context of quinine-adulterated ethanol intake in rodents, a model of compulsive ethanol intake. For example, Darevsky et al. describe reduced variability in lick measures, interpreted as greater automatic responding (Darevsky et al., 2019). Finally, circulating ovarian hormones may also factor into drinking microstructure. While previous studies have shown no change in ethanol consumption across the estrous cycle in female rodents (De Oliveira Sergio et al., 2024; Ford et al., 2002; Satta et al., 2018; for a summary of studies see Radke et al., 2021), estrous phase can influence ethanol bout frequency and size (Ford et al., 2002), suggesting that sex differences in microstructure may be in part driven by hormonal factors. The estrous cycle was not tracked in the current study to avoid disruptions in continuous drinking behavior; however, this represents a future direction of interest. LIQ PARTI represents an ideal tool for use in studying the implications of these factors on group-housed ethanol drinking behavior.

As mentioned previously, prior experience of social isolation has significant and persistent impacts on ethanol preference in mice (Butler, Ariwodola, et al., 2014; Cortés-Patiño et al., 2016; Cullins & Chester, 2024; Evans et al., 2020; Lopez et al., 2011; Lopez & Laber, 2015; Panksepp et al., 2017; Sanna et al., 2011). Holgate et al. (2017) found that male C57Bl/6J mice with previous ethanol exposure showed an increased preference for ethanol in a drinking in the dark (DID) task when moved from an enriched environment to social isolation but were unable to assess resocialization with two-bottle choice in the home cage due to aggressive behavior. Fortunately, we did not observe aggressive behavior following resocialization with the same cage mates. We found that fluid consumption, ethanol preference, and bout microstructure mostly returned to pre-social isolation values when the mice were reintroduced to group housing with LIQ PARTI (**Figures 5-7; Supplemental Figure 5**). During the first group-housed week post-social isolation, male mice had a persistently elevated average lick number per day and fluid consumption compared to pre-social isolation but female mice returned to pre-social isolation values (**Figure 5E,F**; **Figure 6D**). Average total lick number per day and total fluid consumption modestly decreased in females during the second week of post-social isolation and males returned to pre-social isolation levels. In contrast, ethanol preference in male mice quickly returned to pre-social isolation levels with the undulating pattern by the first week, but females had a significantly reduced preference during the first week of post-social isolation and returned to pre-social isolation values by the second week (**Figure 5G,H**; **Figure 6C,E**). Furthermore, bout microstructure in both male and female mice reverted to pre-social isolation value with some minor exceptions (**Figure 7; Supplemental Figure 5**). For example, male mice showed a decreased bout size and bout duration at the ethanol bottle and female mice had a slightly increased lick duration at both bottles and decreased lick frequency at the water bottle post-social isolation compared to pre-social isolation, but to a much lower degree than during social isolation. Thus, social isolation-induced changes to ethanol drinking behavior and bout microstructure in ethanol-experienced mice are highly plastic, largely reverting to the pre-social isolation baseline within two weeks.

Loneliness in humans has historically been associated with contributing to and maintaining alcohol use and AUD, however while those with AUD have been found to have heightened feelings of loneliness, the relationship between social interaction and alcohol use is unclear (Åkerlind & Hörnquist, 1992; Mowbray et al., 2014). The COVID-19 pandemic and lockdown period was an era of intense social isolation for many individuals worldwide. While there was an increase in alcohol sales per capita in the US during the pandemic, there was a significant decrease in the average drinks consumed per day with no difference in drinking days per month, binge drinking days per month, or maximum drinks per day (Castaldelli-Maia et al., 2021; Moskatel & Slusky, 2023). Furthermore, alcohol sales and consumption by all metrics decreased in the US during periods with a high incidence of COVID-19 (Moskatel & Slusky, 2023). Studies in Brazil (Moura et al., 2023), Germany (Deeken et al., 2022), Australia (Bade et al., 2021), and others (Acuff et al., 2022), similarly found that alcohol consumption was in general significantly lower during lockdown periods. However, increased alcohol consumption was significantly associated with both particularly high and particularly low degrees of subjective loneliness (Bragard et al., 2022; Haucke et al., 2024). Additionally, online engagement in an AUD recovery service was significantly decreased, and alcohol withdrawal hospitalizations and alcohol-related deaths significantly increased in the US during the COVID-19 pandemic (Colditz et al., 2021; Sharma et al., 2021; White et al., 2022). Alcohol use, AUD, social isolation, and loneliness thus represent a complex interaction that requires further investigation with significant public health implications. Novel tools, such as LIQ PARTI and LIQ HD, will aid in the investigation of how socialization and social isolation affect consummatory behaviors and the underlying physiology in rodent models.

In conclusion, our study demonstrates the utility of LIQ PARTI and LIQ HD in elucidating sex differences and the impact of social isolation on ethanol consumption and bout microstructure in C57Bl/6J mice. We observed distinct patterns of ethanol preference between male and female mice, with males exhibiting an intriguing undulating response to cage changes. Moreover, chronic ethanol exposure induced significant alterations in bout microstructure, influenced by both sex and social environment.

Social isolation in the ethanol-experience mice led to marked increases in ethanol intake and preference in both sexes, underscoring the role of social context in modulating ethanol drinking behavior. However, the transition back to group housing resulted in the reversal of social isolation-induced changes to ethanol drinking behaviors, highlighting the plasticity of these responses. Overall, our findings contribute to a deeper understanding of the complex interplay between sex, social environments, and ethanol consumption, with future implications for understanding the physiological mechanisms underlying the relationship between loneliness and alcohol use.

## Supplemental Figures

**Supplemental Figure 1.**
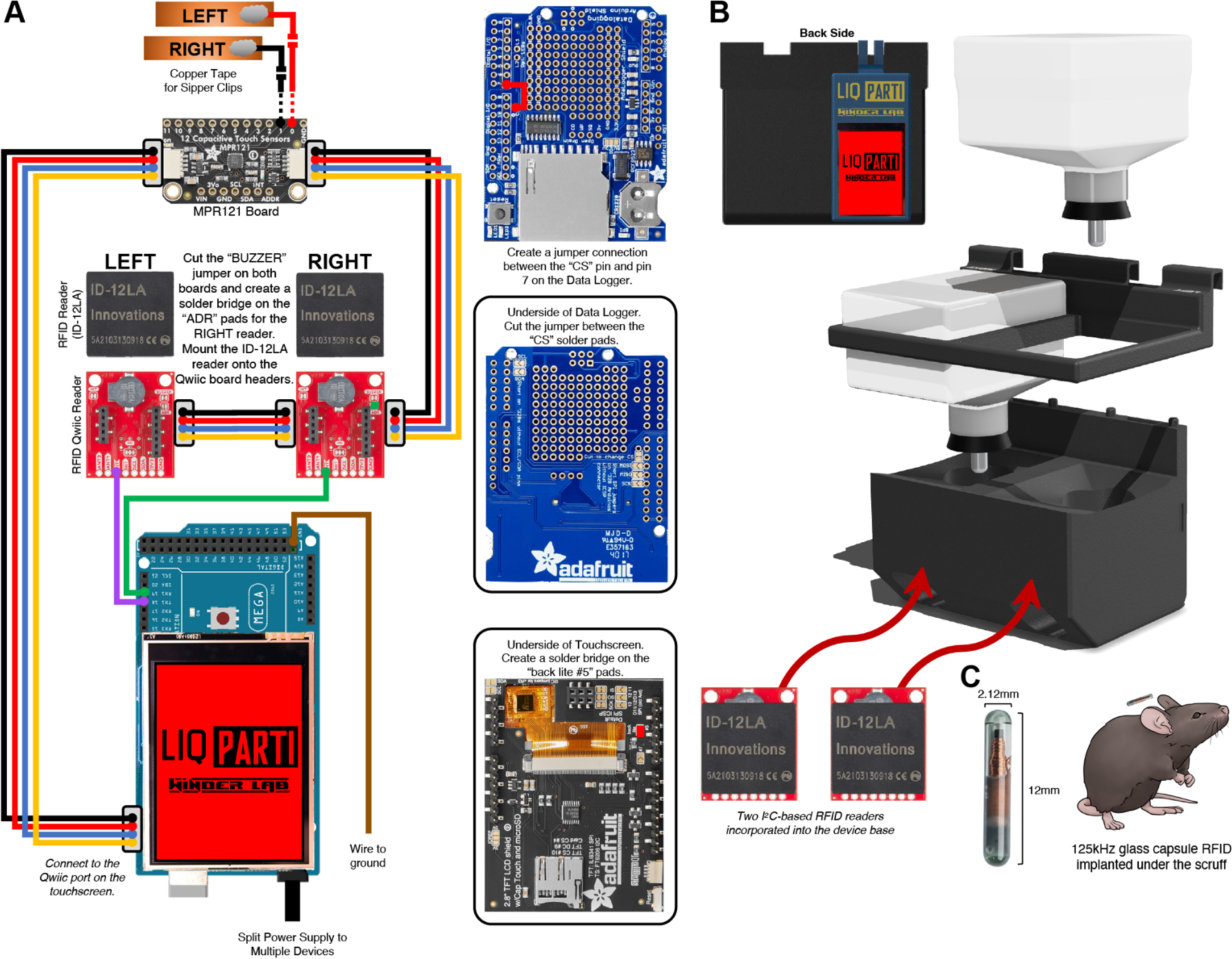
LIQ PARTI wiring diagram and build. **A**, LIQ PARTI electronic parts and wiring diagram including an Arduino Mega, two RFID readers, an MPR121 capacitive touch board, a touchscreen shield, a data logger shield, and conductive copper foil tape. **B**, 3D rendering of LIQ PARTI disassembled components, including 3D-printed parts, rubber stoppers with sippers, RFID readers, and Arduino microcontroller. **C**, Glass capsule RFID tag with dimensions and representative illustration of the RFID tag to scale (estimated) with a mouse. More detailed information can be found at (https://github.com/nickpetersen93/LIQ_PARTI).

**Supplemental Figure 2.**
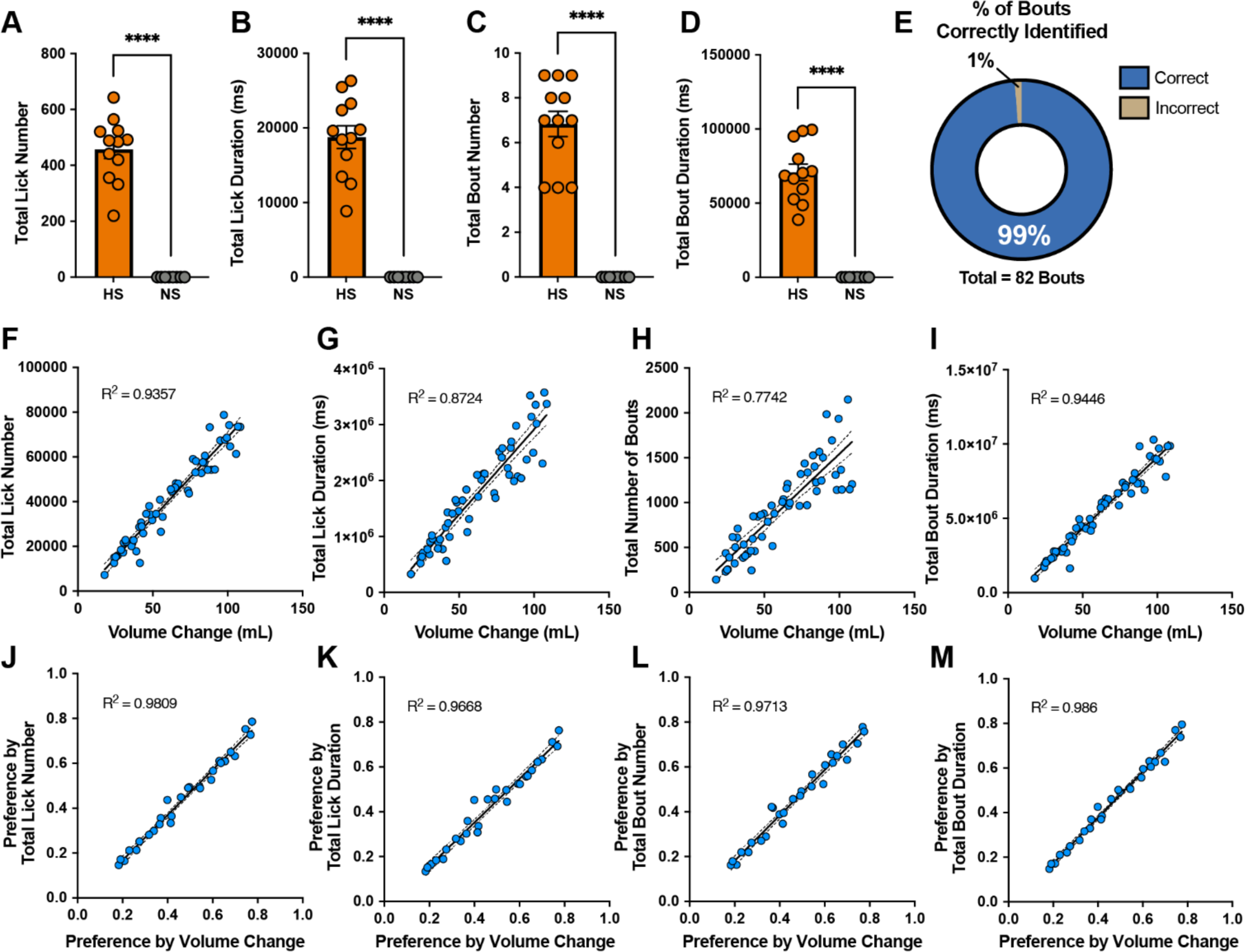
LIQ PARTI accurately identifies individual mice in a group-housed environment and lick and bout detection closely correlates with bottle weight measurements. Total lick number (**A**), lick duration (**B**), bout number (**C**), and bout duration (**D**) of mice injected with hypertonic saline (HS) or normal saline (NS) during the LIQ PARTI video recording of water drinking for the validation experiment (unpaired t-test). (*n* = 12 mice HS group; *n* = 8 mice NS group). **E**, The percentage of video-recorded drinking bouts that were attributed by LIQ PARTI to the correct RFID-tagged mouse. Correlation between whole-cage total lick number and volume change (**F**), total lick duration and volume change (**G**), total bout number and volume change (**H**), and total bout duration and volume change (**I**) for each recording period. Correlation between preference calculated by total lick number and preference calculated by volume change (**J**), preference by total lick duration and preference by volume change (**K**), preference by total bout number and preference by volume change (**L**), and preference by total bout duration and preference by volume change (**M**) for each recording period. In correlation graphs, solid lines represent a fitted simple linear regression model, and dashed lines denote 95% confidence intervals. Data in A-E are reported as individual mouse values and F-M are reported as whole-cage values. *\*\*\*\*p* < 0.0001.

**Supplemental Figure 3.**
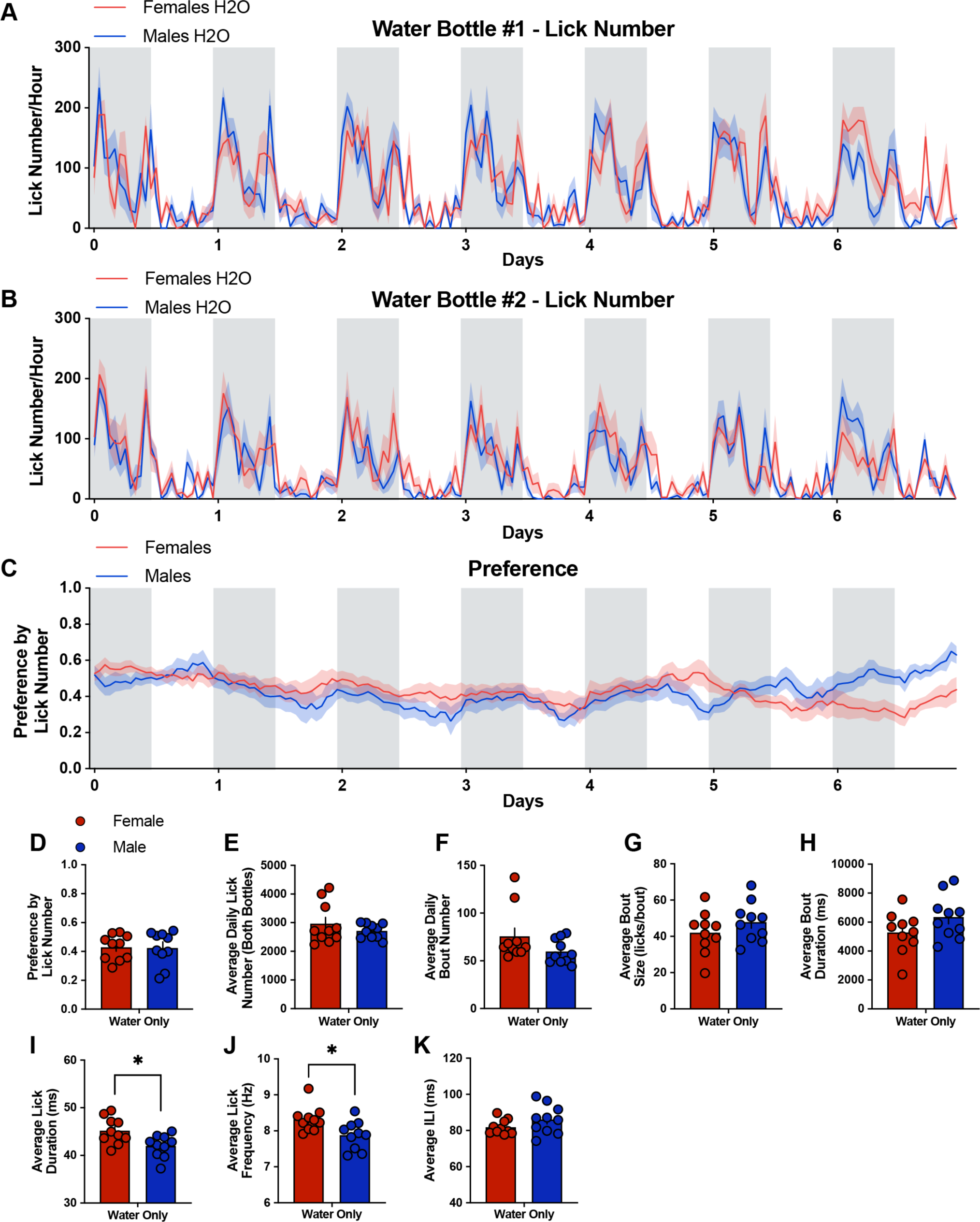
Group-housed male and female mice consume water at similar rates when given continuous access to two water bottles, and female mice show a significantly higher average lick frequency and lick duration compared to males at baseline. Lick number per hour from male and female mice at the left water bottle (**A**) and the right water bottle (**B**) throughout the water-only week with the group-housed LIQ PARTI system. **C**, Bottle preference score calculated by lick number at each water bottle in male and female mice throughout the water-only week. The average preference by lick number (**D**), average lick number per day (**E**), average daily bout number (**F**), average bout size (**G**), average bout duration (**H**), average lick duration (**I**), average lick frequency (**J**), and average ILI (**K**) were calculated based on total values at both bottles and compared between males and females during the water-only week with LIQ PARTI. The solid lines (**A-C**) represent the mean lick number in 1-h bins, and the red/blue shaded areas represent ±SEM. Gray shaded areas represent the dark cycle. Error bars (**D-K**) represent ±SEM. (*n* = 10 mice per group and reported as individual mice). *\*p* < 0.05.

**Supplemental Figure 4.**
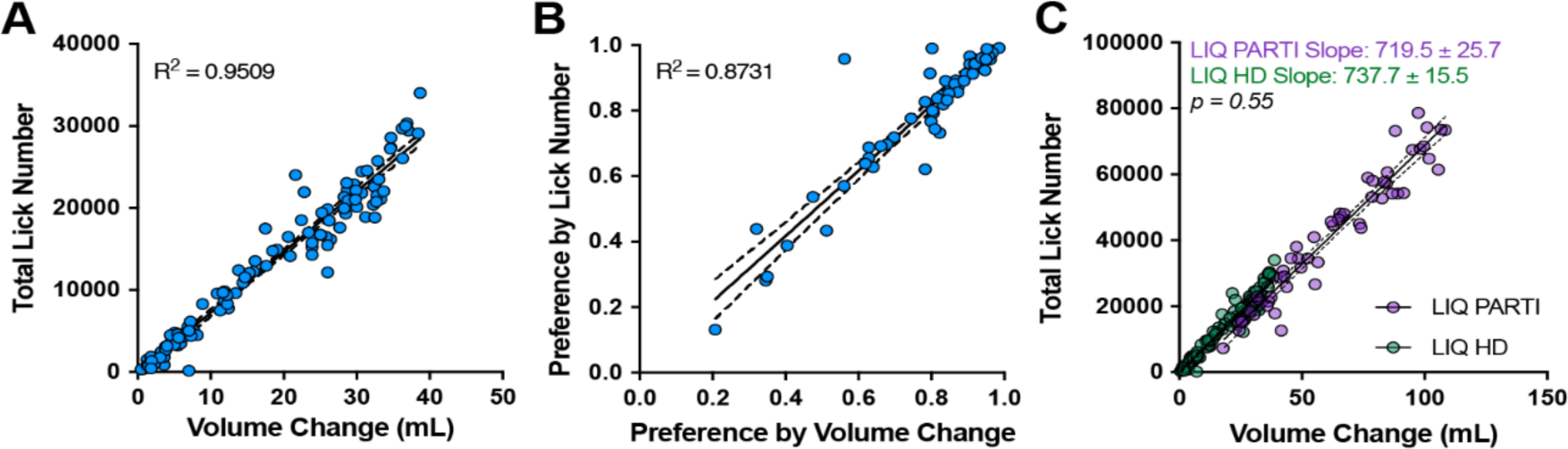
LIQ HD and LIQ HD vs LIQ PARTI validation. Correlation between total lick number and volume change (**A**) and correlation between preference calculated total lick number and preference by volume change (**B**) for each recording period of social isolation with LIQ HD. **C**, comparison of correlations between total lick number and volume change for LIQ HD and LIQ PARTI. There is no significant difference between the fitted region slopes for the LIQ HD and LIQ PARTI systems. Solid lines represent a fitted simple linear regression model, and dashed lines denote 95% confidence intervals. LIQ PARTI data are reported as whole-cage values.

**Supplemental Figure 5.**
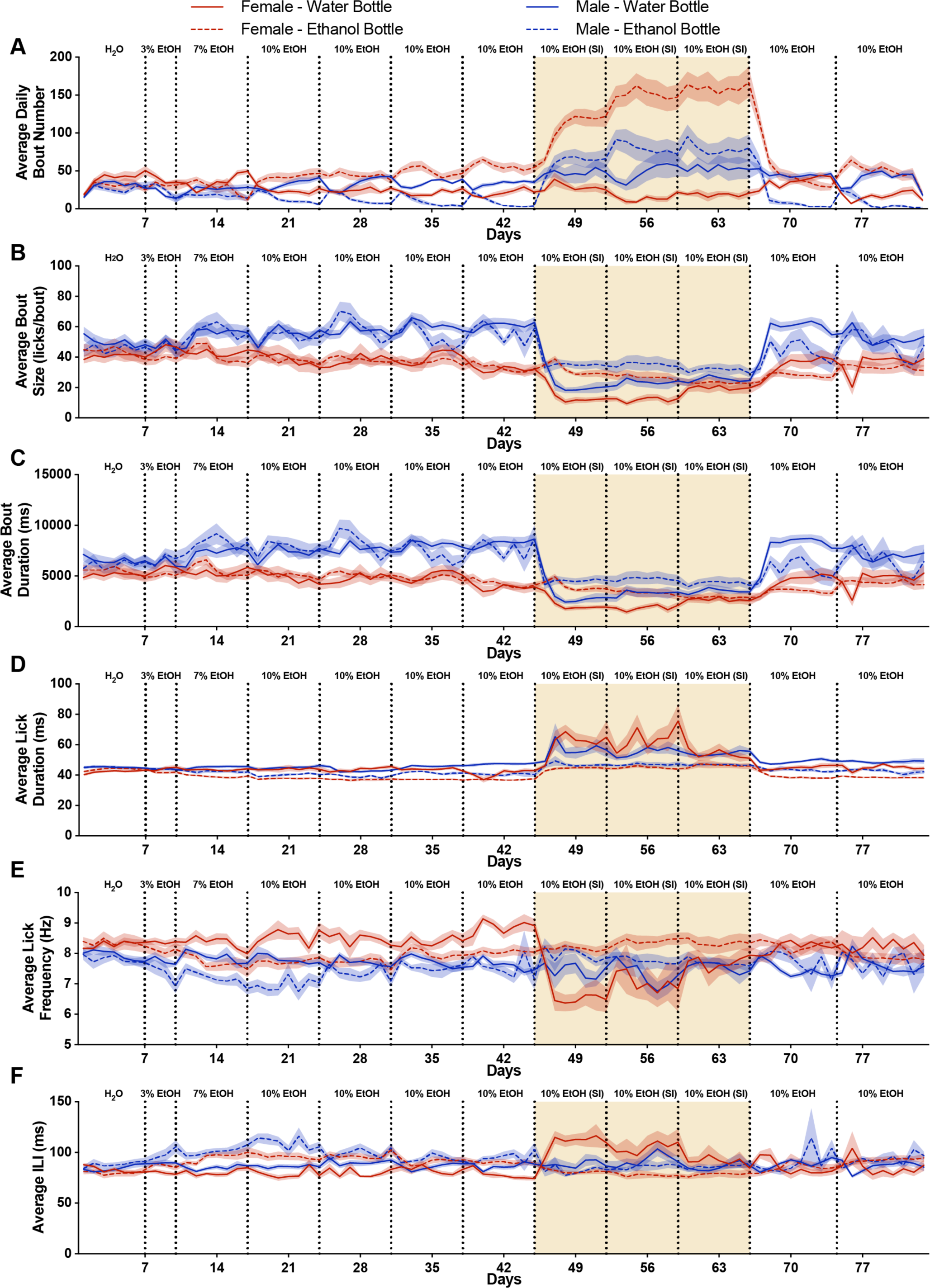
Exposure to ethanol and social isolation significantly alters drinking microstructure in a sex-, bottle, and housing-specific manner over time. Average daily bout number (**A**), bout size (**B**), bout duration (**C**), lick duration (**D**), lick frequency (**E**), and ILI (**F**) for male and female mice at the water and ethanol bottles. Lines represent the means in 24-h bins, and the red/blue shaded areas represent ±SEM. Yellow shaded regions indicate periods of social isolation with LIQ HD and vertical dotted lines represent when cages were cleaned and bottle positions were swapped. (*n* = 10 mice per group, except males post-SI week 2 n = 5, and reported as individual mice).

**Supplemental Video 1.** Manually annotated representative video recording (shown here at 2x speed) for LIQ PARTI RFID-tracking validation. Mice, same-sex and five per cage, were implanted with a glass capsule RFID tag subcutaneously in the scruff of the neck and their tails were marked with different colors for video tracking. In this recording, three of the mice were injected with hypertonic saline, to induce thirst, and two mice were injected with normal saline 15 minutes before starting the recording. The mice were then placed in a clean cage with the LIQ PARTI system containing water in both bottles and allowed to explore the cage and consume water from either bottle for 10 minutes. The annotated circle icons indicate when an RFID tag was scanned, the water drop icons indicate when a drinking bout was identified, and the floppy disk save icon indicates when a drinking bout was saved to the SD card. The icon color matches the color of the mouse’s tail that corresponds to that mouse’s RFID number. The Arduino IDE serial monitor is displayed at the bottom of the video and displays real-time information.

**Extended Data 1:** LIQ PARTI Arduino source code used in this study.

