## Extended Data 1 for "A novel mouse home cage lickometer system reveals sex- and housing-based influences on alcohol drinking": LIQ_functions.ino

```
//functions used throughout code

//======================================================================================

void reset_total_LN() {
  for (int k = 0; k < sipper_number; k++) {
    total_LN[k] = 0;  //reset totals
  }
}

//======================================================================================

void reset_variables() {
  //Reset lick variables
  for (int k = 0; k < sipper_number; k++) {
    LickNumber[k] = 0;
    LickDuration[k] = 0;
    BoutNumber[k] = 0;
    BoutDuration[k] = 0;
    BoutLickNumber[k] = 0;
    BoutLickDuration[k] = 0;
    startmillis = millis();
    tag1 = "";
    tag2 = "";
    lasttag1 = "";
    lasttag2 = "";
    tag1_count = 0;
    lasttag1_count = 0;
    tag2_count = 0;
    lasttag2_count = 0;
    notag1_count = 0;
    notag2_count = 0;
    RFIDerrorLEFT = false;
    RFIDerrorLEFT = false;
    RFIDerrorLEFT = false;
    RFIDerrorLEFT = false;
  }
}

//==========================================================================================

void Record_Licks() {
  for (int k = 0; k < 2; k++) {  //for cap sensor 1
    currtouched1 = cap.touched();

    if ((currtouched1 & _BV(k)) && !licking[k]) {  //sensor is touched
      DEBUG_MPR121_PRINT(k);
      DEBUG_MPR121_PRINTLN(" touched");
      cal_timer = millis();
      time_now[k] = millis();
      licking[k] = true;
      LickNumber[k] += 1;
      BoutLickNumber[k] += 1;
      lick_bout_countdown[k] += 1;
      if (lick_bout_countdown[k] == 1) {
        bout_start_timer[k] = millis();
      }
      if (lick_bout_countdown[k] == 3 && millis() - bout_start_timer[k] <= 1000) {  //bout has started if more than 3 licks in less than 1 second
        BoutNumber[k] += 1;
        bout_timer[k] = millis();
        in_bout[k] = true;
        if (k == 0) {
          DEBUG_LIQ_PRINT("LEFT");
          DEBUG_LIQ_PRINTLN(" bout started!");
        }
        if (k == 1) {
          DEBUG_LIQ_PRINT("RIGHT");
          DEBUG_LIQ_PRINTLN(" bout started!");
        }
      }
      if (in_bout[k]) {
        bout_timer[k] = millis();
      }
    }

    if (!(currtouched1 & _BV(k)) && licking[k]) {  //sensor is released
      DEBUG_MPR121_PRINT(k);
      DEBUG_MPR121_PRINTLN(" released");
      Elapsedtime[k] = millis() - time_now[k];
      LickDuration[k] = LickDuration[k] + Elapsedtime[k];
      licking[k] = false;
      BoutLickDuration[k] = BoutLickDuration[k] + Elapsedtime[k];
      last_lick_time[k] = millis();
      DEBUG_MPR121_PRINTLN(LickNumber[k]);
      DEBUG_MPR121_PRINTLN(LickDuration[k]);
    }
  }
  lasttouched1 = currtouched1;

  for (int k = 0; k < 2; k++) {
    if (lick_bout_countdown[k] <= 2 && millis() - bout_start_timer[k] >= 1000) {
      bout_start_timer[k] = 0;
      lick_bout_countdown[k] = 0;
      BoutLickNumber[k] = 0;
      BoutLickDuration[k] = 0;
    }

    if (lick_bout_countdown[k] >= 3 && millis() - bout_timer[k] >= bout_cutoff) {
      if (k == 0) {
        DEBUG_LIQ_PRINT("LEFT");
        DEBUG_LIQ_PRINTLN(" bout ended!");
      }
      if (k == 1) {
        DEBUG_LIQ_PRINT("RIGHT");
        DEBUG_LIQ_PRINTLN(" bout ended!");
      }
      in_bout[k] = false;
      BoutDuration[k] = last_lick_time[k] - bout_start_timer[k];
      LickFrequency[k] = BoutLickNumber[k] / (BoutDuration[k] / 1000);
      ILI[k] = (BoutDuration[k] - BoutLickDuration[k]) / BoutLickNumber[k];
      write_to_file(k);

      bout_start_timer[k] = 0;
      lick_bout_countdown[k] = 0;
      BoutDuration[k] = 0;
      BoutLickNumber[k] = 0;
      BoutLickDuration[k] = 0;
    }
  }
}


//======================================================================================

void update_sippers() {
  for (int k = 0; k < 2; k++) {
    if (licking[k]) {
      Elapsedtime[k] = millis() - time_now[k];
      LickDuration[k] = LickDuration[k] + Elapsedtime[k];
      if (in_bout[k]) {
        BoutDuration[k] = BoutDuration[k] + (millis() - bout_start_timer[k]);
        BoutLickDuration[k] = BoutLickDuration[k] + Elapsedtime[k];
      }
    }
  }
}

//======================================================================================

void reset_time_now() {
  for (int k = 0; k < 2; k++) {
    time_now[k] = millis();  //reset time now
    bout_start_timer[k] = millis();
  }
}

//======================================================================================

void calc_total_LN() {
  for (int k = 0; k < 2; k++) {
    total_LN[k] = total_LN[k] + LickNumber[k];  //calculate total LN for each sipper
  }
}

//======================================================================================

void wait(unsigned long ms) {
  unsigned long X = millis();
  while (millis() < X + ms) {}  //wait without delay for ms (milliseconds)
}

//======================================================================================

void set_defaults() {
  default_lights_on = lights_on;
  default_lights_off = lights_off;
  default_touch_threshold = touch_threshold;
  default_release_threshold = release_threshold;
  default_LOG_INTERVAL = LOG_INTERVAL;
  dafault_SYNC_INTERVAL = SYNC_INTERVAL;
  default_auto_cal = auto_cal;
  default_auto_cal_time = auto_cal_time;
  default_auto_cal_sec_since_last_lick = auto_cal_sec_since_last_lick;
}

//======================================================================================

void auto_calibration() {
  if (auto_cal) {
    unsigned long temp_cal_time = 1000 * auto_cal_sec_since_last_lick;
    unsigned long temp_cal_timer = millis() - cal_timer;
    now = rtc.now();
    if (auto_cal_flag && now.hour() == auto_cal_time && temp_cal_timer > temp_cal_time) {
      auto_cal_flag = false;
      start_rfid();
      cap.begin(0x5A);
      set_sensor_settings();
    }
    if (now.hour() != auto_cal_time) {
      auto_cal_flag = true;
    }
  }
}

//======================================================================================

void set_sensor_settings() {
  cap.setThresholds(touch_threshold, release_threshold);  //set sensitivity of touch and release of capacitive sensors
  cap.writeRegister(MPR121_CONFIG1, 0);                   // default, 16uA charge current
  cap.writeRegister(0x5F, current);
  cap.writeRegister(0x60, current);
  //        cap.writeRegister(0x61, current);
  //        cap.writeRegister(0x62, current);
  //        cap.writeRegister(0x63, current);
  //        cap.writeRegister(0x64, current);
  //        cap.writeRegister(0x65, current);
  //        cap.writeRegister(0x66, current);
  //        cap.writeRegister(0x67, current);
  //        cap.writeRegister(0x68, current);
  //        cap.writeRegister(0x69, current);
  //        cap.writeRegister(0x6A, current);
  cap.writeRegister(MPR121_DEBOUNCE, debounce);
  cap.writeRegister(MPR121_MHDR, 1);
  cap.writeRegister(MPR121_NHDR, 10);
  cap.writeRegister(MPR121_NCLR, 1);
  cap.writeRegister(MPR121_FDLR, 2);

  cap.writeRegister(MPR121_MHDF, 1);
  cap.writeRegister(MPR121_NHDF, 5);
  cap.writeRegister(MPR121_NCLF, 3);
  cap.writeRegister(MPR121_FDLF, 1);

  cap.writeRegister(MPR121_NHDT, 1);
  cap.writeRegister(MPR121_NCLT, 5);
  cap.writeRegister(MPR121_FDLT, 1);
}
```
