## Extended Data 1 for "A novel mouse home cage lickometer system reveals sex- and housing-based influences on alcohol drinking": LIQ_PARTI_Arduino_Mega.ino

```
//#define DEBUG_LIQ //uncomment to turn on debug mode, prints info to Serial Monitor
//#define DEBUG_RFID //uncomment to turn on debug mode, prints info to Serial Monitor
//#define DEBUG_MPR121 //uncomment to turn on debug mode, prints info to Serial Monitor
//#define DEBUG_Screen //uncomment to turn on debug mode, prints info to Serial Monitor
//#define DEBUG_System //uncomment to turn on debug mode, prints info to Serial Monitor

/********************************************************
  LIQ PARTI
  Written by Nicholas Petersen
  https://github.com/nickpetersen93/LIQ_PARTI

  January, 2023

  This work includes libraries with the following licensing:
  Adafruit GFX Library - Written by Limor Fried/Ladyada for Adafruit Industries. BSD License
  Adafruit ILI9341 Library - Written by Limor Fried/Ladyada for Adafruit Industries. MIT license. BSD License.
  Adafruit FT6206 Library - Written by Limor Fried/Ladyada for Adafruit Industries. MIT license.
  Adafruit MPR121 Library - Written by Limor Fried/Ladyada for Adafruit Industries. MIT license. BSD License.
  SparkFun Qwiic RFID Library  - Written by SparkFun. MIT license.
  RTCLib Library - Written by JeeLabs. MIT license.

  This work is released under This work is licensed under a Creative Commons Attribution-NonCommercial-ShareAlike 4.0 International License:
  https://creativecommons.org/licenses/by-nc-sa/4.0/legalcode

  Copyright (c) 2023 Nicholas Petersen. Creative Commons BY-NC-SA License. Non-Commercial Use Only.
  Adapter’s License must be a Creative Commons BY-NC-SA or Compatible License.
  All text above must be included in any redistribution. See LICENSE.md for detailed information.

  UNLESS OTHERWISE SEPARATELY UNDERTAKEN BY THE LICENSOR, TO THE EXTENT POSSIBLE, THE LICENSOR OFFERS
  THE LICENSED MATERIAL AS-IS AND AS-AVAILABLE, AND MAKES NO REPRESENTATIONS OR WARRANTIES OF ANY KIND
  CONCERNING THE LICENSED MATERIAL, WHETHER EXPRESS, IMPLIED, STATUTORY, OR OTHER. THIS INCLUDES,
  WITHOUT LIMITATION, WARRANTIES OF TITLE, MERCHANTABILITY, FITNESS FOR A PARTICULAR PURPOSE,
  NON-INFRINGEMENT, ABSENCE OF LATENT OR OTHER DEFECTS, ACCURACY, OR THE PRESENCE OR ABSENCE OF ERRORS,
  WHETHER OR NOT KNOWN OR DISCOVERABLE. WHERE DISCLAIMERS OF WARRANTIES ARE NOT ALLOWED IN FULL OR IN
  PART, THIS DISCLAIMER MAY NOT APPLY TO YOU.

  TO THE EXTENT POSSIBLE, IN NO EVENT WILL THE LICENSOR BE LIABLE TO YOU ON ANY LEGAL THEORY (INCLUDING,
  WITHOUT LIMITATION, NEGLIGENCE) OR OTHERWISE FOR ANY DIRECT, SPECIAL, INDIRECT, INCIDENTAL,
  CONSEQUENTIAL, PUNITIVE, EXEMPLARY, OR OTHER LOSSES, COSTS, EXPENSES, OR DAMAGES ARISING OUT OF THIS
  PUBLIC LICENSE OR USE OF THE LICENSED MATERIAL, EVEN IF THE LICENSOR HAS BEEN ADVISED OF THE
  POSSIBILITY OF SUCH LOSSES, COSTS, EXPENSES, OR DAMAGES. WHERE A LIMITATION OF LIABILITY IS NOT
  ALLOWED IN FULL OR IN PART, THIS LIMITATION MAY NOT APPLY TO YOU.

  THE DISCLAIMER OF WARRANTIES AND LIMITATION OF LIABILITY PROVIDED ABOVE SHALL BE INTERPRETED IN A
  MANNER THAT, TO THE EXTENT POSSIBLE, MOST CLOSELY APPROXIMATES AN ABSOLUTE DISCLAIMER AND WAIVER OF
  ALL LIABILITY.


********************************************************/

#include <Adafruit_GFX.h>
#include <SPI.h>
#include <Wire.h>
#include <SD.h>
#include "RTClib.h"
#include "Adafruit_MPR121.h"
#include "SparkFun_Qwiic_Rfid.h"
#include <Adafruit_ILI9341.h>
#include <Adafruit_FT6206.h>
#include <math.h>
#include <Fonts/FreeSans9pt7b.h>
#include <Fonts/FreeSans12pt7b.h>
#include <Fonts/FreeSans18pt7b.h>
#include <Fonts/FreeSansBold9pt7b.h>
#include <Fonts/FreeSansBold12pt7b.h>
#include <Fonts/FreeSansBold18pt7b.h>
#include <Fonts/FreeSansBold24pt7b.h>

#ifdef DEBUG_LIQ
#define DEBUG_LIQ_PRINT(x)     Serial.print (x)
#define DEBUG_LIQ_PRINTDEC(x)     Serial.print (x, DEC)
#define DEBUG_LIQ_PRINTLN(x)  Serial.println (x)
#else
#define DEBUG_LIQ_PRINT(x)
#define DEBUG_LIQ_PRINTDEC(x)
#define DEBUG_LIQ_PRINTLN(x)
#endif

#ifdef DEBUG_RFID
#define DEBUG_RFID_PRINT(x)     Serial.print (x)
#define DEBUG_RFID_PRINTDEC(x)     Serial.print (x, DEC)
#define DEBUG_RFID_PRINTLN(x)  Serial.println (x)
#else
#define DEBUG_RFID_PRINT(x)
#define DEBUG_RFID_PRINTDEC(x)
#define DEBUG_RFID_PRINTLN(x)
#endif

#ifdef DEBUG_MPR121
#define DEBUG_MPR121_PRINT(x)     Serial.print (x)
#define DEBUG_MPR121_PRINTDEC(x)     Serial.print (x, DEC)
#define DEBUG_MPR121_PRINTLN(x)  Serial.println (x)
#else
#define DEBUG_MPR121_PRINT(x)
#define DEBUG_MPR121_PRINTDEC(x)
#define DEBUG_MPR121_PRINTLN(x)
#endif

#ifdef DEBUG_Screen
#define DEBUG_Screen_PRINT(x)     Serial.print (x)
#define DEBUG_Screen_PRINTDEC(x)     Serial.print (x, DEC)
#define DEBUG_Screen_PRINTLN(x)  Serial.println (x)
#else
#define DEBUG_Screen_PRINT(x)
#define DEBUG_Screen_PRINTDEC(x)
#define DEBUG_Screen_PRINTLN(x)
#endif

#ifdef DEBUG_System
#define DEBUG_System_PRINT(x)     Serial.print (x)
#define DEBUG_System_PRINTDEC(x)     Serial.print (x, DEC)
#define DEBUG_System_PRINTLN(x)  Serial.println (x)
#else
#define DEBUG_System_PRINT(x)
#define DEBUG_System_PRINTDEC(x)
#define DEBUG_System_PRINTLN(x)
#endif


// Default RFID I2C addresses
#define RFID_ADDR1 0x7D
#define RFID_ADDR2 0x7C

Qwiic_Rfid myRfid1(RFID_ADDR1);
Qwiic_Rfid myRfid2(RFID_ADDR2);


// The FT6206 uses hardware I2C on the shield
Adafruit_FT6206 ts = Adafruit_FT6206();

// The display also uses hardware SPI, plus #9 & #10
#define TFT_CS 10
#define TFT_DC 9
Adafruit_ILI9341 tft = Adafruit_ILI9341(TFT_CS, TFT_DC);

RTC_PCF8523 rtc; // define the Real Time Clock object
const int chipSelect = 7; // for the data logging shield, changed from digital pin 10
#define redLEDpin 13
DateTime now;

//capacitive touch device
#ifndef _BV
#define _BV(bit) (1 << (bit))
#endif

// You can have up to 4 on one i2c bus but one is enough for testing!
Adafruit_MPR121 cap = Adafruit_MPR121();

// Keeps track of the last pins touched so we know when buttons are 'released'
uint16_t lasttouched1 = 0;
uint16_t currtouched1 = 0;

File logfile; // the logging file
char filename[] = "00000000.CSV";

TS_Point p;
extern uint8_t welcome_logo[];
extern uint8_t main_logo[];
extern uint8_t settings_icon[];
extern uint8_t settings_icon2[];
extern uint8_t PARTI_LOGOLIQ_PARTI_Winder_Lab_Logo[];
extern uint8_t PARTI_LOGOLIQ_PARTI_Logo[];
unsigned long startmillis;
String E_side = "RIGHT";
String display_page = "main";
bool show_time = true;
bool eject_button;
int LOG_COUNTER;
int SYNC_COUNTER;
unsigned long currentMillis;
unsigned long previousMillis;
int syncTime;
int min_now;
int prev_min;
int curr_min;
int hour_now;
bool refresh_page = true;
int timeouts = 0;
bool ok;
int default_lights_on;
int default_lights_off;
int default_touch_threshold;
int default_release_threshold;
int default_LOG_INTERVAL;
int dafault_SYNC_INTERVAL;
bool default_auto_cal;
int default_auto_cal_time;
int default_auto_cal_sec_since_last_lick;
bool auto_cal_flag = true;
unsigned long cal_timer;
String logging[6];
bool log_LN = true;
bool log_LD = true;
bool log_BN = true;
bool log_BD = true;
bool log_BLN = true;
bool log_BLD = true;
bool recording = false;
bool RFIDerrorLEFT;
bool RFIDerrorRIGHT;
bool RFID_lasterrorLEFT;
bool RFID_lasterrorRIGHT;
int tag1_count;
int lasttag1_count;
int tag2_count;
int lasttag2_count;
int notag1_count;
int notag2_count;
String licked_bottle;

const int intPin1 = 18;
const int intPin2 = 19;
String tag1;
String tag2;
String lasttag1;
String lasttag2;

//enough for 1 cage, 2 lickometers
#define sipper_number 2
unsigned long LickDuration[sipper_number];
unsigned long LickNumber[sipper_number];
unsigned long BoutNumber[sipper_number];
float BoutDuration[sipper_number];
float BoutLickNumber[sipper_number];
float BoutLickDuration[sipper_number];
float LickFrequency[sipper_number];
float ILI[sipper_number];
unsigned long Elapsedtime[sipper_number];
unsigned long total_LN[sipper_number];
int lick_bout_countdown[sipper_number];
unsigned long bout_timer[sipper_number];
unsigned long bout_start_timer[sipper_number];
bool in_bout[sipper_number];
bool licking[sipper_number];
unsigned long time_now[sipper_number];
unsigned long last_lick_time[sipper_number];


// how many minutes between grabbing data and logging it.
int LOG_INTERVAL = 1; // IN MINUTES! between entries (reduce to take more/faster data)
// how many milliseconds before writing the logged data permanently to disk
int SYNC_INTERVAL = 10; // IN MINUTES! between calls to flush() - to write data to the card
unsigned long emergency_counter = 60050; //1 minute +50ms

// Typically touch threshold > release threshold for debouncing
// Decrease value to increase sensitivity
int touch_threshold = 35; //default = 12
int release_threshold = 25; //default = 6
int current = 50; //Range of 1-63uA
int debounce = 0; //range 0-7
bool auto_cal = true;
int auto_cal_time = 7;
int auto_cal_sec_since_last_lick = 60;
int bout_cutoff = 1000;

int lights_on = 6; //time of day (24hr time) housing lights turn ON
int lights_off = 18; //time of day (24hr time) houseing lights turn OFF

//==========================================================================================

void setup() {

#ifdef DEBUG_LIQ || DEBUG_MPR121 || DEBUG_RFID || DEBUG_Screen || DEBUG_System
  Serial.begin(115200);
#endif

  pinMode(5, OUTPUT); // set screen brightness pin as output
  analogWrite(5, 255);
  pinMode(intPin1, INPUT_PULLUP);
  pinMode(intPin2, INPUT_PULLUP);

  set_defaults(); //save coded settings as the default settings

  start_display(); //initialize display

  start_rtc(); //iniitialize rtc

  start_rfid(); //iniitialize RFID readers

  WelcomeScreen(); //display welcome screen at startup

  set_brightness(); //set screen brightness depending on light cycle

}

//==========================================================================================

void loop() {

  main_menu_page();

  recording_page();

  pause_page();

  SD_error_page();

  settings_page();

}
```
