## Extended Data 1 for "A novel mouse home cage lickometer system reveals sex- and housing-based influences on alcohol drinking": Logo.c

```
#include <avr/pgmspace.h>
// 'welcome_logo', 240x173px
const unsigned char welcome_logo[] PROGMEM = {
  0x00, 0x00, 0x00, 0x00, 0x00, 0x00, 0x00, 0x00, 0x00, 0x00, 0x00, 0x00, 0x00, 0x00, 0x00, 0x00,
  0x00, 0x00, 0x00, 0x00, 0x00, 0x00, 0x00, 0x00, 0x00, 0x00, 0x00, 0x00, 0x00, 0x00, 0x00, 0x00,
  0x00, 0x00, 0x00, 0x00, 0x00, 0x00, 0x00, 0x00, 0x00, 0x00, 0x00, 0x00, 0x00, 0x00, 0x00, 0x00,
  0x00, 0x00, 0x00, 0x00, 0x00, 0x00, 0x00, 0x00, 0x00, 0x00, 0x00, 0x00, 0x00, 0x00, 0x00, 0x00,
  0x00, 0x00, 0x00, 0x00, 0x00, 0x00, 0x00, 0x00, 0x00, 0x00, 0x00, 0x00, 0x00, 0x00, 0x00, 0x00,
  0x00, 0x00, 0x00, 0x00, 0x00, 0x00, 0x00, 0x00, 0x00, 0x00, 0x00, 0x00, 0x00, 0x00, 0x00, 0x00,
  0x00, 0x00, 0x00, 0x00, 0x00, 0x00, 0x00, 0x00, 0x00, 0x00, 0x00, 0x00, 0x00, 0x00, 0x00, 0x00,
  0x00, 0x00, 0x00, 0x00, 0x00, 0x00, 0x00, 0x00, 0x00, 0x00, 0x00, 0x00, 0x00, 0x00, 0x00, 0x00,
  0x00, 0x00, 0x00, 0x00, 0x00, 0x00, 0x00, 0x00, 0x00, 0x00, 0x00, 0x00, 0x00, 0x00, 0x00, 0x00,
  0x00, 0x00, 0x00, 0x00, 0x00, 0x00, 0x00, 0x00, 0x00, 0x00, 0x00, 0x00, 0x00, 0x00, 0x00, 0x00,
  0x00, 0x00, 0x00, 0x00, 0x00, 0x00, 0x00, 0x00, 0x00, 0x00, 0x00, 0x00, 0x00, 0x00, 0x00, 0x00,
  0x00, 0x00, 0x00, 0x00, 0x00, 0x00, 0x00, 0x00, 0x00, 0x00, 0x00, 0x00, 0x00, 0x00, 0x00, 0x00,
  0x00, 0x00, 0x00, 0x00, 0x00, 0x00, 0x00, 0x00, 0x00, 0x00, 0x00, 0x00, 0x00, 0x00, 0x00, 0x00,
  0x00, 0x00, 0x00, 0x00, 0x00, 0x00, 0x00, 0x00, 0x00, 0x00, 0x00, 0x00, 0x00, 0x00, 0x00, 0x00,
  0x00, 0x00, 0x00, 0x00, 0x00, 0x00, 0x00, 0x00, 0x00, 0x00, 0x00, 0x00, 0x00, 0x00, 0x00, 0x00,
  0x00, 0x00, 0x00, 0x00, 0x00, 0x00, 0x00, 0x00, 0x00, 0x00, 0x00, 0x00, 0x00, 0x00, 0x00, 0x00,
  0x00, 0x00, 0x00, 0x00, 0x00, 0x00, 0x00, 0x00, 0x00, 0x00, 0x00, 0x00, 0x00, 0x00, 0x00, 0x00,
  0x00, 0x00, 0x00, 0x00, 0x00, 0x00, 0x00, 0x00, 0x00, 0x00, 0x00, 0x00, 0x00, 0x00, 0x00, 0x00,
  0x00, 0x00, 0x00, 0x00, 0x00, 0x00, 0x00, 0x00, 0x00, 0x00, 0x00, 0x00, 0x00, 0x00, 0x00, 0x00,
  0x00, 0x00, 0x00, 0x00, 0x00, 0x00, 0x00, 0x00, 0x00, 0x00, 0x00, 0x00, 0x00, 0x00, 0x00, 0x00,
  0x00, 0x00, 0x00, 0x00, 0x00, 0x00, 0x00, 0x00, 0x00, 0x00, 0x00, 0x00, 0x00, 0x00, 0x00, 0x00,
  0x00, 0x00, 0x00, 0x00, 0x00, 0x00, 0x00, 0x00, 0x00, 0x00, 0x00, 0x00, 0x00, 0x00, 0x00, 0x00,
  0x00, 0x00, 0x00, 0x00, 0x00, 0x00, 0x00, 0x00, 0x00, 0x00, 0x00, 0x00, 0x00, 0x00, 0x00, 0x00,
  0x00, 0x00, 0x00, 0x00, 0x00, 0x00, 0x00, 0x00, 0x00, 0x00, 0x00, 0x00, 0x00, 0x00, 0x00, 0x00,
  0x00, 0x00, 0x00, 0x00, 0x00, 0x00, 0x00, 0x00, 0x00, 0x00, 0x00, 0x00, 0x00, 0x00, 0x00, 0x00,
  0x00, 0x00, 0x00, 0x00, 0x00, 0x00, 0x00, 0x00, 0x00, 0x00, 0x00, 0x00, 0x00, 0x00, 0x00, 0x00,
  0x00, 0x00, 0x00, 0x00, 0x00, 0x00, 0x00, 0x00, 0x00, 0x00, 0x00, 0x00, 0x00, 0x00, 0x00, 0x00,
  0x00, 0x00, 0x00, 0x00, 0x00, 0x00, 0x00, 0x00, 0x00, 0x00, 0x00, 0x00, 0x00, 0x00, 0x00, 0x00,
  0x00, 0x00, 0x00, 0x00, 0x00, 0x00, 0x00, 0x00, 0x00, 0x00, 0x00, 0x00, 0x00, 0x00, 0x00, 0x00,
  0x00, 0x00, 0x00, 0x00, 0x00, 0x00, 0x00, 0x00, 0x00, 0x00, 0x00, 0x00, 0x00, 0x00, 0x00, 0x00,
  0x00, 0x00, 0x00, 0x00, 0x00, 0x00, 0x00, 0x00, 0x00, 0x00, 0x00, 0x00, 0x00, 0x00, 0x00, 0x00,
  0x00, 0x00, 0x00, 0x00, 0x00, 0x00, 0x00, 0x00, 0x00, 0x00, 0x00, 0x00, 0x00, 0x00, 0x00, 0x00,
  0x00, 0x00, 0x00, 0x00, 0x00, 0x00, 0x00, 0x00, 0x00, 0x00, 0x00, 0x00, 0x00, 0x00, 0x00, 0x00,
  0x00, 0x00, 0x00, 0x00, 0x00, 0x00, 0x00, 0x00, 0x00, 0x00, 0x00, 0x00, 0x00, 0x00, 0x00, 0x00,
  0x00, 0x00, 0x00, 0x00, 0x00, 0x00, 0x00, 0x00, 0x00, 0x00, 0x00, 0x00, 0x00, 0x00, 0x00, 0x00,
  0x00, 0x00, 0x00, 0x00, 0x00, 0x00, 0x00, 0x00, 0x00, 0x00, 0x00, 0x00, 0x00, 0x00, 0x00, 0x00,
  0x00, 0x00, 0x00, 0x00, 0x00, 0x00, 0x00, 0x00, 0x00, 0x00, 0x00, 0x00, 0x00, 0x00, 0x00, 0x00,
  0x00, 0x00, 0x00, 0x00, 0x00, 0x00, 0x00, 0x00, 0x00, 0x00, 0x00, 0x00, 0x00, 0x00, 0x00, 0x00,
  0x00, 0x00, 0x00, 0x00, 0x00, 0x00, 0x00, 0x00, 0x00, 0x00, 0x00, 0x00, 0x00, 0x00, 0x00, 0x00,
  0x00, 0x00, 0x00, 0x00, 0x00, 0x00, 0x00, 0x00, 0x00, 0x00, 0x00, 0x00, 0x00, 0x00, 0x00, 0x00,
  0x00, 0x00, 0x00, 0x00, 0x00, 0x00, 0x00, 0x00, 0x00, 0x00, 0x00, 0x00, 0x00, 0x00, 0x00, 0x00,
  0x00, 0x00, 0x00, 0x00, 0x00, 0x00, 0x00, 0x00, 0x00, 0x00, 0x00, 0x00, 0x00, 0x00, 0x00, 0x00,
  0x00, 0x00, 0x00, 0x00, 0x00, 0x00, 0x00, 0x00, 0x00, 0x00, 0x00, 0x00, 0x00, 0x00, 0x00, 0x00,
  0x00, 0x00, 0x00, 0x00, 0x00, 0x00, 0x00, 0x00, 0x00, 0x00, 0x00, 0x00, 0x00, 0x00, 0x00, 0x00,
  0x00, 0x00, 0x00, 0x00, 0x00, 0x00, 0x00, 0x00, 0x00, 0x00, 0x00, 0x00, 0x00, 0x00, 0x00, 0x00,
  0x00, 0x00, 0x00, 0x00, 0x00, 0x00, 0x00, 0x00, 0x00, 0x00, 0x00, 0x00, 0x00, 0x00, 0x00, 0x00,
  0x00, 0x00, 0x00, 0x00, 0x00, 0x00, 0x00, 0x00, 0x00, 0x00, 0x00, 0x00, 0x00, 0x00, 0x00, 0x00,
  0x00, 0x00, 0x00, 0x00, 0x00, 0x00, 0x00, 0x00, 0x00, 0x00, 0x00, 0x00, 0x00, 0x00, 0x00, 0x00,
  0x00, 0x00, 0x00, 0x00, 0x00, 0x00, 0x00, 0x00, 0x00, 0x00, 0x00, 0x00, 0x00, 0x00, 0x00, 0x00,
  0x00, 0x00, 0x00, 0x00, 0x00, 0x00, 0x00, 0x00, 0x00, 0x00, 0x00, 0x00, 0x00, 0x00, 0x00, 0x00,
  0x00, 0x00, 0x00, 0x00, 0x00, 0x00, 0x00, 0x00, 0x00, 0x00, 0x00, 0x00, 0x00, 0x00, 0x00, 0x00,
  0x00, 0x00, 0x00, 0x00, 0x00, 0x00, 0x00, 0x00, 0x00, 0x00, 0x00, 0x00, 0x00, 0x00, 0x00, 0x00,
  0x00, 0x00, 0x00, 0x00, 0x00, 0x00, 0x00, 0x00, 0x00, 0x00, 0x00, 0x00, 0x00, 0x00, 0x00, 0x00,
  0x00, 0x00, 0x00, 0x00, 0x00, 0x00, 0x00, 0x00, 0x00, 0x00, 0x00, 0x00, 0x00, 0x00, 0x00, 0x00,
  0x00, 0x00, 0x00, 0x00, 0x00, 0x00, 0x00, 0x00, 0x00, 0x00, 0x00, 0x00, 0x00, 0x00, 0x00, 0x00,
  0x00, 0x00, 0x00, 0x00, 0x00, 0x00, 0x00, 0x00, 0x00, 0x00, 0x00, 0x00, 0x00, 0x00, 0x00, 0x00,
  0x00, 0x00, 0x00, 0x00, 0x00, 0x00, 0x00, 0x00, 0x0f, 0xe0, 0x00, 0x01, 0xfc, 0x00, 0x0f, 0xff,
  0xf8, 0x00, 0x00, 0x00, 0x01, 0xfc, 0x00, 0x03, 0xf8, 0x0f, 0xff, 0xff, 0xf0, 0x00, 0x00, 0x00,
  0x00, 0x00, 0x00, 0x00, 0x00, 0x00, 0x0f, 0xe0, 0x00, 0x01, 0xfc, 0x00, 0x3f, 0xff, 0xfe, 0x00,
  0x00, 0x00, 0x01, 0xfc, 0x00, 0x03, 0xf8, 0x0f, 0xff, 0xff, 0xfc, 0x00, 0x00, 0x00, 0x00, 0x00,
  0x00, 0x00, 0x00, 0x00, 0x0f, 0xe0, 0x00, 0x01, 0xfc, 0x00, 0x7f, 0xff, 0xff, 0x00, 0x00, 0x00,
  0x01, 0xfc, 0x00, 0x03, 0xf8, 0x0f, 0xff, 0xff, 0xfe, 0x00, 0x00, 0x00, 0x00, 0x00, 0x00, 0x00,
  0x00, 0x00, 0x0f, 0xe0, 0x00, 0x01, 0xfc, 0x00, 0xff, 0xff, 0xff, 0x80, 0x00, 0x00, 0x01, 0xfc,
  0x00, 0x03, 0xf8, 0x0f, 0xff, 0xff, 0xff, 0x00, 0x00, 0x00, 0x00, 0x00, 0x00, 0x00, 0x00, 0x00,
  0x0f, 0xe0, 0x00, 0x01, 0xfc, 0x01, 0xff, 0xff, 0xff, 0xc0, 0x00, 0x00, 0x01, 0xfc, 0x00, 0x03,
  0xf8, 0x0f, 0xff, 0xff, 0xff, 0x80, 0x00, 0x00, 0x00, 0x00, 0x00, 0x00, 0x00, 0x00, 0x0f, 0xe0,
  0x00, 0x01, 0xfc, 0x03, 0xff, 0xff, 0xff, 0xe0, 0x00, 0x00, 0x01, 0xfc, 0x00, 0x03, 0xf8, 0x0f,
  0xff, 0xff, 0xff, 0xc0, 0x00, 0x00, 0x00, 0x00, 0x00, 0x00, 0x00, 0x00, 0x0f, 0xe0, 0x00, 0x01,
  0xfc, 0x03, 0xff, 0xff, 0xff, 0xe0, 0x00, 0x00, 0x01, 0xfc, 0x00, 0x03, 0xf8, 0x0f, 0xff, 0xff,
  0xff, 0xc0, 0x00, 0x00, 0x00, 0x00, 0x00, 0x00, 0x00, 0x00, 0x0f, 0xe0, 0x00, 0x01, 0xfc, 0x03,
  0xf8, 0x00, 0x0f, 0xe0, 0x00, 0x00, 0x01, 0xfc, 0x00, 0x03, 0xf8, 0x0f, 0xe0, 0x00, 0x1f, 0xc0,
  0x00, 0x00, 0x00, 0x00, 0x00, 0x00, 0x00, 0x00, 0x0f, 0xe0, 0x00, 0x01, 0xfc, 0x07, 0xf0, 0x00,
  0x07, 0xf0, 0x00, 0x00, 0x01, 0xfc, 0x00, 0x03, 0xf8, 0x0f, 0xe0, 0x00, 0x0f, 0xe0, 0x00, 0x00,
  0x00, 0x00, 0x00, 0x00, 0x00, 0x00, 0x0f, 0xe0, 0x00, 0x01, 0xfc, 0x07, 0xf0, 0x00, 0x07, 0xf0,
  0x00, 0x00, 0x01, 0xfc, 0x00, 0x03, 0xf8, 0x0f, 0xe0, 0x00, 0x0f, 0xe0, 0x00, 0x00, 0x00, 0x00,
  0x00, 0x00, 0x00, 0x00, 0x0f, 0xe0, 0x00, 0x01, 0xfc, 0x07, 0xf0, 0x00, 0x07, 0xf0, 0x00, 0x00,
  0x01, 0xfc, 0x00, 0x03, 0xf8, 0x0f, 0xe0, 0x00, 0x0f, 0xe0, 0x00, 0x00, 0x00, 0x00, 0x00, 0x00,
  0x00, 0x00, 0x0f, 0xe0, 0x00, 0x01, 0xfc, 0x07, 0xf0, 0x00, 0x07, 0xf0, 0x00, 0x00, 0x01, 0xfc,
  0x00, 0x03, 0xf8, 0x0f, 0xe0, 0x00, 0x0f, 0xe0, 0x00, 0x00, 0x00, 0x00, 0x00, 0x00, 0x00, 0x00,
  0x0f, 0xe0, 0x00, 0x01, 0xfc, 0x07, 0xf0, 0x00, 0x07, 0xf0, 0x00, 0x00, 0x01, 0xfc, 0x00, 0x03,
  0xf8, 0x0f, 0xe0, 0x00, 0x0f, 0xe0, 0x00, 0x00, 0x00, 0x00, 0x00, 0x00, 0x00, 0x00, 0x0f, 0xe0,
  0x00, 0x01, 0xfc, 0x07, 0xf0, 0x00, 0x07, 0xf0, 0x00, 0x00, 0x01, 0xfc, 0x00, 0x03, 0xf8, 0x0f,
  0xe0, 0x00, 0x0f, 0xe0, 0x00, 0x00, 0x00, 0x00, 0x00, 0x00, 0x00, 0x00, 0x0f, 0xe0, 0x00, 0x01,
  0xfc, 0x07, 0xf0, 0x00, 0x07, 0xf0, 0x00, 0x00, 0x01, 0xfc, 0x00, 0x03, 0xf8, 0x0f, 0xe0, 0x00,
  0x0f, 0xe0, 0x00, 0x00, 0x00, 0x00, 0x00, 0x00, 0x00, 0x00, 0x0f, 0xe0, 0x00, 0x01, 0xfc, 0x07,
  0xf0, 0x00, 0x07, 0xf0, 0x00, 0x00, 0x01, 0xfc, 0x00, 0x03, 0xf8, 0x0f, 0xe0, 0x00, 0x0f, 0xe0,
  0x00, 0x00, 0x00, 0x00, 0x00, 0x00, 0x00, 0x00, 0x0f, 0xe0, 0x00, 0x01, 0xfc, 0x07, 0xf0, 0x00,
  0x07, 0xf0, 0x00, 0x00, 0x01, 0xfc, 0x00, 0x03, 0xf8, 0x0f, 0xe0, 0x00, 0x0f, 0xe0, 0x00, 0x00,
  0x00, 0x00, 0x00, 0x00, 0x00, 0x00, 0x0f, 0xe0, 0x00, 0x01, 0xfc, 0x07, 0xf0, 0x00, 0x07, 0xf0,
  0x00, 0x00, 0x01, 0xfc, 0x00, 0x03, 0xf8, 0x0f, 0xe0, 0x00, 0x0f, 0xe0, 0x00, 0x00, 0x00, 0x00,
  0x00, 0x00, 0x00, 0x00, 0x0f, 0xe0, 0x00, 0x01, 0xfc, 0x07, 0xf0, 0x00, 0x07, 0xf0, 0x00, 0x00,
  0x01, 0xfc, 0x00, 0x03, 0xf8, 0x0f, 0xe0, 0x00, 0x0f, 0xe0, 0x00, 0x00, 0x00, 0x00, 0x00, 0x00,
  0x00, 0x00, 0x0f, 0xe0, 0x00, 0x01, 0xfc, 0x07, 0xf0, 0x00, 0x07, 0xf0, 0x00, 0x00, 0x01, 0xfc,
  0x00, 0x03, 0xf8, 0x0f, 0xe0, 0x00, 0x0f, 0xe0, 0x00, 0x00, 0x00, 0x00, 0x00, 0x00, 0x00, 0x00,
  0x0f, 0xe0, 0x00, 0x01, 0xfc, 0x07, 0xf0, 0x00, 0x07, 0xf0, 0x00, 0x00, 0x01, 0xfc, 0x00, 0x03,
  0xf8, 0x0f, 0xe0, 0x00, 0x0f, 0xe0, 0x00, 0x00, 0x00, 0x00, 0x00, 0x00, 0x00, 0x00, 0x0f, 0xe0,
  0x00, 0x01, 0xfc, 0x07, 0xf0, 0x00, 0x07, 0xf0, 0x00, 0x00, 0x01, 0xfc, 0x00, 0x03, 0xf8, 0x0f,
  0xe0, 0x00, 0x0f, 0xe0, 0x00, 0x00, 0x00, 0x00, 0x00, 0x00, 0x00, 0x00, 0x0f, 0xe0, 0x00, 0x01,
  0xfc, 0x07, 0xf0, 0x00, 0x07, 0xf0, 0x00, 0x00, 0x01, 0xfc, 0x00, 0x03, 0xf8, 0x0f, 0xe0, 0x00,
  0x0f, 0xe0, 0x00, 0x00, 0x00, 0x00, 0x00, 0x00, 0x00, 0x00, 0x0f, 0xe0, 0x00, 0x01, 0xfc, 0x07,
  0xf0, 0x00, 0x07, 0xf0, 0x00, 0x00, 0x01, 0xfc, 0x00, 0x03, 0xf8, 0x0f, 0xe0, 0x00, 0x0f, 0xe0,
  0x00, 0x00, 0x00, 0x00, 0x00, 0x00, 0x00, 0x00, 0x0f, 0xe0, 0x00, 0x01, 0xfc, 0x07, 0xf0, 0x00,
  0x07, 0xf0, 0x00, 0x00, 0x01, 0xfc, 0x00, 0x03, 0xf8, 0x0f, 0xe0, 0x00, 0x0f, 0xe0, 0x00, 0x00,
  0x00, 0x00, 0x00, 0x00, 0x00, 0x00, 0x0f, 0xe0, 0x00, 0x01, 0xfc, 0x07, 0xf0, 0x00, 0x07, 0xf0,
  0x00, 0x00, 0x01, 0xfc, 0x00, 0x03, 0xf8, 0x0f, 0xe0, 0x00, 0x0f, 0xe0, 0x00, 0x00, 0x00, 0x00,
  0x00, 0x00, 0x00, 0x00, 0x0f, 0xe0, 0x00, 0x01, 0xfc, 0x07, 0xf0, 0x00, 0x07, 0xf0, 0x00, 0x00,
  0x01, 0xfc, 0x00, 0x03, 0xf8, 0x0f, 0xe0, 0x00, 0x0f, 0xe0, 0x00, 0x00, 0x00, 0x00, 0x00, 0x00,
  0x00, 0x00, 0x0f, 0xe0, 0x00, 0x01, 0xfc, 0x07, 0xf0, 0x00, 0x07, 0xf0, 0x00, 0x00, 0x01, 0xfc,
  0x00, 0x03, 0xf8, 0x0f, 0xe0, 0x00, 0x0f, 0xe0, 0x00, 0x00, 0x00, 0x00, 0x00, 0x00, 0x00, 0x00,
  0x0f, 0xe0, 0x00, 0x01, 0xfc, 0x07, 0xf0, 0x00, 0x07, 0xf0, 0x00, 0x00, 0x01, 0xfc, 0x00, 0x03,
  0xf8, 0x0f, 0xe0, 0x00, 0x0f, 0xe0, 0x00, 0x00, 0x00, 0x00, 0x00, 0x00, 0x00, 0x00, 0x0f, 0xe0,
  0x00, 0x01, 0xfc, 0x07, 0xf0, 0x00, 0x07, 0xf0, 0x00, 0x00, 0x01, 0xfc, 0x00, 0x03, 0xf8, 0x0f,
  0xe0, 0x00, 0x0f, 0xe0, 0x00, 0x00, 0x00, 0x00, 0x00, 0x00, 0x00, 0x00, 0x0f, 0xe0, 0x00, 0x01,
  0xfc, 0x07, 0xf0, 0x00, 0x07, 0xf0, 0x00, 0x00, 0x01, 0xfc, 0x00, 0x03, 0xf8, 0x0f, 0xe0, 0x00,
  0x0f, 0xe0, 0x00, 0x00, 0x00, 0x00, 0x00, 0x00, 0x00, 0x00, 0x0f, 0xe0, 0x00, 0x01, 0xfc, 0x07,
  0xf0, 0x00, 0x07, 0xf0, 0x00, 0x00, 0x0f, 0xff, 0xff, 0xff, 0xf8, 0x0f, 0xe0, 0x00, 0x0f, 0xe0,
  0x00, 0x00, 0x00, 0x00, 0x00, 0x00, 0x00, 0x00, 0x0f, 0xe0, 0x00, 0x01, 0xfc, 0x07, 0xf0, 0x00,
  0x07, 0xf0, 0x00, 0x00, 0x0f, 0xff, 0xff, 0xff, 0xf8, 0x0f, 0xe0, 0x00, 0x0f, 0xe0, 0x00, 0x00,
  0x00, 0x00, 0x00, 0x00, 0x00, 0x00, 0x0f, 0xe0, 0x00, 0x01, 0xfc, 0x07, 0xf0, 0x00, 0x07, 0xf0,
  0x00, 0x00, 0x0f, 0xff, 0xff, 0xff, 0xf8, 0x0f, 0xe0, 0x00, 0x0f, 0xe0, 0x00, 0x00, 0x00, 0x00,
  0x00, 0x00, 0x00, 0x00, 0x0f, 0xe0, 0x00, 0x01, 0xfc, 0x07, 0xf0, 0x00, 0x07, 0xf0, 0x00, 0x00,
  0x0f, 0xff, 0xff, 0xff, 0xf8, 0x0f, 0xe0, 0x00, 0x0f, 0xe0, 0x00, 0x00, 0x00, 0x00, 0x00, 0x00,
  0x00, 0x00, 0x0f, 0xe0, 0x00, 0x01, 0xfc, 0x07, 0xf0, 0x00, 0x07, 0xf0, 0x00, 0x00, 0x0f, 0xff,
  0xff, 0xff, 0xf8, 0x0f, 0xe0, 0x00, 0x0f, 0xe0, 0x00, 0x00, 0x00, 0x00, 0x00, 0x00, 0x00, 0x00,
  0x0f, 0xe0, 0x00, 0x01, 0xfc, 0x07, 0xf0, 0x00, 0x07, 0xf0, 0x00, 0x00, 0x0f, 0xff, 0xff, 0xff,
  0xf8, 0x0f, 0xe0, 0x00, 0x0f, 0xe0, 0x00, 0x00, 0x00, 0x00, 0x00, 0x00, 0x00, 0x00, 0x0f, 0xe0,
  0x00, 0x01, 0xfc, 0x07, 0xf0, 0x00, 0x07, 0xf0, 0x00, 0x00, 0x0f, 0xff, 0xff, 0xff, 0xf8, 0x0f,
  0xe0, 0x00, 0x0f, 0xe0, 0x00, 0x00, 0x00, 0x00, 0x00, 0x00, 0x00, 0x00, 0x0f, 0xe0, 0x00, 0x01,
  0xfc, 0x07, 0xf0, 0x00, 0x07, 0xf0, 0x00, 0x00, 0x01, 0xfc, 0x00, 0x03, 0xf8, 0x0f, 0xe0, 0x00,
  0x0f, 0xe0, 0x00, 0x00, 0x00, 0x00, 0x00, 0x00, 0x00, 0x00, 0x0f, 0xe0, 0x00, 0x01, 0xfc, 0x07,
  0xf0, 0x00, 0x07, 0xf0, 0x00, 0x00, 0x01, 0xfc, 0x00, 0x03, 0xf8, 0x0f, 0xe0, 0x00, 0x0f, 0xe0,
  0x00, 0x00, 0x00, 0x00, 0x00, 0x00, 0x00, 0x00, 0x0f, 0xe0, 0x00, 0x01, 0xfc, 0x07, 0xf0, 0x00,
  0x07, 0xf0, 0x00, 0x00, 0x01, 0xfc, 0x00, 0x03, 0xf8, 0x0f, 0xe0, 0x00, 0x0f, 0xe0, 0x00, 0x00,
  0x00, 0x00, 0x00, 0x00, 0x00, 0x00, 0x0f, 0xe0, 0x00, 0x01, 0xfc, 0x07, 0xf0, 0x00, 0x07, 0xf0,
  0x00, 0x00, 0x01, 0xfc, 0x00, 0x03, 0xf8, 0x0f, 0xe0, 0x00, 0x0f, 0xe0, 0x00, 0x00, 0x00, 0x00,
  0x00, 0x00, 0x00, 0x00, 0x0f, 0xe0, 0x00, 0x01, 0xfc, 0x07, 0xf0, 0x00, 0x07, 0xf0, 0x00, 0x00,
  0x01, 0xfc, 0x00, 0x03, 0xf8, 0x0f, 0xe0, 0x00, 0x0f, 0xe0, 0x00, 0x00, 0x00, 0x00, 0x00, 0x00,
  0x00, 0x00, 0x0f, 0xe0, 0x00, 0x01, 0xfc, 0x07, 0xf8, 0x00, 0x0f, 0xe0, 0x00, 0x00, 0x01, 0xfc,
  0x00, 0x03, 0xf8, 0x0f, 0xe0, 0x00, 0x1f, 0xc0, 0x00, 0x00, 0x00, 0x00, 0x00, 0x00, 0x00, 0x00,
  0x0f, 0xff, 0xff, 0x01, 0xfc, 0x03, 0xff, 0xff, 0xff, 0xe0, 0x00, 0x00, 0x01, 0xfc, 0x00, 0x03,
  0xf8, 0x0f, 0xff, 0xff, 0xff, 0xc0, 0x00, 0x00, 0x00, 0x00, 0x00, 0x00, 0x00, 0x00, 0x0f, 0xff,
  0xff, 0x01, 0xfc, 0x03, 0xff, 0xff, 0xff, 0xe0, 0x00, 0x00, 0x01, 0xfc, 0x00, 0x03, 0xf8, 0x0f,
  0xff, 0xff, 0xff, 0xc0, 0x00, 0x00, 0x00, 0x00, 0x00, 0x00, 0x00, 0x00, 0x0f, 0xff, 0xff, 0x01,
  0xfc, 0x01, 0xff, 0xff, 0xff, 0xc0, 0x00, 0x00, 0x01, 0xfc, 0x00, 0x03, 0xf8, 0x0f, 0xff, 0xff,
  0xff, 0x80, 0x00, 0x00, 0x00, 0x00, 0x00, 0x00, 0x00, 0x00, 0x0f, 0xff, 0xff, 0x01, 0xfc, 0x00,
  0xff, 0xff, 0xff, 0x80, 0x00, 0x00, 0x01, 0xfc, 0x00, 0x03, 0xf8, 0x0f, 0xff, 0xff, 0xff, 0x00,
  0x00, 0x00, 0x00, 0x00, 0x00, 0x00, 0x00, 0x00, 0x0f, 0xff, 0xff, 0x01, 0xfc, 0x00, 0x7f, 0xff,
  0xff, 0x00, 0x00, 0x00, 0x01, 0xfc, 0x00, 0x03, 0xf8, 0x0f, 0xff, 0xff, 0xfe, 0x00, 0x00, 0x00,
  0x00, 0x00, 0x00, 0x00, 0x00, 0x00, 0x0f, 0xff, 0xff, 0x01, 0xfc, 0x00, 0x3f, 0xff, 0xfe, 0x00,
  0x00, 0x00, 0x01, 0xfc, 0x00, 0x03, 0xf8, 0x0f, 0xff, 0xff, 0xfc, 0x00, 0x00, 0x00, 0x00, 0x00,
  0x00, 0x00, 0x00, 0x00, 0x0f, 0xff, 0xff, 0x01, 0xfc, 0x00, 0x0f, 0xff, 0xf8, 0x00, 0x00, 0x00,
  0x01, 0xfc, 0x00, 0x03, 0xf8, 0x0f, 0xff, 0xff, 0xf0, 0x00, 0x00, 0x00, 0x00, 0x00, 0x00, 0x00,
  0x00, 0x00, 0x00, 0x00, 0x00, 0x00, 0x00, 0x00, 0x00, 0x3f, 0xf0, 0x00, 0x00, 0x00, 0x00, 0x00,
  0x00, 0x00, 0x00, 0x00, 0x00, 0x00, 0x00, 0x00, 0x00, 0x00, 0x00, 0x00, 0x00, 0x00, 0x00, 0x00,
  0x00, 0x00, 0x00, 0x00, 0x00, 0x00, 0x00, 0x1f, 0xf8, 0x00, 0x00, 0x00, 0x00, 0x00, 0x00, 0x00,
  0x00, 0x00, 0x00, 0x00, 0x00, 0x00, 0x00, 0x00, 0x00, 0x00, 0x00, 0x00, 0x00, 0x00, 0x00, 0x00,
  0x00, 0x00, 0x00, 0x00, 0x00, 0x0f, 0xfc, 0x00, 0x00, 0x00, 0x00, 0x00, 0x00, 0x00, 0x00, 0x00,
  0x00, 0x00, 0x00, 0x00, 0x00, 0x00, 0x00, 0x00, 0x00, 0x00, 0x00, 0x00, 0x00, 0x00, 0x00, 0x00,
  0x00, 0x00, 0x00, 0x07, 0xfc, 0x00, 0x00, 0x00, 0x00, 0x00, 0x00, 0x00, 0x00, 0x00, 0x00, 0x00,
  0x00, 0x00, 0x00, 0x00, 0x00, 0x00, 0x00, 0x00, 0x00, 0x00, 0x00, 0x00, 0x00, 0x00, 0x00, 0x00,
  0x00, 0x03, 0xf8, 0x00, 0x00, 0x00, 0x00, 0x00, 0x00, 0x00, 0x00, 0x00, 0x00, 0x00, 0x00, 0x00,
  0x00, 0x00, 0x00, 0x00, 0x00, 0x00, 0x00, 0x00, 0x00, 0x00, 0x00, 0x00, 0x00, 0x00, 0x00, 0x01,
  0xf0, 0x00, 0x00, 0x00, 0x00, 0x00, 0x00, 0x00, 0x00, 0x00, 0x00, 0x00, 0x00, 0x00, 0x00, 0x00,
  0x00, 0x00, 0x00, 0x00, 0x00, 0x00, 0x00, 0x00, 0x00, 0x00, 0x00, 0x00, 0x00, 0x00, 0xe0, 0x00,
  0x00, 0x00, 0x00, 0x00, 0x00, 0x00, 0x00, 0x00, 0x00, 0x00, 0x00, 0x00, 0x00, 0x00, 0x00, 0x00,
  0x00, 0x00, 0x00, 0x00, 0x00, 0x00, 0x00, 0x00, 0x00, 0x00, 0x00, 0x00, 0x40, 0x00, 0x00, 0x00,
  0x00, 0x00, 0x00, 0x00, 0x00, 0x00, 0x00, 0x00, 0x00, 0x00, 0x00, 0x00, 0x00, 0x00, 0x00, 0x00,
  0x00, 0x00, 0x00, 0x00, 0x00, 0x00, 0x00, 0x00, 0x00, 0x00, 0x00, 0x00, 0x00, 0x00, 0x00, 0x00,
  0x00, 0x00, 0x00, 0x00, 0x00, 0x00, 0x00, 0x00, 0x00, 0x00, 0x00, 0x00, 0x00, 0x00, 0x00, 0x00,
  0x00, 0x00, 0x00, 0x00, 0x00, 0x00, 0x00, 0x00, 0x00, 0x00, 0x00, 0x00, 0x00, 0x00, 0x00, 0x00,
  0x00, 0x00, 0x00, 0x00, 0x00, 0x00, 0x00, 0x00, 0x00, 0x00, 0x00, 0x00, 0x00, 0x00, 0x00, 0x00,
  0x00, 0x00, 0x00, 0x00, 0x00, 0x00, 0x00, 0x00, 0x00, 0x00, 0x00, 0x00, 0x00, 0x00, 0x00, 0x00,
  0x00, 0x00, 0x00, 0x00, 0x00, 0x00, 0x00, 0x00, 0x00, 0x00, 0x00, 0x00, 0x00, 0x00, 0x00, 0x00,
  0x00, 0x00, 0x00, 0x00, 0x00, 0x00, 0x00, 0x00, 0x00, 0x00, 0x00, 0x00, 0x00, 0x00, 0x00, 0x00,
  0x00, 0x00, 0x00, 0x00, 0x00, 0x00, 0x00, 0x00, 0x00, 0x00, 0x00, 0x00, 0x00, 0x00, 0x00, 0x00,
  0x00, 0x00, 0x00, 0x00, 0x00, 0x00, 0x00, 0x00, 0x00, 0x00, 0x00, 0x00, 0x00, 0x00, 0x00, 0x00,
  0x00, 0x00, 0x00, 0x00, 0x00, 0x00, 0x00, 0x00, 0x00, 0x00, 0x00, 0x00, 0x00, 0x00, 0x00, 0x00,
  0x00, 0x00, 0x00, 0x00, 0x00, 0x00, 0x00, 0x00, 0x00, 0x00, 0x00, 0x00, 0x00, 0x00, 0x00, 0x00,
  0x00, 0x00, 0x00, 0x00, 0x00, 0x00, 0x00, 0x00, 0x00, 0x00, 0x00, 0x00, 0x00, 0x00, 0x00, 0x00,
  0x00, 0x00, 0x00, 0x00, 0x00, 0x00, 0x00, 0x00, 0x00, 0x00, 0x00, 0x00, 0x00, 0x00, 0x00, 0x00,
  0x00, 0x00, 0x00, 0x00, 0x00, 0x00, 0x00, 0x00, 0x00, 0x00, 0x00, 0x00, 0x00, 0x00, 0x00, 0x00,
  0x00, 0x00, 0x00, 0x00, 0x00, 0x00, 0x00, 0x00, 0x00, 0x00, 0x00, 0x00, 0x00, 0x00, 0x00, 0x00,
  0x00, 0x00, 0x00, 0x00, 0x00, 0x00, 0x00, 0x00, 0x00, 0x00, 0x00, 0x00, 0x00, 0x00, 0x00, 0x00,
  0x00, 0x00, 0x00, 0x00, 0x00, 0x00, 0x00, 0x00, 0x00, 0x00, 0x00, 0x00, 0x00, 0x00, 0x00, 0x00,
  0x00, 0x00, 0x00, 0x00, 0x00, 0x00, 0x00, 0x00, 0x00, 0x00, 0x00, 0x00, 0x00, 0x00, 0x00, 0x00,
  0x00, 0x00, 0x00, 0x00, 0x00, 0x00, 0x00, 0x00, 0x00, 0x00, 0x00, 0x00, 0x00, 0x00, 0x00, 0x00,
  0x00, 0x00, 0x00, 0x00, 0x00, 0x00, 0x00, 0x00, 0x00, 0x00, 0x00, 0x00, 0x00, 0x00, 0x00, 0x00,
  0x00, 0x00, 0x00, 0x00, 0x00, 0x00, 0x00, 0x00, 0x00, 0x00, 0x00, 0x00, 0x00, 0x00, 0x00, 0x00,
  0x00, 0x00, 0x00, 0x00, 0x00, 0x00, 0x00, 0x00, 0x00, 0x00, 0x00, 0x00, 0x00, 0x00, 0x00, 0x00,
  0x00, 0x00, 0x00, 0x00, 0x00, 0x00, 0x00, 0x00, 0x00, 0x00, 0x00, 0x00, 0x00, 0x00, 0x00, 0x00,
  0x00, 0x00, 0x00, 0x00, 0x00, 0x00, 0x00, 0x00, 0x00, 0x00, 0x00, 0x00, 0x00, 0x00, 0x00, 0x00,
  0x00, 0x00, 0x00, 0x00, 0x00, 0x00, 0x00, 0x00, 0x00, 0x00, 0x00, 0x00, 0x00, 0x00, 0x00, 0x00,
  0x00, 0x00, 0x00, 0x00, 0x00, 0x00, 0x00, 0x00, 0x00, 0x00, 0x00, 0x00, 0x00, 0x00, 0x00, 0x00,
  0x00, 0x00, 0x00, 0x00, 0x00, 0x00, 0x00, 0x00, 0x00, 0x00, 0x00, 0x00, 0x00, 0x00, 0x00, 0x00,
  0x00, 0x00, 0x00, 0x00, 0x00, 0x00, 0x00, 0x00, 0x00, 0x00, 0x00, 0x00, 0x00, 0x00, 0x00, 0x00,
  0x00, 0x00, 0x00, 0x00, 0x00, 0x00, 0x00, 0x00, 0x00, 0x00, 0x00, 0x00, 0x00, 0x00, 0x00, 0x00,
  0x00, 0x00, 0x00, 0x00, 0x00, 0x00, 0x00, 0x00, 0x00, 0x00, 0x00, 0x00, 0x00, 0x00, 0x00, 0x00,
  0x00, 0x00, 0x00, 0x00, 0x00, 0x00, 0x00, 0x00, 0x00, 0x00, 0x00, 0x00, 0x00, 0x00, 0x00, 0x00,
  0x00, 0x00, 0x00, 0x00, 0x00, 0x00, 0x00, 0x00, 0x00, 0x00, 0x00, 0x00, 0x00, 0x00, 0x00, 0x00,
  0x00, 0x00, 0x00, 0x00, 0x00, 0x00, 0x00, 0x00, 0x00, 0x00, 0x00, 0x00, 0x00, 0x00, 0x00, 0x00,
  0x00, 0x00, 0x00, 0x00, 0x00, 0x00, 0x00, 0x00, 0x00, 0x00, 0x00, 0x00, 0x00, 0x00, 0x00, 0x00,
  0x00, 0x00, 0x00, 0x00, 0x00, 0x00, 0x00, 0x00, 0x00, 0x00, 0x00, 0x00, 0x00, 0x00, 0x00, 0x00,
  0x00, 0x00, 0x00, 0x00, 0x00, 0x00, 0x00, 0x00, 0x00, 0x00, 0x00, 0x00, 0x00, 0x00, 0x00, 0x00,
  0x00, 0x00, 0x00, 0x00, 0x00, 0x00, 0x00, 0x00, 0x00, 0x00, 0x00, 0x00, 0x00, 0x00, 0x00, 0x00,
  0x00, 0x00, 0x00, 0x00, 0x00, 0x00, 0x00, 0x00, 0x00, 0x00, 0x00, 0x00, 0x00, 0x00, 0x00, 0x00,
  0x00, 0x00, 0x00, 0x00, 0x00, 0x00, 0x00, 0x00, 0x00, 0x00, 0x00, 0x00, 0x00, 0x00, 0x00, 0x00,
  0x00, 0x00, 0x00, 0x00, 0x00, 0x00, 0x00, 0x00, 0x00, 0x00, 0x00, 0x00, 0x00, 0x00, 0x00, 0x00,
  0x00, 0x00, 0x00, 0x00, 0x00, 0x00, 0x00, 0x00, 0x00, 0x00, 0x00, 0x00, 0x00, 0x00, 0x00, 0x00,
  0x00, 0x00, 0x00, 0x00, 0x00, 0x00, 0x00, 0x00, 0x00, 0x00, 0x00, 0x00, 0x00, 0x00, 0x00, 0x00,
  0x00, 0x00, 0x00, 0x00, 0x00, 0x00, 0x00, 0x00, 0x00, 0x00, 0x00, 0x00, 0x00, 0x00, 0x00, 0x00,
  0x00, 0x00, 0x00, 0x00, 0x00, 0x00, 0x00, 0x00, 0x00, 0x00, 0x00, 0x00, 0x00, 0x00, 0x00, 0x00,
  0x00, 0x00, 0x00, 0x00, 0x00, 0x00, 0x00, 0x00, 0x00, 0x00, 0x00, 0x00, 0x00, 0x00, 0x00, 0x00,
  0x00, 0x00, 0x00, 0x00, 0x00, 0x00, 0x00, 0x00, 0x00, 0x00, 0x00, 0x00, 0x00, 0x00, 0x00, 0x00,
  0x00, 0x00, 0x00, 0x00, 0x00, 0x00, 0x00, 0x00, 0x00, 0x00, 0x00, 0x00, 0x00, 0x00, 0x00, 0x00,
  0x00, 0x00, 0x00, 0x00, 0x00, 0x00, 0x00, 0x00, 0x00, 0x00, 0x00, 0x00, 0x00, 0x00, 0x00, 0x00,
  0x00, 0x00, 0x00, 0x00, 0x00, 0x00, 0x00, 0x00, 0x00, 0x00, 0x00, 0x00, 0x00, 0x00, 0x00, 0x00,
  0x00, 0x00, 0x00, 0x00, 0x00, 0x00, 0x00, 0x00, 0x00, 0x00, 0x00, 0x00, 0x00, 0x00, 0x00, 0x00,
  0x00, 0x00, 0x00, 0x00, 0x00, 0x00, 0x00, 0x00, 0x00, 0x00, 0x00, 0x00, 0x00, 0x00, 0x00, 0x00,
  0x00, 0x00, 0x00, 0x00, 0x00, 0x00, 0x00, 0x00, 0x00, 0x00, 0x00, 0x00, 0x00, 0x00, 0x00, 0x00,
  0x00, 0x00, 0x00, 0x00, 0x00, 0x00, 0x00, 0x00, 0x00, 0x00, 0x00, 0x00, 0x00, 0x00, 0x00, 0x00,
  0x00, 0x00, 0x00, 0x00, 0x00, 0x00, 0x00, 0x00, 0x00, 0x00, 0x00, 0x00, 0x00, 0x00, 0x00, 0x00,
  0x00, 0x00, 0x00, 0x00, 0x00, 0x00, 0x00, 0x00, 0x00, 0x00, 0x00, 0x00, 0x00, 0x00, 0x00, 0x00,
  0x00, 0x00, 0x00, 0x00, 0x00, 0x00, 0x00, 0x00, 0x00, 0x00, 0x00, 0x00, 0x00, 0x00, 0x00, 0x00,
  0x00, 0x00, 0x00, 0x00, 0x00, 0x00, 0x00, 0x00, 0x00, 0x00, 0x00, 0x00, 0x00, 0x00, 0x00, 0x00,
  0x00, 0x00, 0x00, 0x00, 0x00, 0x00, 0x00, 0x00, 0x00, 0x00, 0x00, 0x00, 0x00, 0x00, 0x00, 0x00,
  0x00, 0x00, 0x00, 0x00, 0x00, 0x00, 0x00, 0x00, 0x00, 0x00, 0x00, 0x00, 0x00, 0x00, 0x00, 0x00,
  0x00, 0x00, 0x00, 0x00, 0x00, 0x00, 0x00, 0x00, 0x00, 0x00, 0x00, 0x00, 0x00, 0x00, 0x00, 0x00,
  0x00, 0x00, 0x00, 0x00, 0x00, 0x00, 0x00, 0x00, 0x00, 0x00, 0x00, 0x00, 0x00, 0x00, 0x00, 0x00,
  0x00, 0x00, 0x00, 0x00, 0x00, 0x00, 0x00, 0x00, 0x00, 0x00, 0x00, 0x00, 0x00, 0x00, 0x00, 0x00,
  0x00, 0x00, 0x00, 0x00, 0x00, 0x00, 0x00, 0x00, 0x00, 0x00, 0x00, 0x00, 0x00, 0x00, 0x00, 0x00,
  0x00, 0x00, 0x00, 0x00, 0x00, 0x00, 0x00, 0x00, 0x00, 0x00, 0x00, 0x00, 0x00, 0x00, 0x00, 0x00,
  0x00, 0x00, 0x00, 0x00, 0x00, 0x00, 0x00, 0x00, 0x00, 0x00, 0x00, 0x00, 0x00, 0x00, 0x00, 0x00,
  0x00, 0x00, 0x00, 0x00, 0x00, 0x00, 0x00, 0x00, 0x00, 0x00, 0x00, 0x00, 0x00, 0x00, 0x00, 0x00,
  0x00, 0x00, 0x00, 0x00, 0x00, 0x00, 0x00, 0x00, 0x00, 0x00, 0x00, 0x00, 0x00, 0x00, 0x00, 0x00,
  0x00, 0x00, 0x00, 0x00, 0x00, 0x00, 0x00, 0x00, 0x00, 0x00, 0x00, 0x00, 0x00, 0x00, 0x00, 0x00,
  0x00, 0x00, 0x00, 0x00, 0x00, 0x00, 0x00, 0x00, 0x00, 0x00, 0x00, 0x00, 0x00, 0x00, 0x00, 0x00,
  0x00, 0x00, 0x00, 0x00, 0x00, 0x00, 0x00, 0x00, 0x00, 0x00, 0x00, 0x00, 0x00, 0x00, 0x00, 0x00,
  0x00, 0x00, 0x00, 0x00, 0x00, 0x00, 0x00, 0x00, 0x00, 0x00, 0x00, 0x00, 0x00, 0x00, 0x00, 0x00,
  0x00, 0x00, 0x00, 0x00, 0x00, 0x00, 0x00, 0x00, 0x00, 0x00, 0x00, 0x00, 0x00, 0x00, 0x00, 0x00,
  0x00, 0x00, 0x00, 0x00, 0x00, 0x00, 0x00, 0x00, 0x00, 0x00, 0x00, 0x00, 0x00, 0x00, 0x00, 0x00,
  0x00, 0x00, 0x00, 0x00, 0x00, 0x00, 0x00, 0x00, 0x00, 0x00, 0x00, 0x00, 0x00, 0x00, 0x00, 0x00,
  0x00, 0x00, 0x00, 0x00, 0x00, 0x00, 0x00, 0x00, 0x00, 0x00, 0x00, 0x00, 0x00, 0x00, 0x00, 0x00,
  0x00, 0x00, 0x00, 0x00, 0x00, 0x00, 0x00, 0x00, 0x00, 0x00, 0x00, 0x00, 0x00, 0x00, 0x00, 0x00,
  0x00, 0x00, 0x00, 0x00, 0x00, 0x00, 0x00, 0x00, 0x00, 0x00, 0x00, 0x00, 0x00, 0x00, 0x00, 0x00,
  0x00, 0x00, 0x00, 0x00, 0x00, 0x00, 0x00, 0x00, 0x00, 0x00, 0x00, 0x00, 0x00, 0x00, 0x00, 0x00,
  0x00, 0x00, 0x00, 0x00, 0x00, 0x00, 0x00, 0x00, 0x00, 0x00, 0x00, 0x00, 0x00, 0x00, 0x00, 0x00,
  0x00, 0x00, 0x00, 0x00, 0x00, 0x00, 0x00, 0x00, 0x00, 0x00, 0x00, 0x00, 0x00, 0x00, 0x00, 0x00,
  0x00, 0x00, 0x00, 0x00, 0x00, 0x00, 0x00, 0x00, 0x00, 0x00, 0x3f, 0xc7, 0xfe, 0x3e, 0x04, 0x7f,
  0xff, 0xff, 0xff, 0xff, 0xff, 0xff, 0xff, 0xff, 0xff, 0xff, 0xff, 0xff, 0xff, 0xff, 0xff, 0xff,
  0xff, 0xff, 0xff, 0xe0, 0x00, 0x00, 0x00, 0x00, 0x3f, 0xc7, 0xfe, 0x3e, 0x06, 0x3f, 0xff, 0xff,
  0xff, 0xff, 0xff, 0xff, 0xff, 0xff, 0xff, 0xff, 0xff, 0xff, 0xff, 0xff, 0xff, 0xff, 0xff, 0xff,
  0xff, 0xe0, 0x00, 0x00, 0x00, 0x00, 0x3f, 0xc0, 0x00, 0x00, 0x07, 0x00, 0x00, 0x00, 0x00, 0x00,
  0x00, 0x00, 0x00, 0x00, 0x00, 0x00, 0x00, 0x00, 0x00, 0x00, 0x00, 0x00, 0x00, 0x00, 0x00, 0x00,
  0x00, 0x00, 0x00, 0x00, 0x3f, 0xc0, 0x00, 0x00, 0x07, 0x80, 0x00, 0x00, 0x00, 0x00, 0x00, 0x00,
  0x00, 0x00, 0x00, 0x00, 0x00, 0x00, 0x00, 0x00, 0x00, 0x00, 0x00, 0x00, 0x00, 0x00, 0x00, 0x00,
  0x00, 0x00, 0x3f, 0xc0, 0x00, 0x00, 0x07, 0xc0, 0x00, 0x00, 0x00, 0x00, 0x00, 0x00, 0x00, 0x00,
  0x00, 0x00, 0x00, 0x00, 0x00, 0x00, 0x00, 0x00, 0x00, 0x00, 0x00, 0x00, 0x00, 0x00, 0x00, 0x00,
  0x3f, 0xc1, 0xfe, 0x3f, 0xc7, 0xe0, 0xfe, 0x3f, 0xff, 0xf8, 0x0f, 0xff, 0xfe, 0x7f, 0xff, 0xf0,
  0x00, 0x03, 0xfc, 0x00, 0xff, 0xff, 0xe3, 0xff, 0xff, 0x80, 0x00, 0x00, 0x00, 0x00, 0x3f, 0xc1,
  0xfe, 0x3f, 0xc7, 0xf0, 0xfe, 0x3f, 0xff, 0xfc, 0x1f, 0xff, 0xfe, 0x7f, 0xff, 0xf8, 0x00, 0x03,
  0xfc, 0x01, 0xff, 0xff, 0xe3, 0xff, 0xff, 0xc0, 0x00, 0x00, 0x00, 0x00, 0x3f, 0xc9, 0xfe, 0x3f,
  0xc7, 0xf8, 0xfe, 0x3f, 0xc7, 0xfe, 0x3f, 0xf0, 0x00, 0x00, 0x0f, 0xfc, 0x00, 0x03, 0xfc, 0x03,
  0xff, 0x1f, 0xe3, 0xfc, 0x7f, 0xe0, 0x00, 0x00, 0x00, 0x00, 0x3f, 0xdd, 0xfe, 0x3f, 0xc7, 0xfc,
  0xfe, 0x3f, 0xc1, 0xfe, 0x3f, 0xc0, 0x00, 0x00, 0x03, 0xfc, 0x00, 0x03, 0xfc, 0x03, 0xfc, 0x1f,
  0xe3, 0xfc, 0x1f, 0xe0, 0x00, 0x00, 0x00, 0x00, 0x3f, 0xff, 0xfe, 0x3f, 0xc7, 0xfe, 0xfe, 0x3f,
  0xc1, 0xfe, 0x3f, 0xc0, 0x00, 0x00, 0x03, 0xfc, 0x00, 0x03, 0xfc, 0x03, 0xfc, 0x1f, 0xe3, 0xfc,
  0x1f, 0xe0, 0x00, 0x00, 0x00, 0x00, 0x3f, 0xff, 0xfe, 0x3f, 0xc7, 0xff, 0xfe, 0x3f, 0xc1, 0xfe,
  0x3f, 0xfe, 0x00, 0x7f, 0xff, 0xfc, 0x00, 0x03, 0xfc, 0x03, 0xff, 0xff, 0xe3, 0xff, 0xff, 0xe0,
  0x00, 0x00, 0x00, 0x00, 0x3f, 0xff, 0xfe, 0x3f, 0xc7, 0xff, 0xfe, 0x3f, 0xc1, 0xfe, 0x3f, 0xfe,
  0x00, 0x7f, 0xff, 0xf8, 0x00, 0x03, 0xfc, 0x03, 0xff, 0xff, 0xe3, 0xff, 0xff, 0xe0, 0x00, 0x00,
  0x00, 0x00, 0x3f, 0xff, 0xfe, 0x3f, 0xc7, 0xff, 0xfe, 0x3f, 0xc1, 0xfe, 0x3f, 0xfe, 0x00, 0x7f,
  0xff, 0x80, 0x00, 0x03, 0xfc, 0x03, 0xff, 0xff, 0xe3, 0xff, 0xff, 0xe0, 0x00, 0x00, 0x00, 0x00,
  0x3f, 0xf7, 0xfe, 0x3f, 0xc7, 0xfb, 0xfe, 0x3f, 0xc1, 0xfe, 0x3f, 0xc0, 0x00, 0x7f, 0x83, 0xf0,
  0x00, 0x03, 0xfc, 0x03, 0xfc, 0x1f, 0xe3, 0xfc, 0x1f, 0xe0, 0x00, 0x00, 0x00, 0x00, 0x3f, 0xe3,
  0xfe, 0x3f, 0xc7, 0xf9, 0xfe, 0x3f, 0xc1, 0xfe, 0x3f, 0xc0, 0x00, 0x7f, 0x83, 0xf8, 0x00, 0x03,
  0xfc, 0x03, 0xfc, 0x1f, 0xe3, 0xfc, 0x1f, 0xe0, 0x00, 0x00, 0x00, 0x00, 0x3f, 0xc1, 0xfe, 0x3f,
  0xc7, 0xf8, 0xfe, 0x3f, 0xff, 0xfe, 0x3f, 0xff, 0xfe, 0x7f, 0x83, 0xfc, 0x00, 0x03, 0xff, 0xe3,
  0xfc, 0x1f, 0xe3, 0xff, 0xff, 0xe0, 0x00, 0x00, 0x00, 0x00, 0x3f, 0x80, 0xfe, 0x3f, 0xc7, 0xf8,
  0x7e, 0x3f, 0xff, 0xfe, 0x3f, 0xff, 0xfe, 0x7f, 0x83, 0xfc, 0x00, 0x03, 0xff, 0xe3, 0xfc, 0x1f,
  0xe3, 0xff, 0xff, 0xe0, 0x00, 0x00, 0x00, 0x00, 0x3f, 0x00, 0x7e, 0x3f, 0xc7, 0xf8, 0x3e, 0x3f,
  0xff, 0xfe, 0x3f, 0xff, 0xfe, 0x7f, 0x83, 0xfc, 0x00, 0x03, 0xff, 0xe3, 0xfc, 0x1f, 0xe3, 0xff,
  0xff, 0xe0, 0x00, 0x00, 0x00, 0x00, 0x00, 0x00, 0x3e, 0x00, 0x00, 0x00, 0x1e, 0x00, 0x00, 0x00,
  0x00, 0x00, 0x00, 0x00, 0x00, 0x00, 0x00, 0x00, 0x00, 0x00, 0x00, 0x1f, 0xe0, 0x00, 0x00, 0x00,
  0x00, 0x00, 0x00, 0x00, 0x00, 0x00, 0x1e, 0x00, 0x00, 0x00, 0x0e, 0x00, 0x00, 0x00, 0x00, 0x00,
  0x00, 0x00, 0x00, 0x00, 0x00, 0x00, 0x00, 0x00, 0x00, 0x1f, 0xe0, 0x00, 0x00, 0x00, 0x00, 0x00,
  0x00, 0x00, 0x00, 0x00, 0x0e, 0x00, 0x00, 0x00, 0x06, 0x00, 0x00, 0x00, 0x00, 0x00, 0x00, 0x00,
  0x00, 0x00, 0x00, 0x00, 0x00, 0x00, 0x00, 0x1f, 0xe0, 0x00, 0x00, 0x00, 0x00, 0x00, 0x00, 0x00,
  0x3f, 0xff, 0xe6, 0x3f, 0xff, 0xff, 0xe2, 0x3f, 0xff, 0xff, 0xff, 0xff, 0xff, 0xff, 0xff, 0xff,
  0xff, 0xff, 0xff, 0xff, 0xff, 0x9f, 0xe3, 0xff, 0xff, 0xfc, 0x00, 0x00, 0x00, 0x00, 0x3f, 0xff,
  0xf2, 0x3f, 0xff, 0xff, 0xf0, 0x3f, 0xff, 0xff, 0xff, 0xff, 0xff, 0xff, 0xff, 0xff, 0xff, 0xff,
  0xff, 0xff, 0xff, 0x9f, 0xe3, 0xff, 0xff, 0xfc, 0x00, 0x00, 0x00, 0x00, 0x00, 0x00, 0x00, 0x00,
  0x00, 0x00, 0x00, 0x00, 0x00, 0x00, 0x00, 0x00, 0x00, 0x00, 0x00, 0x00, 0x00, 0x00, 0x00, 0x00,
  0x00, 0x00, 0x00, 0x00, 0x00, 0x00, 0x00, 0x00, 0x00, 0x00, 0x00, 0x00, 0x00, 0x00, 0x00, 0x00,
  0x00, 0x00, 0x00, 0x00, 0x00, 0x00, 0x00, 0x00, 0x00, 0x00, 0x00, 0x00, 0x00, 0x00, 0x00, 0x00,
  0x00, 0x00, 0x00, 0x00, 0x00, 0x00, 0x00, 0x00, 0x00, 0x00, 0x00, 0x00, 0x00, 0x00, 0x00, 0x00,
  0x00, 0x00, 0x00, 0x00, 0x00, 0x00, 0x00, 0x00, 0x00, 0x00, 0x00, 0x00, 0x00, 0x00, 0x00, 0x00,
  0x00, 0x00, 0x00, 0x00, 0x00, 0x00, 0x00, 0x00, 0x00, 0x00, 0x00, 0x00, 0x00, 0x00, 0x00, 0x00,
  0x00, 0x00, 0x00, 0x00, 0x00, 0x00, 0x00, 0x00, 0x00, 0x00, 0x00, 0x00, 0x00, 0x00, 0x00, 0x00,
  0x00, 0x00, 0x00, 0x00, 0x00, 0x00, 0x00, 0x00, 0x00, 0x00, 0x00, 0x00, 0x00, 0x00, 0x00, 0x00,
  0x00, 0x00, 0x00, 0x00, 0x00, 0x00, 0x00, 0x00, 0x00, 0x00, 0x00, 0x00, 0x00, 0x00, 0x00, 0x00,
  0x00, 0x00, 0x00, 0x00, 0x00, 0x00, 0x00, 0x00, 0x00, 0x00, 0x00, 0x00, 0x00, 0x00, 0x00, 0x00,
  0x00, 0x00, 0x00, 0x00, 0x00, 0x00, 0x00, 0x00, 0x00, 0x00, 0x00, 0x00, 0x00, 0x00, 0x00, 0x00,
  0x00, 0x00, 0x00, 0x00, 0x00, 0x00, 0x00, 0x00, 0x00, 0x00, 0x00, 0x00, 0x00, 0x00, 0x00, 0x00,
  0x00, 0x00, 0x00, 0x00, 0x00, 0x00, 0x00, 0x00, 0x00, 0x00, 0x00, 0x00, 0x00, 0x00, 0x00, 0x00,
  0x00, 0x00, 0x00, 0x00, 0x00, 0x00, 0x00, 0x00, 0x00, 0x00, 0x00, 0x00, 0x00, 0x00, 0x00, 0x00,
  0x00, 0x00, 0x00, 0x00, 0x00, 0x00, 0x00, 0x00, 0x00, 0x00, 0x00, 0x00, 0x00, 0x00, 0x00, 0x00,
  0x00, 0x00, 0x00, 0x00, 0x00, 0x00, 0x00, 0x00, 0x00, 0x00, 0x00, 0x00, 0x00, 0x00, 0x00, 0x00,
  0x00, 0x00, 0x00, 0x00, 0x00, 0x00, 0x00, 0x00, 0x00, 0x00, 0x00, 0x00, 0x00, 0x00, 0x00, 0x00,
  0x00, 0x00, 0x00, 0x00, 0x00, 0x00, 0x00, 0x00, 0x00, 0x00, 0x00, 0x00, 0x00, 0x00, 0x00, 0x00,
  0x00, 0x00, 0x00, 0x00, 0x00, 0x00, 0x00, 0x00, 0x00, 0x00, 0x00, 0x00, 0x00, 0x00, 0x00, 0x00,
  0x00, 0x00, 0x00, 0x00, 0x00, 0x00, 0x00, 0x00, 0x00, 0x00, 0x00, 0x00, 0x00, 0x00, 0x00, 0x00,
  0x00, 0x00, 0x00, 0x00, 0x00, 0x00, 0x00, 0x00, 0x00, 0x00, 0x00, 0x00, 0x00, 0x00, 0x00, 0x00,
  0x00, 0x00, 0x00, 0x00, 0x00, 0x00, 0x00, 0x00, 0x00, 0x00, 0x00, 0x00, 0x00, 0x00, 0x00, 0x00,
  0x00, 0x00, 0x00, 0x00, 0x00, 0x00, 0x00, 0x00, 0x00, 0x00, 0x00, 0x00, 0x00, 0x00, 0x00, 0x00,
  0x00, 0x00, 0x00, 0x00, 0x00, 0x00, 0x00, 0x00, 0x00, 0x00, 0x00, 0x00, 0x00, 0x00, 0x00, 0x00,
  0x00, 0x00, 0x00, 0x00, 0x00, 0x00, 0x00, 0x00, 0x00, 0x00, 0x00, 0x00, 0x00, 0x00, 0x00, 0x00,
  0x00, 0x00, 0x00, 0x00, 0x00, 0x00, 0x00, 0x00, 0x00, 0x00, 0x00, 0x00, 0x00, 0x00, 0x00, 0x00,
  0x00, 0x00, 0x00, 0x00, 0x00, 0x00, 0x00, 0x00, 0x00, 0x00, 0x00, 0x00, 0x00, 0x00, 0x00, 0x00,
  0x00, 0x00, 0x00, 0x00, 0x00, 0x00, 0x00, 0x00, 0x00, 0x00, 0x00, 0x00, 0x00, 0x00, 0x00, 0x00,
  0x00, 0x00, 0x00, 0x00, 0x00, 0x00, 0x00, 0x00, 0x00, 0x00, 0x00, 0x00, 0x00, 0x00, 0x00, 0x00,
  0x00, 0x00, 0x00, 0x00, 0x00, 0x00, 0x00, 0x00, 0x00, 0x00, 0x00, 0x00, 0x00, 0x00, 0x00, 0x00,
  0x00, 0x00, 0x00, 0x00, 0x00, 0x00, 0x00, 0x00, 0x00, 0x00, 0x00, 0x00, 0x00, 0x00, 0x00, 0x00,
  0x00, 0x00, 0x00, 0x00, 0x00, 0x00, 0x00, 0x00, 0x00, 0x00, 0x00, 0x00, 0x00, 0x00, 0x00, 0x00,
  0x00, 0x00, 0x00, 0x00, 0x00, 0x00, 0x00, 0x00, 0x00, 0x00, 0x00, 0x00, 0x00, 0x00, 0x00, 0x00,
  0x00, 0x00, 0x00, 0x00, 0x00, 0x00
};
// 'main_logo', 240x60px
const unsigned char main_logo[] PROGMEM = {
  0x00, 0x00, 0x00, 0x00, 0x00, 0x00, 0x00, 0x00, 0x00, 0x00, 0x00, 0x00, 0x00, 0x00, 0x00, 0x00,
  0x00, 0x00, 0x00, 0x00, 0x00, 0x00, 0x00, 0x00, 0x00, 0x00, 0x00, 0x00, 0x00, 0x00, 0x00, 0x00,
  0x00, 0x00, 0x00, 0x00, 0x00, 0x00, 0x00, 0x00, 0x00, 0x00, 0x00, 0x00, 0x00, 0x00, 0x00, 0x00,
  0x00, 0x00, 0x00, 0x00, 0x00, 0x00, 0x00, 0x00, 0x00, 0x00, 0x00, 0x00, 0x00, 0x00, 0x00, 0x00,
  0x00, 0x00, 0x00, 0x00, 0x00, 0x00, 0x00, 0x00, 0x00, 0x00, 0x00, 0x00, 0x00, 0x00, 0x00, 0x00,
  0x00, 0x00, 0x00, 0x00, 0x00, 0x00, 0x00, 0x00, 0x00, 0x00, 0x00, 0x00, 0x00, 0x00, 0x00, 0x00,
  0x00, 0x00, 0x00, 0x00, 0x00, 0x00, 0x00, 0x00, 0x00, 0x00, 0x00, 0x00, 0x00, 0x00, 0x00, 0x00,
  0x00, 0x00, 0x00, 0x00, 0x00, 0x00, 0x00, 0x00, 0x00, 0x00, 0x00, 0x00, 0x00, 0x00, 0x00, 0x00,
  0x00, 0x00, 0x00, 0x00, 0x00, 0x00, 0x00, 0x00, 0x00, 0x00, 0x00, 0x00, 0x00, 0x00, 0x00, 0x00,
  0x00, 0x00, 0x00, 0x00, 0x00, 0x00, 0x00, 0x00, 0x00, 0x00, 0x00, 0x00, 0x00, 0x00, 0x00, 0x00,
  0x00, 0x00, 0x00, 0x00, 0x00, 0x00, 0x00, 0x00, 0x00, 0x00, 0x00, 0x00, 0x00, 0x00, 0x00, 0x00,
  0x00, 0x00, 0x00, 0x00, 0x00, 0x00, 0x00, 0x00, 0x00, 0x00, 0x00, 0x00, 0x00, 0x00, 0x00, 0x00,
  0x00, 0x00, 0x00, 0x00, 0x00, 0x00, 0x00, 0x00, 0x00, 0x00, 0x00, 0x00, 0x00, 0x00, 0x00, 0x00,
  0x00, 0x00, 0x00, 0x00, 0x00, 0x00, 0x00, 0x00, 0x00, 0x00, 0x00, 0x00, 0x00, 0x00, 0x00, 0x00,
  0x00, 0x00, 0x00, 0x00, 0x00, 0x00, 0x00, 0x00, 0x00, 0x00, 0x00, 0x00, 0x00, 0x00, 0x00, 0x00,
  0x00, 0x00, 0x00, 0x00, 0x00, 0x00, 0x00, 0x7c, 0x00, 0x07, 0xc0, 0x03, 0xff, 0x80, 0x00, 0x00,
  0x1f, 0x00, 0x03, 0xe0, 0x7f, 0xff, 0xc0, 0x00, 0x00, 0x00, 0x00, 0x00, 0x00, 0x00, 0x00, 0x00,
  0x00, 0x00, 0x00, 0x00, 0x00, 0x7c, 0x00, 0x07, 0xc0, 0x1f, 0xff, 0xf0, 0x00, 0x00, 0x1f, 0x00,
  0x03, 0xe0, 0x7f, 0xff, 0xf8, 0x00, 0x00, 0x00, 0x00, 0x00, 0x00, 0x00, 0x00, 0x00, 0x00, 0x00,
  0x00, 0x00, 0x00, 0x7c, 0x00, 0x07, 0xc0, 0x3f, 0xff, 0xf8, 0x00, 0x00, 0x1f, 0x00, 0x03, 0xe0,
  0x7f, 0xff, 0xfc, 0x00, 0x00, 0x00, 0x00, 0x00, 0x00, 0x00, 0x00, 0x00, 0x00, 0x00, 0x00, 0x00,
  0x00, 0x7c, 0x00, 0x07, 0xc0, 0x7f, 0xff, 0xfc, 0x00, 0x00, 0x1f, 0x00, 0x03, 0xe0, 0x7f, 0xff,
  0xfe, 0x00, 0x00, 0x00, 0x00, 0x00, 0x00, 0x00, 0x00, 0x00, 0x00, 0x00, 0x00, 0x00, 0x00, 0x7c,
  0x00, 0x07, 0xc0, 0x7f, 0xff, 0xfc, 0x00, 0x00, 0x1f, 0x00, 0x03, 0xe0, 0x7f, 0xff, 0xfe, 0x00,
  0x00, 0x00, 0x00, 0x00, 0x00, 0x00, 0x00, 0x00, 0x00, 0x00, 0x00, 0x00, 0x00, 0x7c, 0x00, 0x07,
  0xc0, 0xfc, 0x00, 0x7e, 0x00, 0x00, 0x1f, 0x00, 0x03, 0xe0, 0x7c, 0x00, 0x3f, 0x00, 0x00, 0x00,
  0x00, 0x00, 0x00, 0x00, 0x00, 0x00, 0x00, 0x00, 0x00, 0x00, 0x00, 0x7c, 0x00, 0x07, 0xc0, 0xf8,
  0x00, 0x3e, 0x00, 0x00, 0x1f, 0x00, 0x03, 0xe0, 0x7c, 0x00, 0x1f, 0x00, 0x00, 0x00, 0x00, 0x00,
  0x00, 0x00, 0x00, 0x00, 0x00, 0x00, 0x00, 0x00, 0x00, 0x7c, 0x00, 0x07, 0xc0, 0xf8, 0x00, 0x3e,
  0x00, 0x00, 0x1f, 0x00, 0x03, 0xe0, 0x7c, 0x00, 0x1f, 0x00, 0x00, 0x00, 0x00, 0x00, 0x00, 0x00,
  0x00, 0x00, 0x00, 0x00, 0x00, 0x00, 0x00, 0x7c, 0x00, 0x07, 0xc0, 0xf8, 0x00, 0x3e, 0x00, 0x00,
  0x1f, 0x00, 0x03, 0xe0, 0x7c, 0x00, 0x1f, 0x00, 0x00, 0x00, 0x00, 0x00, 0x00, 0x00, 0x00, 0x00,
  0x00, 0x00, 0x00, 0x00, 0x00, 0x7c, 0x00, 0x07, 0xc0, 0xf8, 0x00, 0x3e, 0x00, 0x00, 0x1f, 0x00,
  0x03, 0xe0, 0x7c, 0x00, 0x1f, 0x00, 0x00, 0x00, 0x00, 0x00, 0x00, 0x00, 0x00, 0x00, 0x00, 0x00,
  0x00, 0x00, 0x00, 0x7c, 0x00, 0x07, 0xc0, 0xf8, 0x00, 0x3e, 0x00, 0x00, 0x1f, 0x00, 0x03, 0xe0,
  0x7c, 0x00, 0x1f, 0x00, 0x00, 0x00, 0x00, 0x00, 0x00, 0x00, 0x00, 0x00, 0x00, 0x00, 0x00, 0x00,
  0x00, 0x7c, 0x00, 0x07, 0xc0, 0xf8, 0x00, 0x3e, 0x00, 0x00, 0x1f, 0x00, 0x03, 0xe0, 0x7c, 0x00,
  0x1f, 0x00, 0x00, 0x00, 0x00, 0x00, 0x00, 0x00, 0x00, 0x00, 0x00, 0x00, 0x00, 0x00, 0x00, 0x7c,
  0x00, 0x07, 0xc0, 0xf8, 0x00, 0x3e, 0x00, 0x00, 0x1f, 0x00, 0x03, 0xe0, 0x7c, 0x00, 0x1f, 0x00,
  0x00, 0x00, 0x00, 0x00, 0x00, 0x00, 0x00, 0x00, 0x00, 0x00, 0x00, 0x00, 0x00, 0x7c, 0x00, 0x07,
  0xc0, 0xf8, 0x00, 0x3e, 0x00, 0x00, 0x1f, 0x00, 0x03, 0xe0, 0x7c, 0x00, 0x1f, 0x00, 0x00, 0x00,
  0x00, 0x00, 0x00, 0x00, 0x00, 0x00, 0x00, 0x00, 0x00, 0x00, 0x00, 0x7c, 0x00, 0x07, 0xc0, 0xf8,
  0x00, 0x3e, 0x00, 0x00, 0x1f, 0x00, 0x03, 0xe0, 0x7c, 0x00, 0x1f, 0x00, 0x00, 0x00, 0x00, 0x00,
  0x00, 0x00, 0x00, 0x00, 0x00, 0x00, 0x00, 0x00, 0x00, 0x7c, 0x00, 0x07, 0xc0, 0xf8, 0x00, 0x3e,
  0x00, 0x00, 0x1f, 0x00, 0x03, 0xe0, 0x7c, 0x00, 0x1f, 0x00, 0x00, 0x00, 0x00, 0x00, 0x00, 0x00,
  0x00, 0x00, 0x00, 0x00, 0x00, 0x00, 0x00, 0x7c, 0x00, 0x07, 0xc0, 0xf8, 0x00, 0x3e, 0x00, 0x00,
  0x1f, 0x00, 0x03, 0xe0, 0x7c, 0x00, 0x1f, 0x00, 0x00, 0x00, 0x00, 0x00, 0x00, 0x00, 0x00, 0x00,
  0x00, 0x00, 0x00, 0x00, 0x00, 0x7c, 0x00, 0x07, 0xc0, 0xf8, 0x00, 0x3e, 0x00, 0x00, 0x1f, 0x00,
  0x03, 0xe0, 0x7c, 0x00, 0x1f, 0x00, 0x00, 0x00, 0x00, 0x00, 0x00, 0x00, 0x00, 0x00, 0x00, 0x00,
  0x00, 0x00, 0x00, 0x7c, 0x00, 0x07, 0xc0, 0xf8, 0x00, 0x3e, 0x00, 0x00, 0x1f, 0x00, 0x03, 0xe0,
  0x7c, 0x00, 0x1f, 0x00, 0x00, 0x00, 0x00, 0x00, 0x00, 0x00, 0x00, 0x00, 0x00, 0x00, 0x00, 0x00,
  0x00, 0x7c, 0x00, 0x07, 0xc0, 0xf8, 0x00, 0x3e, 0x00, 0x00, 0x1f, 0x00, 0x03, 0xe0, 0x7c, 0x00,
  0x1f, 0x00, 0x00, 0x00, 0x00, 0x00, 0x00, 0x00, 0x00, 0x00, 0x00, 0x00, 0x00, 0x00, 0x00, 0x7c,
  0x00, 0x07, 0xc0, 0xf8, 0x00, 0x3e, 0x00, 0x00, 0x1f, 0x00, 0x03, 0xe0, 0x7c, 0x00, 0x1f, 0x00,
  0x00, 0x00, 0x00, 0x00, 0x00, 0x00, 0x00, 0x00, 0x00, 0x00, 0x00, 0x00, 0x00, 0x7c, 0x00, 0x07,
  0xc0, 0xf8, 0x00, 0x3e, 0x00, 0x00, 0x1f, 0x00, 0x03, 0xe0, 0x7c, 0x00, 0x1f, 0x00, 0x00, 0x00,
  0x00, 0x00, 0x00, 0x00, 0x00, 0x00, 0x00, 0x00, 0x00, 0x00, 0x00, 0x7c, 0x00, 0x07, 0xc0, 0xf8,
  0x00, 0x3e, 0x00, 0x00, 0x1f, 0x00, 0x03, 0xe0, 0x7c, 0x00, 0x1f, 0x00, 0x00, 0x00, 0x00, 0x00,
  0x00, 0x00, 0x00, 0x00, 0x00, 0x00, 0x00, 0x00, 0x00, 0x7c, 0x00, 0x07, 0xc0, 0xf8, 0x00, 0x3e,
  0x00, 0x00, 0xff, 0xff, 0xff, 0xe0, 0x7c, 0x00, 0x1f, 0x00, 0x00, 0x00, 0x00, 0x00, 0x00, 0x00,
  0x00, 0x00, 0x00, 0x00, 0x00, 0x00, 0x00, 0x7c, 0x00, 0x07, 0xc0, 0xf8, 0x00, 0x3e, 0x00, 0x00,
  0xff, 0xff, 0xff, 0xe0, 0x7c, 0x00, 0x1f, 0x00, 0x00, 0x00, 0x00, 0x00, 0x00, 0x00, 0x00, 0x00,
  0x00, 0x00, 0x00, 0x00, 0x00, 0x7c, 0x00, 0x07, 0xc0, 0xf8, 0x00, 0x3e, 0x00, 0x00, 0xff, 0xff,
  0xff, 0xe0, 0x7c, 0x00, 0x1f, 0x00, 0x00, 0x00, 0x00, 0x00, 0x00, 0x00, 0x00, 0x00, 0x00, 0x00,
  0x00, 0x00, 0x00, 0x7c, 0x00, 0x07, 0xc0, 0xf8, 0x00, 0x3e, 0x00, 0x00, 0xff, 0xff, 0xff, 0xe0,
  0x7c, 0x00, 0x1f, 0x00, 0x00, 0x00, 0x00, 0x00, 0x00, 0x00, 0x00, 0x00, 0x00, 0x00, 0x00, 0x00,
  0x00, 0x7c, 0x00, 0x07, 0xc0, 0xf8, 0x00, 0x3e, 0x00, 0x00, 0xff, 0xff, 0xff, 0xe0, 0x7c, 0x00,
  0x1f, 0x00, 0x00, 0x00, 0x00, 0x00, 0x00, 0x00, 0x00, 0x00, 0x00, 0x00, 0x00, 0x00, 0x00, 0x7c,
  0x00, 0x07, 0xc0, 0xf8, 0x00, 0x3e, 0x00, 0x00, 0x1f, 0x00, 0x03, 0xe0, 0x7c, 0x00, 0x1f, 0x00,
  0x00, 0x00, 0x00, 0x00, 0x00, 0x00, 0x00, 0x00, 0x00, 0x00, 0x00, 0x00, 0x00, 0x7c, 0x00, 0x07,
  0xc0, 0xf8, 0x00, 0x3e, 0x00, 0x00, 0x1f, 0x00, 0x03, 0xe0, 0x7c, 0x00, 0x1f, 0x00, 0x00, 0x00,
  0x00, 0x00, 0x00, 0x00, 0x00, 0x00, 0x00, 0x00, 0x00, 0x00, 0x00, 0x7c, 0x00, 0x07, 0xc0, 0xf8,
  0x00, 0x3e, 0x00, 0x00, 0x1f, 0x00, 0x03, 0xe0, 0x7c, 0x00, 0x1f, 0x00, 0x00, 0x00, 0x00, 0x00,
  0x00, 0x00, 0x00, 0x00, 0x00, 0x00, 0x00, 0x00, 0x00, 0x7c, 0x00, 0x07, 0xc0, 0xf8, 0x00, 0x3e,
  0x00, 0x00, 0x1f, 0x00, 0x03, 0xe0, 0x7c, 0x00, 0x1f, 0x00, 0x00, 0x00, 0x00, 0x00, 0x00, 0x00,
  0x00, 0x00, 0x00, 0x00, 0x00, 0x00, 0x00, 0x7c, 0x00, 0x07, 0xc0, 0xf8, 0x00, 0x3e, 0x00, 0x00,
  0x1f, 0x00, 0x03, 0xe0, 0x7c, 0x00, 0x1f, 0x00, 0x00, 0x00, 0x00, 0x00, 0x00, 0x00, 0x00, 0x00,
  0x00, 0x00, 0x00, 0x00, 0x00, 0x7c, 0x00, 0x07, 0xc0, 0xfc, 0x00, 0x7e, 0x00, 0x00, 0x1f, 0x00,
  0x03, 0xe0, 0x7c, 0x00, 0x3f, 0x00, 0x00, 0x00, 0x00, 0x00, 0x00, 0x00, 0x00, 0x00, 0x00, 0x00,
  0x00, 0x00, 0x00, 0x7f, 0xff, 0x07, 0xc0, 0x7f, 0xff, 0xfc, 0x00, 0x00, 0x1f, 0x00, 0x03, 0xe0,
  0x7f, 0xff, 0xfe, 0x00, 0x00, 0x00, 0x00, 0x00, 0x00, 0x00, 0x00, 0x00, 0x00, 0x00, 0x00, 0x00,
  0x00, 0x7f, 0xff, 0x07, 0xc0, 0x7f, 0xff, 0xfc, 0x00, 0x00, 0x1f, 0x00, 0x03, 0xe0, 0x7f, 0xff,
  0xfe, 0x00, 0x00, 0x00, 0x00, 0x00, 0x00, 0x00, 0x00, 0x00, 0x00, 0x00, 0x00, 0x00, 0x00, 0x7f,
  0xff, 0x07, 0xc0, 0x3f, 0xff, 0xf8, 0x00, 0x00, 0x1f, 0x00, 0x03, 0xe0, 0x7f, 0xff, 0xfc, 0x00,
  0x00, 0x00, 0x00, 0x00, 0x00, 0x00, 0x00, 0x00, 0x00, 0x00, 0x00, 0x00, 0x00, 0x7f, 0xff, 0x07,
  0xc0, 0x1f, 0xff, 0xf0, 0x00, 0x00, 0x1f, 0x00, 0x03, 0xe0, 0x7f, 0xff, 0xf8, 0x00, 0x00, 0x00,
  0x00, 0x00, 0x00, 0x00, 0x00, 0x00, 0x00, 0x00, 0x00, 0x00, 0x00, 0x7f, 0xff, 0x07, 0xc0, 0x03,
  0xff, 0x80, 0x00, 0x00, 0x1f, 0x00, 0x03, 0xe0, 0x7f, 0xff, 0xc0, 0x00, 0x00, 0x00, 0x00, 0x00,
  0x00, 0x00, 0x00, 0x00, 0x00, 0x00, 0x00, 0x00, 0x00, 0x00, 0x00, 0x00, 0x00, 0x00, 0x3f, 0x80,
  0x00, 0x00, 0x00, 0x00, 0x00, 0x00, 0x00, 0x00, 0x00, 0x00, 0x00, 0x00, 0x00, 0x00, 0x00, 0x00,
  0x00, 0x00, 0x00, 0x00, 0x00, 0x00, 0x00, 0x00, 0x00, 0x00, 0x00, 0x00, 0x1f, 0xc0, 0x00, 0x00,
  0x00, 0x00, 0x00, 0x00, 0x00, 0x00, 0x00, 0x00, 0x00, 0x00, 0x00, 0x00, 0x00, 0x00, 0x00, 0x00,
  0x00, 0x00, 0x00, 0x00, 0x00, 0x00, 0x00, 0x00, 0x00, 0x00, 0x0f, 0xe0, 0x00, 0x00, 0x00, 0x00,
  0x00, 0x00, 0x00, 0x00, 0x00, 0x00, 0x00, 0x00, 0x00, 0x00, 0x00, 0x00, 0x00, 0x00, 0x00, 0x00,
  0x00, 0x00, 0x00, 0x00, 0x00, 0x00, 0x00, 0x00, 0x07, 0x80, 0x00, 0x00, 0x00, 0x00, 0x00, 0x00,
  0x00, 0x00, 0x00, 0x00, 0x00, 0x00, 0x00, 0x00, 0x00, 0x00, 0x00, 0x00, 0x00, 0x00, 0x00, 0x00,
  0x00, 0x00, 0x00, 0x00, 0x00, 0x00, 0x03, 0x80, 0x00, 0x00, 0x00, 0x00, 0x00, 0x00, 0x00, 0x00,
  0x00, 0x00, 0x00, 0x00, 0x00, 0x00, 0x00, 0x00, 0x00, 0x00, 0x00, 0x00, 0x00, 0x00, 0x00, 0x00,
  0x00, 0x00, 0x00, 0x00, 0x01, 0x00, 0x00, 0x00, 0x00, 0x00, 0x00, 0x00, 0x00, 0x00, 0x00, 0x00,
  0x00, 0x00, 0x00, 0x00, 0x00, 0x00, 0x00, 0x00, 0x00, 0x00, 0x00, 0x00, 0x00, 0x00, 0x00, 0x00,
  0x00, 0x00, 0x00, 0x00, 0x00, 0x00, 0x00, 0x00, 0x00, 0x00, 0x00, 0x00, 0x00, 0x00, 0x00, 0x00,
  0x00, 0x00, 0x00, 0x00, 0x00, 0x00, 0x00, 0x00, 0x00, 0x00, 0x00, 0x00, 0x00, 0x00, 0x00, 0x00,
  0x00, 0x00, 0x00, 0x00, 0x00, 0x00, 0x00, 0x00, 0x00, 0x00, 0x00, 0x00, 0x00, 0x00, 0x00, 0x00,
  0x00, 0x00, 0x00, 0x00, 0x00, 0x00, 0x00, 0x00, 0x00, 0x00, 0x00, 0x00, 0x00, 0x00, 0x00, 0x00,
  0x00, 0x00, 0x00, 0x00, 0x00, 0x00, 0x00, 0x00, 0x00, 0x00, 0x00, 0x00, 0x00, 0x00, 0x00, 0x00,
  0x00, 0x00, 0x00, 0x00, 0x00, 0x00, 0x00, 0x00, 0x00, 0x00, 0x00, 0x00, 0x00, 0x00, 0x00, 0x00,
  0x00, 0x00, 0x00, 0x00, 0x00, 0x00, 0x00, 0x00, 0x00, 0x00, 0x00, 0x00, 0x00, 0x00, 0x00, 0x00,
  0x00, 0x00, 0x00, 0x00, 0x00, 0x00, 0x00, 0x00, 0x00, 0x00, 0x00, 0x00, 0x00, 0x00, 0x00, 0x00,
  0x00, 0x00, 0x00, 0x00, 0x00, 0x00, 0x00, 0x00, 0x00, 0x00, 0x00, 0x00, 0x00, 0x00, 0x00, 0x00,
  0x00, 0x00, 0x00, 0x00, 0x00, 0x00, 0x00, 0x00, 0x00, 0x00, 0x00, 0x00, 0x00, 0x00, 0x00, 0x00,
  0x00, 0x00, 0x00, 0x00, 0x00, 0x00, 0x00, 0x00, 0x00, 0x00, 0x00, 0x00, 0x00, 0x00, 0x00, 0x00,
  0x00, 0x00, 0x00, 0x00, 0x00, 0x00, 0x00, 0x00, 0x00, 0x00, 0x00, 0x00, 0x00, 0x00, 0x00, 0x00,
  0x00, 0x00, 0x00, 0x00, 0x00, 0x00, 0x00, 0x00
};


const unsigned char settings_icon[] PROGMEM = {
  // 'Very-Basic-Settings-Filled-icon-2, 32x32px
  0x00, 0x03, 0xc0, 0x00, 0x00, 0x07, 0xe0, 0x00, 0x00, 0x07, 0xe0, 0x00, 0x02, 0x07, 0xe0, 0x40,
  0x07, 0x07, 0xe0, 0xe0, 0x0f, 0xdf, 0xfb, 0xf0, 0x1f, 0xff, 0xff, 0xf8, 0x0f, 0xff, 0xff, 0xf0,
  0x07, 0xff, 0xff, 0xe0, 0x07, 0xff, 0xff, 0xe0, 0x03, 0xff, 0xff, 0xc0, 0x07, 0xfe, 0x7f, 0xe0,
  0x07, 0xf8, 0x1f, 0xe0, 0x7f, 0xf0, 0x0f, 0xfe, 0xff, 0xf0, 0x0f, 0xff, 0xff, 0xe0, 0x07, 0xff,
  0xff, 0xe0, 0x07, 0xff, 0xff, 0xf0, 0x0f, 0xff, 0x7f, 0xf0, 0x0f, 0xfe, 0x07, 0xf8, 0x1f, 0xe0,
  0x07, 0xfe, 0x7f, 0xe0, 0x03, 0xff, 0xff, 0xc0, 0x07, 0xff, 0xff, 0xe0, 0x07, 0xff, 0xff, 0xe0,
  0x0f, 0xff, 0xff, 0xf0, 0x1f, 0xff, 0xff, 0xf8, 0x0f, 0xdf, 0xfb, 0xf0, 0x07, 0x07, 0xe0, 0xe0,
  0x02, 0x07, 0xe0, 0x40, 0x00, 0x07, 0xe0, 0x00, 0x00, 0x07, 0xe0, 0x00, 0x00, 0x03, 0xc0, 0x00
};

const unsigned char settings_icon2[] PROGMEM = {
  // 'Settings-icon-3, 32x32px
  0x00, 0x00, 0x00, 0x00, 0x00, 0x7f, 0xfe, 0x00, 0x01, 0xff, 0xff, 0x80, 0x07, 0xff, 0xff, 0xe0,
  0x0f, 0xff, 0xff, 0xf0, 0x1f, 0xff, 0xff, 0xf8, 0x1f, 0xc7, 0xff, 0xf8, 0x3f, 0xc3, 0xff, 0xfc,
  0x3f, 0xe3, 0xff, 0xfc, 0x7f, 0xe3, 0xff, 0xfe, 0x7c, 0xc3, 0xff, 0xfe, 0x7c, 0x03, 0xff, 0xfe,
  0x7e, 0x01, 0xff, 0xfe, 0x7f, 0x00, 0xff, 0xfe, 0x7f, 0xf8, 0x7f, 0xfe, 0x7f, 0xfc, 0x3f, 0xfe,
  0x7f, 0xfe, 0x1f, 0xfe, 0x7f, 0xff, 0x0f, 0xfe, 0x7f, 0xff, 0x80, 0x7e, 0x7f, 0xff, 0xc0, 0x3e,
  0x7f, 0xff, 0xc0, 0x3e, 0x7f, 0xff, 0xc3, 0x3e, 0x7f, 0xff, 0xc7, 0xfe, 0x3f, 0xff, 0xc7, 0xfc,
  0x3f, 0xff, 0xe3, 0xfc, 0x1f, 0xff, 0xf3, 0xf8, 0x1f, 0xff, 0xff, 0xf8, 0x0f, 0xff, 0xff, 0xf0,
  0x07, 0xff, 0xff, 0xe0, 0x01, 0xff, 0xff, 0x80, 0x00, 0x7f, 0xfe, 0x00, 0x00, 0x00, 0x00, 0x00
};

// 'LIQ PARTI Winder Lab Logo', 240x123px
const unsigned char PARTI_LOGOLIQ_PARTI_Winder_Lab_Logo[] PROGMEM = {
  0x00, 0x00, 0x00, 0x00, 0x00, 0x00, 0x00, 0x00, 0x00, 0x00, 0x00, 0x00, 0x00, 0x00, 0x00, 0x00,
  0x00, 0x00, 0x00, 0x00, 0x00, 0x00, 0x00, 0x00, 0x00, 0x00, 0x00, 0x00, 0x00, 0x00, 0x00, 0x00,
  0x00, 0x00, 0x00, 0x00, 0x00, 0x00, 0x00, 0x00, 0x00, 0x00, 0x00, 0x00, 0x00, 0x00, 0x00, 0x00,
  0x00, 0x00, 0x00, 0x00, 0x00, 0x00, 0x00, 0x00, 0x00, 0x00, 0x00, 0x00, 0x00, 0x00, 0x00, 0x00,
  0x00, 0x00, 0x00, 0x00, 0x00, 0x00, 0x00, 0x00, 0x00, 0x00, 0x00, 0x00, 0x00, 0x00, 0x00, 0x00,
  0x00, 0x00, 0x00, 0x00, 0x00, 0x00, 0x20, 0x00, 0x00, 0x00, 0x00, 0x00, 0x00, 0x00, 0x00, 0x00,
  0x00, 0x00, 0x00, 0x00, 0x00, 0x00, 0x03, 0xff, 0xff, 0xff, 0xff, 0xff, 0xff, 0xff, 0xff, 0xff,
  0xff, 0xff, 0xff, 0xff, 0xe0, 0x00, 0x00, 0x00, 0x00, 0x00, 0x00, 0x00, 0x00, 0x00, 0x00, 0x00,
  0x00, 0x00, 0x00, 0x00, 0x03, 0xff, 0xff, 0xff, 0xff, 0xff, 0xff, 0xff, 0xff, 0xff, 0xff, 0xff,
  0xff, 0xff, 0xe0, 0x00, 0x00, 0x00, 0x00, 0x00, 0x00, 0x00, 0x00, 0x00, 0x00, 0x00, 0x00, 0x00,
  0x00, 0x00, 0x03, 0xff, 0xff, 0xff, 0xff, 0xff, 0xff, 0xff, 0xff, 0xff, 0xff, 0xff, 0xff, 0xff,
  0xe0, 0x00, 0x00, 0x00, 0x00, 0x00, 0x00, 0x00, 0x00, 0x00, 0x00, 0x00, 0x00, 0x00, 0x00, 0x00,
  0x03, 0xff, 0xff, 0xff, 0xff, 0xff, 0xff, 0xff, 0xff, 0xff, 0xff, 0xff, 0xff, 0xff, 0xe0, 0x00,
  0x00, 0x00, 0x00, 0x00, 0x00, 0x03, 0xf0, 0x00, 0x07, 0xc0, 0x03, 0xff, 0xc0, 0x00, 0x03, 0xff,
  0xff, 0xff, 0xff, 0xff, 0xff, 0xff, 0xff, 0xff, 0xff, 0xff, 0xff, 0xff, 0xe0, 0x00, 0x00, 0x00,
  0x00, 0x00, 0x00, 0x03, 0xf0, 0x00, 0x0f, 0xc0, 0x0f, 0xff, 0xf8, 0x00, 0x03, 0xff, 0xff, 0xff,
  0xff, 0xff, 0xff, 0xff, 0xff, 0xff, 0xff, 0xff, 0xff, 0xff, 0xe0, 0x00, 0x00, 0x00, 0x00, 0x00,
  0x00, 0x03, 0xf0, 0x00, 0x0f, 0xc0, 0x3f, 0xff, 0xfc, 0x00, 0x03, 0xff, 0xff, 0xff, 0xff, 0xff,
  0xff, 0xff, 0xff, 0xff, 0xff, 0xff, 0xff, 0xff, 0xe0, 0x00, 0x00, 0x00, 0x00, 0x00, 0x00, 0x03,
  0xf0, 0x00, 0x0f, 0xc0, 0x7f, 0xff, 0xfe, 0x00, 0x03, 0xfc, 0x00, 0x07, 0xff, 0xc1, 0xff, 0x80,
  0x00, 0xf8, 0x00, 0x00, 0x70, 0x3f, 0xe0, 0x00, 0x00, 0x00, 0x00, 0x00, 0x00, 0x03, 0xf0, 0x00,
  0x0f, 0xc0, 0x7f, 0xff, 0xff, 0x00, 0x03, 0xfc, 0x00, 0x03, 0xff, 0xc1, 0xff, 0x80, 0x00, 0x78,
  0x00, 0x00, 0x70, 0x3f, 0xe0, 0x00, 0x00, 0x00, 0x00, 0x00, 0x00, 0x03, 0xf0, 0x00, 0x0f, 0xc0,
  0xff, 0xff, 0xff, 0x00, 0x03, 0xfc, 0x00, 0x03, 0xff, 0x81, 0xff, 0x80, 0x00, 0x78, 0x00, 0x00,
  0x70, 0x3f, 0xe0, 0x00, 0x00, 0x00, 0x00, 0x00, 0x00, 0x03, 0xf0, 0x00, 0x0f, 0xc0, 0xff, 0xff,
  0xff, 0x80, 0x03, 0xfc, 0x00, 0x03, 0xff, 0x80, 0xff, 0x80, 0x00, 0x78, 0x00, 0x00, 0x70, 0x3f,
  0xe0, 0x00, 0x00, 0x00, 0x00, 0x00, 0x00, 0x03, 0xf0, 0x00, 0x0f, 0xc0, 0xfc, 0x00, 0x3f, 0x80,
  0x03, 0xfc, 0x00, 0x03, 0xff, 0x80, 0xff, 0x80, 0x00, 0x78, 0x00, 0x00, 0x70, 0x3f, 0xe0, 0x00,
  0x00, 0x00, 0x00, 0x00, 0x00, 0x03, 0xf0, 0x00, 0x0f, 0xc1, 0xfc, 0x00, 0x1f, 0x80, 0x03, 0xfc,
  0x0f, 0x83, 0xff, 0x80, 0xff, 0x81, 0xe0, 0x7f, 0xf0, 0x3f, 0xf0, 0x3f, 0xe0, 0x00, 0x00, 0x00,
  0x00, 0x00, 0x00, 0x03, 0xf0, 0x00, 0x0f, 0xc1, 0xfc, 0x00, 0x1f, 0x80, 0x03, 0xfc, 0x0f, 0x83,
  0xff, 0x80, 0xff, 0x83, 0xf0, 0x7f, 0xf8, 0x3f, 0xf0, 0x3f, 0xe0, 0x00, 0x00, 0x00, 0x00, 0x00,
  0x00, 0x03, 0xf0, 0x00, 0x0f, 0xc1, 0xfc, 0x00, 0x1f, 0x80, 0x03, 0xfc, 0x0f, 0x83, 0xff, 0x80,
  0xff, 0x83, 0xf0, 0x7f, 0xf0, 0x3f, 0xf0, 0x3f, 0xe0, 0x00, 0x00, 0x00, 0x00, 0x00, 0x00, 0x03,
  0xf0, 0x00, 0x0f, 0xc1, 0xfc, 0x00, 0x1f, 0x80, 0x03, 0xfc, 0x0f, 0x83, 0xff, 0x00, 0xff, 0x83,
  0xf0, 0x7f, 0xf8, 0x3f, 0xf0, 0x3f, 0xe0, 0x00, 0x00, 0x00, 0x00, 0x00, 0x00, 0x03, 0xf0, 0x00,
  0x0f, 0xc1, 0xfc, 0x00, 0x1f, 0x80, 0x03, 0xfc, 0x0f, 0x83, 0xff, 0x00, 0x7f, 0x83, 0xf0, 0x7f,
  0xf8, 0x3f, 0xf0, 0x3f, 0xe0, 0x00, 0x00, 0x00, 0x00, 0x00, 0x00, 0x03, 0xf0, 0x00, 0x0f, 0xc1,
  0xfc, 0x00, 0x1f, 0x80, 0x03, 0xfc, 0x0f, 0x83, 0xff, 0x00, 0x7f, 0x83, 0xf0, 0x7f, 0xf8, 0x3f,
  0xf0, 0x3f, 0xe0, 0x00, 0x00, 0x00, 0x00, 0x00, 0x00, 0x03, 0xf0, 0x00, 0x0f, 0xc1, 0xfc, 0x00,
  0x1f, 0x80, 0x03, 0xfc, 0x0f, 0x83, 0xff, 0x00, 0x7f, 0x83, 0xf0, 0x7f, 0xf8, 0x3f, 0xf0, 0x3f,
  0xe0, 0x00, 0x00, 0x00, 0x00, 0x00, 0x00, 0x03, 0xf0, 0x00, 0x0f, 0xc1, 0xfc, 0x00, 0x1f, 0x80,
  0x03, 0xfc, 0x0f, 0x83, 0xff, 0x00, 0x7f, 0x83, 0xf0, 0x7f, 0xf8, 0x3f, 0xf0, 0x3f, 0xe0, 0x00,
  0x00, 0x00, 0x00, 0x00, 0x00, 0x03, 0xf0, 0x00, 0x0f, 0xc1, 0xfc, 0x00, 0x1f, 0x80, 0x03, 0xfc,
  0x0f, 0x83, 0xfe, 0x00, 0x7f, 0x83, 0xf0, 0x7f, 0xf8, 0x3f, 0xf0, 0x3f, 0xe0, 0x00, 0x00, 0x00,
  0x00, 0x00, 0x00, 0x03, 0xf0, 0x00, 0x0f, 0xc1, 0xfc, 0x00, 0x1f, 0x80, 0x03, 0xfc, 0x0f, 0x83,
  0xfe, 0x08, 0x7f, 0x83, 0xf0, 0x7f, 0xf8, 0x3f, 0xf0, 0x3f, 0xe0, 0x00, 0x00, 0x00, 0x00, 0x00,
  0x00, 0x03, 0xf0, 0x00, 0x0f, 0xc1, 0xfc, 0x00, 0x1f, 0x80, 0x03, 0xfc, 0x0f, 0x83, 0xfe, 0x08,
  0x7f, 0x83, 0xf0, 0x7f, 0xf8, 0x3f, 0xf0, 0x3f, 0xe0, 0x00, 0x00, 0x00, 0x00, 0x00, 0x00, 0x03,
  0xf0, 0x00, 0x0f, 0xc1, 0xfc, 0x00, 0x1f, 0x80, 0x03, 0xfc, 0x0f, 0x83, 0xfe, 0x08, 0x3f, 0x83,
  0xf0, 0x7f, 0xf8, 0x3f, 0xf0, 0x3f, 0xe0, 0x00, 0x00, 0x00, 0x00, 0x00, 0x00, 0x03, 0xf0, 0x00,
  0x0f, 0xc1, 0xfc, 0x00, 0x1f, 0x80, 0x03, 0xfc, 0x0f, 0x83, 0xfe, 0x18, 0x3f, 0x83, 0xe0, 0x7f,
  0xf8, 0x3f, 0xf0, 0x3f, 0xe0, 0x00, 0x00, 0x00, 0x00, 0x00, 0x00, 0x03, 0xf0, 0x00, 0x0f, 0xc1,
  0xfc, 0x00, 0x1f, 0x80, 0x03, 0xfc, 0x0e, 0x03, 0xfe, 0x18, 0x3f, 0x83, 0xc0, 0x7f, 0xf8, 0x3f,
  0xf0, 0x3f, 0xe0, 0x00, 0x00, 0x00, 0x00, 0x00, 0x00, 0x03, 0xf0, 0x00, 0x0f, 0xc1, 0xfc, 0x00,
  0x1f, 0x80, 0x03, 0xfc, 0x0c, 0x03, 0xfc, 0x18, 0x3f, 0x83, 0x80, 0xff, 0xf8, 0x3f, 0xf0, 0x3f,
  0xe0, 0x00, 0x00, 0x00, 0x00, 0x00, 0x00, 0x03, 0xf0, 0x00, 0x0f, 0xc1, 0xfc, 0x00, 0x1f, 0x80,
  0x03, 0xfc, 0x08, 0x03, 0xfc, 0x18, 0x3f, 0x83, 0x01, 0xff, 0xf8, 0x3f, 0xf0, 0x3f, 0xe0, 0x00,
  0x00, 0x00, 0x00, 0x00, 0x00, 0x03, 0xf0, 0x00, 0x0f, 0xc1, 0xfc, 0x00, 0x1f, 0x80, 0x03, 0xfc,
  0x00, 0x0f, 0xfc, 0x1c, 0x3f, 0x80, 0x03, 0xff, 0xf8, 0x3f, 0xf0, 0x3f, 0xe0, 0x00, 0x00, 0x00,
  0x00, 0x00, 0x00, 0x03, 0xf0, 0x00, 0x0f, 0xc1, 0xfc, 0x00, 0x1f, 0x80, 0x03, 0xfc, 0x00, 0x1f,
  0xfc, 0x1c, 0x1f, 0x80, 0x07, 0xff, 0xf8, 0x3f, 0xf0, 0x3f, 0xe0, 0x00, 0x00, 0x00, 0x00, 0x00,
  0x00, 0x03, 0xf0, 0x00, 0x0f, 0xc1, 0xfc, 0x00, 0x1f, 0x80, 0x03, 0xfc, 0x00, 0x3f, 0xfc, 0x1c,
  0x1f, 0x80, 0x07, 0xff, 0xf8, 0x3f, 0xf0, 0x3f, 0xe0, 0x00, 0x00, 0x00, 0x00, 0x00, 0x00, 0x03,
  0xf0, 0x00, 0x0f, 0xc1, 0xfc, 0x00, 0x1f, 0x80, 0x03, 0xfc, 0x00, 0xff, 0xfc, 0x3c, 0x1f, 0x80,
  0x07, 0xff, 0xf8, 0x3f, 0xf0, 0x3f, 0xe0, 0x00, 0x00, 0x00, 0x00, 0x00, 0x00, 0x03, 0xf0, 0x00,
  0x0f, 0xc1, 0xfc, 0x00, 0x1f, 0x80, 0x03, 0xfc, 0x01, 0xff, 0xf8, 0x3c, 0x1f, 0x80, 0x07, 0xff,
  0xf8, 0x3f, 0xf0, 0x3f, 0xe0, 0x00, 0x00, 0x00, 0x00, 0x00, 0x00, 0x03, 0xf0, 0x00, 0x0f, 0xc1,
  0xfc, 0x00, 0x1f, 0x80, 0x03, 0xfc, 0x03, 0xff, 0xf8, 0x3c, 0x1f, 0x81, 0x03, 0xff, 0xf8, 0x3f,
  0xf0, 0x3f, 0xe0, 0x00, 0x00, 0x00, 0x00, 0x00, 0x00, 0x03, 0xf0, 0x00, 0x0f, 0xc1, 0xfc, 0x00,
  0x1f, 0x80, 0x03, 0xfc, 0x0f, 0xff, 0xf8, 0x3c, 0x1f, 0x83, 0x83, 0xff, 0xf8, 0x3f, 0xf0, 0x3f,
  0xe0, 0x00, 0x00, 0x00, 0x00, 0x00, 0x00, 0x03, 0xf0, 0x00, 0x0f, 0xc1, 0xfc, 0x00, 0x1f, 0x80,
  0x03, 0xfc, 0x0f, 0xff, 0xf8, 0x3c, 0x0f, 0x83, 0x83, 0xff, 0xf8, 0x3f, 0xf0, 0x3f, 0xe0, 0x00,
  0x00, 0x00, 0x00, 0x00, 0x00, 0x03, 0xf0, 0x00, 0x0f, 0xc1, 0xfc, 0x00, 0x1f, 0x80, 0x03, 0xfc,
  0x0f, 0xff, 0xf8, 0x00, 0x0f, 0x83, 0x81, 0xff, 0xf8, 0x3f, 0xf0, 0x3f, 0xe0, 0x00, 0x00, 0x00,
  0x00, 0x00, 0x00, 0x03, 0xf0, 0x00, 0x0f, 0xc1, 0xfc, 0x00, 0x1f, 0x80, 0x03, 0xfc, 0x0f, 0xff,
  0xf8, 0x00, 0x0f, 0x83, 0xc1, 0xff, 0xf8, 0x3f, 0xf0, 0x3f, 0xe0, 0x00, 0x00, 0x00, 0x00, 0x00,
  0x00, 0x03, 0xf0, 0x00, 0x0f, 0xc1, 0xfc, 0x00, 0x1f, 0x80, 0x03, 0xfc, 0x0f, 0xff, 0xf0, 0x00,
  0x0f, 0x83, 0xc1, 0xff, 0xf8, 0x3f, 0xf0, 0x3f, 0xe0, 0x00, 0x00, 0x00, 0x00, 0x00, 0x00, 0x03,
  0xf0, 0x00, 0x0f, 0xc0, 0xfc, 0x00, 0x3f, 0x80, 0x03, 0xfc, 0x0f, 0xff, 0xf0, 0x00, 0x0f, 0x83,
  0xc0, 0xff, 0xf8, 0x3f, 0xf0, 0x3f, 0xe0, 0x00, 0x00, 0x00, 0x00, 0x00, 0x00, 0x03, 0xf0, 0x00,
  0x0f, 0xc0, 0xfe, 0x00, 0x7f, 0x80, 0x03, 0xfc, 0x0f, 0xff, 0xf0, 0x00, 0x0f, 0x83, 0xc0, 0xff,
  0xf8, 0x3f, 0xf0, 0x3f, 0xe0, 0x00, 0x00, 0x00, 0x00, 0x00, 0x00, 0x03, 0xff, 0xfe, 0x0f, 0xc0,
  0xff, 0xff, 0xff, 0x00, 0x03, 0xfc, 0x0f, 0xff, 0xf0, 0x00, 0x07, 0x83, 0xe0, 0xff, 0xf8, 0x3f,
  0xf0, 0x3f, 0xe0, 0x00, 0x00, 0x00, 0x00, 0x00, 0x00, 0x03, 0xff, 0xfe, 0x0f, 0xc0, 0x7f, 0xff,
  0xff, 0x00, 0x03, 0xfc, 0x0f, 0xff, 0xf0, 0x7f, 0x07, 0x83, 0xe0, 0xff, 0xf8, 0x3f, 0xf0, 0x3f,
  0xe0, 0x00, 0x00, 0x00, 0x00, 0x00, 0x00, 0x03, 0xff, 0xfe, 0x0f, 0xc0, 0x7f, 0xff, 0xfe, 0x00,
  0x03, 0xfc, 0x0f, 0xff, 0xe0, 0x7f, 0x07, 0x83, 0xe0, 0x7f, 0xf8, 0x3f, 0xf0, 0x3f, 0xe0, 0x00,
  0x00, 0x00, 0x00, 0x00, 0x00, 0x03, 0xff, 0xfe, 0x0f, 0xc0, 0x3f, 0xff, 0xfc, 0x00, 0x03, 0xfc,
  0x0f, 0xff, 0xe0, 0xff, 0x07, 0x83, 0xe0, 0x7f, 0xf8, 0x3f, 0xf0, 0x3f, 0xe0, 0x00, 0x00, 0x00,
  0x00, 0x00, 0x00, 0x03, 0xff, 0xfe, 0x0f, 0xc0, 0x1f, 0xff, 0xf8, 0x00, 0x03, 0xfc, 0x0f, 0xff,
  0xe0, 0xff, 0x07, 0x83, 0xf0, 0x7f, 0xf8, 0x3f, 0xf0, 0x3f, 0xe0, 0x00, 0x00, 0x00, 0x00, 0x00,
  0x00, 0x03, 0xff, 0xfe, 0x0f, 0xc0, 0x03, 0xff, 0xe0, 0x00, 0x03, 0xfc, 0x0f, 0xff, 0xe0, 0xff,
  0x07, 0x83, 0xf0, 0x3f, 0xf8, 0x3f, 0xf0, 0x3f, 0xe0, 0x00, 0x00, 0x00, 0x00, 0x00, 0x00, 0x00,
  0x00, 0x00, 0x00, 0x00, 0x00, 0x3f, 0xe0, 0x00, 0x03, 0xfc, 0x0f, 0xff, 0xe0, 0xff, 0x03, 0x83,
  0xf0, 0x3f, 0xf8, 0x3f, 0xf0, 0x3f, 0xe0, 0x00, 0x00, 0x00, 0x00, 0x00, 0x00, 0x00, 0x00, 0x00,
  0x00, 0x00, 0x00, 0x1f, 0xf0, 0x00, 0x03, 0xff, 0xff, 0xff, 0xff, 0xff, 0xff, 0xff, 0xff, 0xff,
  0xff, 0xff, 0xff, 0xff, 0xe0, 0x00, 0x00, 0x00, 0x00, 0x00, 0x00, 0x00, 0x00, 0x00, 0x00, 0x00,
  0x00, 0x0f, 0xf0, 0x00, 0x03, 0xff, 0xff, 0xff, 0xff, 0xff, 0xff, 0xff, 0xff, 0xff, 0xff, 0xff,
  0xff, 0xff, 0xe0, 0x00, 0x00, 0x00, 0x00, 0x00, 0x00, 0x00, 0x00, 0x00, 0x00, 0x00, 0x00, 0x07,
  0xe0, 0x00, 0x03, 0xff, 0xff, 0xff, 0xff, 0xff, 0xff, 0xff, 0xff, 0xff, 0xff, 0xff, 0xff, 0xff,
  0xe0, 0x00, 0x00, 0x00, 0x00, 0x00, 0x00, 0x00, 0x00, 0x00, 0x00, 0x00, 0x00, 0x03, 0xc0, 0x00,
  0x03, 0xff, 0xff, 0xff, 0xff, 0xff, 0xff, 0xff, 0xff, 0xff, 0xff, 0xff, 0xff, 0xff, 0xe0, 0x00,
  0x00, 0x00, 0x00, 0x00, 0x00, 0x00, 0x00, 0x00, 0x00, 0x00, 0x00, 0x01, 0x80, 0x00, 0x03, 0xff,
  0xff, 0xff, 0xff, 0xff, 0xff, 0xff, 0xff, 0xff, 0xff, 0xff, 0xff, 0xff, 0xe0, 0x00, 0x00, 0x00,
  0x00, 0x00, 0x00, 0x00, 0x00, 0x00, 0x00, 0x00, 0x00, 0x00, 0x00, 0x00, 0x03, 0xff, 0xff, 0xff,
  0xff, 0xff, 0xff, 0xff, 0xff, 0xff, 0xff, 0xff, 0xff, 0xff, 0xe0, 0x00, 0x00, 0x00, 0x00, 0x00,
  0x00, 0x00, 0x00, 0x00, 0x00, 0x00, 0x00, 0x00, 0x00, 0x00, 0x03, 0xff, 0xff, 0xff, 0xff, 0xff,
  0xff, 0xff, 0xff, 0xff, 0xff, 0xff, 0xff, 0xff, 0xe0, 0x00, 0x00, 0x00, 0x00, 0x00, 0x00, 0x00,
  0x00, 0x00, 0x00, 0x00, 0x00, 0x00, 0x00, 0x00, 0x00, 0x00, 0x00, 0x00, 0x00, 0x00, 0x00, 0x00,
  0x00, 0x00, 0x00, 0x00, 0x00, 0x00, 0x00, 0x00, 0x00, 0x00, 0x00, 0x00, 0x00, 0x00, 0x00, 0x00,
  0x00, 0x00, 0x00, 0x00, 0x00, 0x00, 0x00, 0x00, 0x00, 0x00, 0x00, 0x00, 0x00, 0x00, 0x00, 0x00,
  0x00, 0x00, 0x00, 0x00, 0x00, 0x00, 0x00, 0x00, 0x00, 0x00, 0x00, 0x00, 0x00, 0x00, 0x00, 0x00,
  0x00, 0x00, 0x00, 0x00, 0x00, 0x00, 0x00, 0x00, 0x00, 0x00, 0x00, 0x00, 0x00, 0x00, 0x00, 0x00,
  0x00, 0x00, 0x00, 0x00, 0x00, 0x00, 0x00, 0x00, 0x00, 0x00, 0x00, 0x00, 0x00, 0x00, 0x00, 0x00,
  0x00, 0x00, 0x00, 0x00, 0x00, 0x00, 0x00, 0x00, 0x00, 0x00, 0x00, 0x00, 0x00, 0x00, 0x00, 0x00,
  0x00, 0x00, 0x00, 0x00, 0x00, 0x00, 0x00, 0x00, 0x00, 0x00, 0x00, 0x00, 0x00, 0x00, 0x00, 0x00,
  0x00, 0x00, 0x00, 0x00, 0x00, 0x00, 0x00, 0x00, 0x00, 0x00, 0x00, 0x00, 0x00, 0x00, 0x00, 0x00,
  0x00, 0x00, 0x00, 0x00, 0x00, 0x00, 0x00, 0x00, 0x00, 0x00, 0x00, 0x00, 0x00, 0x00, 0x00, 0x00,
  0x00, 0x00, 0x00, 0x00, 0x00, 0x00, 0x00, 0x00, 0x00, 0x00, 0x00, 0x00, 0x00, 0x00, 0x00, 0x00,
  0x00, 0x00, 0x00, 0x00, 0x00, 0x00, 0x00, 0x00, 0x00, 0x00, 0x00, 0x00, 0x00, 0x00, 0x00, 0x00,
  0x00, 0x00, 0x00, 0x00, 0x00, 0x00, 0x00, 0x00, 0x00, 0x00, 0x00, 0x00, 0x00, 0x00, 0x00, 0x00,
  0x00, 0x00, 0x00, 0x00, 0x00, 0x00, 0x00, 0x00, 0x00, 0x00, 0x00, 0x00, 0x00, 0x00, 0x00, 0x00,
  0x00, 0x00, 0x00, 0x00, 0x00, 0x00, 0x00, 0x00, 0x00, 0x00, 0x00, 0x00, 0x00, 0x00, 0x00, 0x00,
  0x00, 0x00, 0x00, 0x00, 0x00, 0x00, 0x00, 0x00, 0x00, 0x00, 0x00, 0x00, 0x00, 0x00, 0x00, 0x00,
  0x00, 0x00, 0x00, 0x00, 0x00, 0x00, 0x00, 0x00, 0x00, 0x00, 0x00, 0x00, 0x00, 0x00, 0x00, 0x00,
  0x00, 0x00, 0x00, 0x00, 0x00, 0x00, 0x00, 0x00, 0x00, 0x00, 0x00, 0x00, 0x00, 0x00, 0x00, 0x00,
  0x00, 0x00, 0x00, 0x00, 0x00, 0x00, 0x00, 0x00, 0x00, 0x00, 0x00, 0x00, 0x00, 0x00, 0x00, 0x00,
  0x00, 0x00, 0x00, 0x00, 0x00, 0x00, 0x00, 0x00, 0x00, 0x00, 0x00, 0x00, 0x00, 0x00, 0x00, 0x00,
  0x00, 0x00, 0x00, 0x00, 0x00, 0x00, 0x00, 0x00, 0x00, 0x00, 0x00, 0x00, 0x00, 0x00, 0x00, 0x00,
  0x00, 0x00, 0x00, 0x00, 0x00, 0x00, 0x00, 0x00, 0x00, 0x00, 0x00, 0x00, 0x00, 0x00, 0x00, 0x00,
  0x00, 0x00, 0x00, 0x00, 0x00, 0x00, 0x00, 0x00, 0x00, 0x00, 0x00, 0x00, 0x00, 0x00, 0x00, 0x00,
  0x00, 0x00, 0x00, 0x00, 0x00, 0x00, 0x00, 0x00, 0x00, 0x00, 0x00, 0x00, 0x00, 0x00, 0x00, 0x00,
  0x00, 0x00, 0x00, 0x00, 0x00, 0x00, 0x00, 0x00, 0x00, 0x00, 0x00, 0x00, 0x00, 0x00, 0x00, 0x00,
  0x00, 0x00, 0x00, 0x00, 0x00, 0x00, 0x00, 0x00, 0x00, 0x00, 0x00, 0x00, 0x00, 0x00, 0x00, 0x00,
  0x00, 0x00, 0x00, 0x00, 0x00, 0x00, 0x00, 0x00, 0x00, 0x00, 0x00, 0x00, 0x00, 0x00, 0x00, 0x00,
  0x00, 0x00, 0x00, 0x00, 0x00, 0x00, 0x00, 0x00, 0x00, 0x00, 0x00, 0x00, 0x00, 0x00, 0x00, 0x00,
  0x00, 0x00, 0x00, 0x00, 0x00, 0x00, 0x00, 0x00, 0x00, 0x00, 0x00, 0x00, 0x00, 0x00, 0x00, 0x00,
  0x00, 0x00, 0x00, 0x00, 0x00, 0x00, 0x00, 0x00, 0x00, 0x00, 0x00, 0x00, 0x00, 0x00, 0x00, 0x00,
  0x00, 0x00, 0x00, 0x00, 0x00, 0x00, 0x00, 0x00, 0x00, 0x00, 0x00, 0x00, 0x00, 0x00, 0x00, 0x00,
  0x00, 0x00, 0x00, 0x00, 0x00, 0x00, 0x00, 0x00, 0x00, 0x00, 0x00, 0x00, 0x00, 0x00, 0x00, 0x00,
  0x00, 0x00, 0x00, 0x00, 0x00, 0x00, 0x00, 0x00, 0x00, 0x00, 0x00, 0x00, 0x00, 0x00, 0x00, 0x00,
  0x00, 0x00, 0x00, 0x00, 0x00, 0x00, 0x00, 0x00, 0x00, 0x00, 0x00, 0x00, 0x00, 0x00, 0x00, 0x00,
  0x00, 0x00, 0x00, 0x00, 0x00, 0x00, 0x00, 0x00, 0x00, 0x00, 0x00, 0x00, 0x00, 0x00, 0x00, 0x00,
  0x00, 0x00, 0x00, 0x00, 0x00, 0x00, 0x00, 0x00, 0x00, 0x00, 0x00, 0x00, 0x00, 0x00, 0x00, 0x00,
  0x00, 0x00, 0x00, 0x00, 0x00, 0x00, 0x00, 0x00, 0x00, 0x00, 0x00, 0x00, 0x00, 0x00, 0x00, 0x00,
  0x00, 0x00, 0x00, 0x00, 0x00, 0x00, 0x00, 0x00, 0x00, 0x00, 0x00, 0x00, 0x00, 0x00, 0x00, 0x00,
  0x00, 0x00, 0x00, 0x00, 0x00, 0x00, 0x00, 0x00, 0x00, 0x00, 0x00, 0x00, 0x00, 0x00, 0x00, 0x00,
  0x00, 0x00, 0x00, 0x00, 0x00, 0x00, 0x00, 0x00, 0x00, 0x00, 0x00, 0x00, 0x00, 0x00, 0x00, 0x00,
  0x00, 0x00, 0x00, 0x00, 0x00, 0x00, 0x00, 0x00, 0x00, 0x00, 0x00, 0x00, 0x00, 0x00, 0x00, 0x00,
  0x00, 0x00, 0x00, 0x00, 0x00, 0x00, 0x00, 0x00, 0x00, 0x00, 0x00, 0x00, 0x00, 0x00, 0x00, 0x00,
  0x00, 0x00, 0x00, 0x00, 0x00, 0x00, 0x00, 0x00, 0x00, 0x00, 0x00, 0x00, 0x00, 0x00, 0x00, 0x00,
  0x00, 0x00, 0x00, 0x00, 0x00, 0x00, 0x00, 0x00, 0x00, 0x00, 0x00, 0x00, 0x00, 0x00, 0x00, 0x00,
  0x00, 0x00, 0x00, 0x00, 0x00, 0x00, 0x00, 0x00, 0x00, 0x00, 0x00, 0x00, 0x00, 0x00, 0x00, 0x00,
  0x00, 0x00, 0x00, 0x00, 0x00, 0x00, 0x00, 0x00, 0x00, 0x00, 0x00, 0x00, 0x00, 0x00, 0x00, 0x00,
  0x00, 0x00, 0x00, 0x00, 0x00, 0x00, 0x00, 0x00, 0x00, 0x00, 0x00, 0x00, 0x00, 0x00, 0x00, 0x00,
  0x00, 0x00, 0x00, 0x00, 0x00, 0x00, 0x00, 0x00, 0x00, 0x00, 0x00, 0x00, 0x00, 0x00, 0x00, 0x00,
  0x00, 0x00, 0x00, 0x00, 0x00, 0x00, 0x00, 0x00, 0x00, 0x00, 0x00, 0x00, 0x00, 0x00, 0x00, 0x00,
  0x00, 0x00, 0x00, 0x00, 0x00, 0x00, 0x00, 0x00, 0x00, 0x00, 0x00, 0x00, 0x00, 0x00, 0x00, 0x00,
  0x00, 0x00, 0x00, 0x00, 0x00, 0x00, 0x00, 0x00, 0x00, 0x00, 0x00, 0x00, 0x00, 0x00, 0x00, 0x00,
  0x00, 0x00, 0x00, 0x00, 0x00, 0x00, 0x00, 0x00, 0x00, 0x00, 0x00, 0x00, 0x00, 0x00, 0x00, 0x00,
  0x00, 0x00, 0x00, 0x00, 0x00, 0x00, 0x00, 0x00, 0x00, 0x00, 0x00, 0x00, 0x00, 0x00, 0x00, 0x00,
  0x00, 0x00, 0x00, 0x00, 0x00, 0x00, 0x00, 0x00, 0x00, 0x00, 0x00, 0x00, 0x00, 0x00, 0x00, 0x00,
  0x00, 0x00, 0x00, 0x00, 0x00, 0x00, 0x00, 0x00, 0x00, 0x00, 0x00, 0x00, 0x00, 0x00, 0x00, 0x00,
  0x00, 0x00, 0x00, 0x00, 0x00, 0x00, 0x00, 0x00, 0x00, 0x00, 0x00, 0x00, 0x00, 0x00, 0x00, 0x00,
  0x00, 0x00, 0x00, 0x00, 0x00, 0x00, 0x00, 0x00, 0x00, 0x00, 0x00, 0x00, 0x00, 0x00, 0x00, 0x00,
  0x00, 0x00, 0x00, 0x00, 0x00, 0x00, 0x00, 0x00, 0x00, 0x00, 0x00, 0x00, 0x00, 0x00, 0x00, 0x00,
  0x00, 0x00, 0x00, 0x00, 0x00, 0x00, 0x00, 0x00, 0x00, 0x00, 0x00, 0x00, 0x00, 0x00, 0x00, 0x00,
  0x00, 0x00, 0x00, 0x00, 0x00, 0x00, 0x00, 0x00, 0x00, 0x00, 0x00, 0x00, 0x00, 0x00, 0x00, 0x00,
  0x00, 0x00, 0x00, 0x00, 0x00, 0x00, 0x00, 0x00, 0x00, 0x00, 0x00, 0x00, 0x00, 0x00, 0x00, 0x00,
  0x00, 0x00, 0x00, 0x00, 0x00, 0x00, 0x00, 0x00, 0x00, 0x00, 0x00, 0x00, 0x00, 0x00, 0x00, 0x00,
  0x00, 0x00, 0x00, 0x00, 0x00, 0x00, 0x00, 0x00, 0x00, 0x00, 0x00, 0x00, 0x00, 0x00, 0x00, 0x00,
  0x00, 0x00, 0x00, 0x00, 0x00, 0x00, 0x00, 0x00, 0x00, 0x00, 0x00, 0x00, 0x00, 0x00, 0x00, 0x00,
  0x00, 0x00, 0x00, 0x00, 0x00, 0x00, 0x00, 0x00, 0x00, 0x00, 0x00, 0x00, 0x00, 0x00, 0x00, 0x00,
  0x00, 0x00, 0x00, 0x00, 0x00, 0x00, 0x00, 0x00, 0x00, 0x00, 0x00, 0x00, 0x00, 0x00, 0x00, 0x00,
  0x00, 0x00, 0x00, 0x00, 0x00, 0x00, 0x00, 0x00, 0x00, 0x00, 0x00, 0x00, 0x00, 0x00, 0x00, 0x00,
  0x00, 0x00, 0x00, 0x00, 0x00, 0x00, 0x00, 0x00, 0x00, 0x00, 0x00, 0x00, 0x00, 0x00, 0x00, 0x00,
  0x00, 0x00, 0x00, 0x00, 0x00, 0x00, 0x00, 0x00, 0x00, 0x00, 0x00, 0x00, 0x00, 0x00, 0x00, 0x00,
  0x00, 0x00, 0x00, 0x00, 0x00, 0x00, 0x00, 0x00, 0x00, 0x00, 0x00, 0x00, 0x00, 0x00, 0x00, 0x00,
  0x00, 0x00, 0x00, 0x00, 0x00, 0x00, 0x00, 0x00, 0x00, 0x00, 0x00, 0x00, 0x00, 0x00, 0x00, 0x00,
  0x00, 0x00, 0x00, 0x00, 0x00, 0x00, 0x00, 0x00, 0x00, 0x00, 0x00, 0x00, 0x00, 0x00, 0x00, 0x00,
  0x00, 0x00, 0x00, 0x00, 0x00, 0x00, 0x00, 0x00, 0x00, 0x00, 0x00, 0x00, 0x00, 0x00, 0x00, 0x00,
  0x00, 0x00, 0x00, 0x00, 0x00, 0x00, 0x00, 0x00, 0x00, 0x00, 0x00, 0x00, 0x00, 0x00, 0x00, 0x00,
  0x00, 0x00, 0x00, 0x00, 0x00, 0x00, 0x00, 0x00, 0x00, 0x00, 0x00, 0x00, 0x00, 0x00, 0x00, 0x07,
  0xf3, 0xff, 0x3e, 0x03, 0xff, 0xff, 0xff, 0xff, 0xff, 0xff, 0xff, 0xff, 0xff, 0xff, 0xff, 0xff,
  0xff, 0xff, 0xff, 0xff, 0xff, 0xf8, 0x00, 0x00, 0x00, 0x00, 0x00, 0x00, 0x00, 0x07, 0xfb, 0xff,
  0x3e, 0x1b, 0xff, 0xff, 0xff, 0xff, 0xff, 0xff, 0xff, 0xff, 0xff, 0xff, 0xff, 0xff, 0xff, 0xff,
  0xff, 0xff, 0xff, 0xf8, 0x00, 0x00, 0x00, 0x00, 0x00, 0x00, 0x00, 0x07, 0xfb, 0xff, 0x3e, 0x1d,
  0xff, 0xff, 0xff, 0xff, 0xff, 0xff, 0xff, 0xff, 0xff, 0xff, 0xff, 0xff, 0xff, 0xff, 0xff, 0xff,
  0xff, 0xf8, 0x00, 0x00, 0x00, 0x00, 0x00, 0x00, 0x00, 0x07, 0xf8, 0x00, 0x00, 0x1e, 0x00, 0x00,
  0x00, 0x00, 0x00, 0x00, 0x00, 0x00, 0x00, 0x00, 0x00, 0x00, 0x00, 0x00, 0x00, 0x00, 0x00, 0x00,
  0x00, 0x00, 0x00, 0x00, 0x00, 0x00, 0x00, 0x07, 0xf8, 0x00, 0x00, 0x1f, 0x00, 0x00, 0x00, 0x00,
  0x00, 0x00, 0x00, 0x00, 0x00, 0x00, 0x00, 0x00, 0x00, 0x00, 0x00, 0x00, 0x00, 0x00, 0x00, 0x00,
  0x00, 0x00, 0x00, 0x00, 0x00, 0x07, 0xf8, 0x7f, 0x3f, 0x9f, 0x8f, 0xf3, 0xff, 0xfe, 0x07, 0xff,
  0xfd, 0xff, 0xff, 0x00, 0x00, 0xfe, 0x00, 0xff, 0xff, 0xbf, 0xff, 0xe0, 0x00, 0x00, 0x00, 0x00,
  0x00, 0x00, 0x00, 0x07, 0xf8, 0x7f, 0x3f, 0x9f, 0xcf, 0xf3, 0xff, 0xff, 0x0f, 0xff, 0xfd, 0xff,
  0xff, 0x80, 0x00, 0xfe, 0x03, 0xff, 0xff, 0xbf, 0xff, 0xf0, 0x00, 0x00, 0x00, 0x00, 0x00, 0x00,
  0x00, 0x07, 0xfa, 0x7f, 0x3f, 0x9f, 0xef, 0xf3, 0xff, 0xff, 0x1f, 0xff, 0xf9, 0xff, 0xff, 0xc0,
  0x00, 0xfe, 0x03, 0xff, 0xff, 0xbf, 0xff, 0xf8, 0x00, 0x00, 0x00, 0x00, 0x00, 0x00, 0x00, 0x07,
  0xff, 0x7f, 0x3f, 0x9f, 0xff, 0xf3, 0xf8, 0x7f, 0x9f, 0xe0, 0x00, 0x00, 0x3f, 0xc0, 0x00, 0xfe,
  0x03, 0xf8, 0x7f, 0xbf, 0xc7, 0xf8, 0x00, 0x00, 0x00, 0x00, 0x00, 0x00, 0x00, 0x07, 0xff, 0xff,
  0x3f, 0x9f, 0xff, 0xf3, 0xf8, 0x7f, 0x9f, 0xc0, 0x00, 0x00, 0x7f, 0xc0, 0x00, 0xfe, 0x07, 0xf8,
  0x7f, 0xbf, 0xc7, 0xf8, 0x00, 0x00, 0x00, 0x00, 0x00, 0x00, 0x00, 0x07, 0xff, 0xff, 0x3f, 0x9f,
  0xff, 0xf3, 0xf8, 0x7f, 0x9f, 0xfe, 0x01, 0xff, 0xff, 0x80, 0x00, 0xfe, 0x07, 0xff, 0xff, 0xbf,
  0xff, 0xf8, 0x00, 0x00, 0x00, 0x00, 0x00, 0x00, 0x00, 0x07, 0xff, 0xff, 0x3f, 0x9f, 0xff, 0xf3,
  0xf8, 0x7f, 0x9f, 0xfe, 0x01, 0xff, 0xff, 0x00, 0x00, 0xfe, 0x07, 0xff, 0xff, 0xbf, 0xff, 0xf8,
  0x00, 0x00, 0x00, 0x00, 0x00, 0x00, 0x00, 0x07, 0xff, 0xff, 0x3f, 0x9f, 0xff, 0xf3, 0xf8, 0x7f,
  0x9f, 0xc0, 0x01, 0xfe, 0x7f, 0x00, 0x00, 0xfe, 0x07, 0xf8, 0x7f, 0xbf, 0xc7, 0xf8, 0x00, 0x00,
  0x00, 0x00, 0x00, 0x00, 0x00, 0x07, 0xfd, 0xff, 0x3f, 0x9f, 0xff, 0xf3, 0xf8, 0x7f, 0x9f, 0xc0,
  0x01, 0xfc, 0x3f, 0x80, 0x00, 0xfe, 0x07, 0xf8, 0x7f, 0xbf, 0xc3, 0xf8, 0x00, 0x00, 0x00, 0x00,
  0x00, 0x00, 0x00, 0x07, 0xf8, 0x7f, 0x3f, 0x9f, 0xef, 0xf3, 0xff, 0xff, 0x9f, 0xff, 0xfd, 0xfc,
  0x3f, 0xc0, 0x00, 0xff, 0xf7, 0xf8, 0x7f, 0xbf, 0xff, 0xf8, 0x00, 0x00, 0x00, 0x00, 0x00, 0x00,
  0x00, 0x07, 0xf0, 0x7f, 0x3f, 0x9f, 0xe7, 0xf3, 0xff, 0xff, 0x9f, 0xff, 0xfd, 0xfc, 0x3f, 0xc0,
  0x00, 0xff, 0xf7, 0xf8, 0x7f, 0xbf, 0xff, 0xf8, 0x00, 0x00, 0x00, 0x00, 0x00, 0x00, 0x00, 0x07,
  0xe0, 0x3f, 0x3f, 0x9f, 0xe3, 0xf3, 0xff, 0xff, 0x9f, 0xff, 0xfd, 0xfc, 0x1f, 0xc0, 0x00, 0xff,
  0xf7, 0xf8, 0x7f, 0xbf, 0xff, 0xf8, 0x00, 0x00, 0x00, 0x00, 0x00, 0x00, 0x00, 0x00, 0x00, 0x1f,
  0x00, 0x00, 0x01, 0xf0, 0x00, 0x00, 0x00, 0x00, 0x00, 0x00, 0x00, 0x00, 0x00, 0x00, 0x00, 0x00,
  0x7f, 0x80, 0x00, 0x00, 0x00, 0x00, 0x00, 0x00, 0x00, 0x00, 0x00, 0x00, 0x00, 0x1f, 0x00, 0x00,
  0x00, 0xf0, 0x00, 0x00, 0x00, 0x00, 0x00, 0x00, 0x00, 0x00, 0x00, 0x00, 0x00, 0x00, 0x7f, 0x80,
  0x00, 0x00, 0x00, 0x00, 0x00, 0x00, 0x00, 0x00, 0x00, 0x03, 0xff, 0xff, 0x3f, 0xff, 0xff, 0x73,
  0xff, 0xff, 0xff, 0xff, 0xff, 0xff, 0xff, 0xff, 0xff, 0xff, 0xff, 0xff, 0xff, 0xbf, 0xff, 0xff,
  0x00, 0x00, 0x00, 0x00, 0x00, 0x00, 0x00, 0x07, 0xff, 0xff, 0x3f, 0xff, 0xff, 0xb3, 0xff, 0xff,
  0xff, 0xff, 0xff, 0xff, 0xff, 0xff, 0xff, 0xff, 0xff, 0xff, 0xff, 0xbf, 0xff, 0xff, 0x00, 0x00,
  0x00, 0x00, 0x00, 0x00, 0x00, 0x03, 0xff, 0xf8, 0x3f, 0xff, 0xff, 0x83, 0xff, 0xff, 0xff, 0xff,
  0xff, 0xff, 0xff, 0xff, 0xff, 0xff, 0xff, 0xff, 0x7f, 0x3f, 0xff, 0xff, 0x00, 0x00, 0x00, 0x00,
  0x00, 0x00, 0x00, 0x00, 0x00, 0x00, 0x00, 0x00, 0x00, 0x00, 0x00, 0x00, 0x00, 0x00, 0x00, 0x00,
  0x00, 0x00, 0x00, 0x00, 0x00, 0x00, 0x00, 0x00, 0x00, 0x00, 0x00, 0x00, 0x00, 0x00, 0x00, 0x00,
  0x00, 0x00, 0x00, 0x00, 0x00, 0x00, 0x00, 0x00, 0x00, 0x00, 0x00, 0x00, 0x00, 0x00, 0x00, 0x00,
  0x00, 0x00, 0x00, 0x00, 0x00, 0x00, 0x00, 0x00, 0x00, 0x00, 0x00, 0x00, 0x00, 0x00, 0x00, 0x00,
  0x00, 0x00, 0x00, 0x00, 0x00, 0x00, 0x00, 0x00, 0x00, 0x00, 0x00, 0x00, 0x00, 0x00, 0x00, 0x00,
  0x00, 0x00, 0x00, 0x00, 0x00, 0x00, 0x00, 0x00, 0x00, 0x00
};

// 'LIQ PARTI Logo', 240x60px
const unsigned char PARTI_LOGOLIQ_PARTI_Logo[] PROGMEM = {
  0x00, 0x00, 0x00, 0x00, 0x00, 0x00, 0x00, 0x00, 0x00, 0x00, 0x00, 0x00, 0x00, 0x00, 0x00, 0x00,
  0x00, 0x00, 0x00, 0x00, 0x00, 0x00, 0x00, 0x00, 0x00, 0x00, 0x00, 0x00, 0x00, 0x00, 0x00, 0x00,
  0x00, 0x00, 0x00, 0x00, 0x00, 0x00, 0x00, 0x00, 0x00, 0x00, 0x00, 0x00, 0x00, 0x00, 0x00, 0x00,
  0x00, 0x00, 0x00, 0x00, 0x00, 0x00, 0x00, 0x00, 0x00, 0x00, 0x00, 0x00, 0x00, 0x00, 0x00, 0x00,
  0x00, 0x00, 0x00, 0x00, 0x00, 0x00, 0x00, 0x00, 0x00, 0x00, 0x00, 0x00, 0x00, 0x00, 0x00, 0x00,
  0x00, 0x00, 0x00, 0x00, 0x00, 0x00, 0x20, 0x00, 0x00, 0x00, 0x00, 0x00, 0x00, 0x00, 0x00, 0x00,
  0x00, 0x00, 0x00, 0x00, 0x00, 0x00, 0x03, 0xff, 0xff, 0xff, 0xff, 0xff, 0xff, 0xff, 0xff, 0xff,
  0xff, 0xff, 0xff, 0xff, 0xe0, 0x00, 0x00, 0x00, 0x00, 0x00, 0x00, 0x00, 0x00, 0x00, 0x00, 0x00,
  0x00, 0x00, 0x00, 0x00, 0x03, 0xff, 0xff, 0xff, 0xff, 0xff, 0xff, 0xff, 0xff, 0xff, 0xff, 0xff,
  0xff, 0xff, 0xe0, 0x00, 0x00, 0x00, 0x00, 0x00, 0x00, 0x00, 0x00, 0x00, 0x00, 0x00, 0x00, 0x00,
  0x00, 0x00, 0x03, 0xff, 0xff, 0xff, 0xff, 0xff, 0xff, 0xff, 0xff, 0xff, 0xff, 0xff, 0xff, 0xff,
  0xe0, 0x00, 0x00, 0x00, 0x00, 0x00, 0x00, 0x00, 0x00, 0x00, 0x00, 0x00, 0x00, 0x00, 0x00, 0x00,
  0x03, 0xff, 0xff, 0xff, 0xff, 0xff, 0xff, 0xff, 0xff, 0xff, 0xff, 0xff, 0xff, 0xff, 0xe0, 0x00,
  0x00, 0x00, 0x00, 0x00, 0x00, 0x03, 0xf0, 0x00, 0x07, 0xc0, 0x03, 0xff, 0xc0, 0x00, 0x03, 0xff,
  0xff, 0xff, 0xff, 0xff, 0xff, 0xff, 0xff, 0xff, 0xff, 0xff, 0xff, 0xff, 0xe0, 0x00, 0x00, 0x00,
  0x00, 0x00, 0x00, 0x03, 0xf0, 0x00, 0x0f, 0xc0, 0x0f, 0xff, 0xf8, 0x00, 0x03, 0xff, 0xff, 0xff,
  0xff, 0xff, 0xff, 0xff, 0xff, 0xff, 0xff, 0xff, 0xff, 0xff, 0xe0, 0x00, 0x00, 0x00, 0x00, 0x00,
  0x00, 0x03, 0xf0, 0x00, 0x0f, 0xc0, 0x3f, 0xff, 0xfc, 0x00, 0x03, 0xff, 0xff, 0xff, 0xff, 0xff,
  0xff, 0xff, 0xff, 0xff, 0xff, 0xff, 0xff, 0xff, 0xe0, 0x00, 0x00, 0x00, 0x00, 0x00, 0x00, 0x03,
  0xf0, 0x00, 0x0f, 0xc0, 0x7f, 0xff, 0xfe, 0x00, 0x03, 0xfc, 0x00, 0x07, 0xff, 0xc1, 0xff, 0x80,
  0x00, 0xf8, 0x00, 0x00, 0x70, 0x3f, 0xe0, 0x00, 0x00, 0x00, 0x00, 0x00, 0x00, 0x03, 0xf0, 0x00,
  0x0f, 0xc0, 0x7f, 0xff, 0xff, 0x00, 0x03, 0xfc, 0x00, 0x03, 0xff, 0xc1, 0xff, 0x80, 0x00, 0x78,
  0x00, 0x00, 0x70, 0x3f, 0xe0, 0x00, 0x00, 0x00, 0x00, 0x00, 0x00, 0x03, 0xf0, 0x00, 0x0f, 0xc0,
  0xff, 0xff, 0xff, 0x00, 0x03, 0xfc, 0x00, 0x03, 0xff, 0x81, 0xff, 0x80, 0x00, 0x78, 0x00, 0x00,
  0x70, 0x3f, 0xe0, 0x00, 0x00, 0x00, 0x00, 0x00, 0x00, 0x03, 0xf0, 0x00, 0x0f, 0xc0, 0xff, 0xff,
  0xff, 0x80, 0x03, 0xfc, 0x00, 0x03, 0xff, 0x80, 0xff, 0x80, 0x00, 0x78, 0x00, 0x00, 0x70, 0x3f,
  0xe0, 0x00, 0x00, 0x00, 0x00, 0x00, 0x00, 0x03, 0xf0, 0x00, 0x0f, 0xc0, 0xfc, 0x00, 0x3f, 0x80,
  0x03, 0xfc, 0x00, 0x03, 0xff, 0x80, 0xff, 0x80, 0x00, 0x78, 0x00, 0x00, 0x70, 0x3f, 0xe0, 0x00,
  0x00, 0x00, 0x00, 0x00, 0x00, 0x03, 0xf0, 0x00, 0x0f, 0xc1, 0xfc, 0x00, 0x1f, 0x80, 0x03, 0xfc,
  0x0f, 0x83, 0xff, 0x80, 0xff, 0x81, 0xe0, 0x7f, 0xf0, 0x3f, 0xf0, 0x3f, 0xe0, 0x00, 0x00, 0x00,
  0x00, 0x00, 0x00, 0x03, 0xf0, 0x00, 0x0f, 0xc1, 0xfc, 0x00, 0x1f, 0x80, 0x03, 0xfc, 0x0f, 0x83,
  0xff, 0x80, 0xff, 0x83, 0xf0, 0x7f, 0xf8, 0x3f, 0xf0, 0x3f, 0xe0, 0x00, 0x00, 0x00, 0x00, 0x00,
  0x00, 0x03, 0xf0, 0x00, 0x0f, 0xc1, 0xfc, 0x00, 0x1f, 0x80, 0x03, 0xfc, 0x0f, 0x83, 0xff, 0x80,
  0xff, 0x83, 0xf0, 0x7f, 0xf0, 0x3f, 0xf0, 0x3f, 0xe0, 0x00, 0x00, 0x00, 0x00, 0x00, 0x00, 0x03,
  0xf0, 0x00, 0x0f, 0xc1, 0xfc, 0x00, 0x1f, 0x80, 0x03, 0xfc, 0x0f, 0x83, 0xff, 0x00, 0xff, 0x83,
  0xf0, 0x7f, 0xf8, 0x3f, 0xf0, 0x3f, 0xe0, 0x00, 0x00, 0x00, 0x00, 0x00, 0x00, 0x03, 0xf0, 0x00,
  0x0f, 0xc1, 0xfc, 0x00, 0x1f, 0x80, 0x03, 0xfc, 0x0f, 0x83, 0xff, 0x00, 0x7f, 0x83, 0xf0, 0x7f,
  0xf8, 0x3f, 0xf0, 0x3f, 0xe0, 0x00, 0x00, 0x00, 0x00, 0x00, 0x00, 0x03, 0xf0, 0x00, 0x0f, 0xc1,
  0xfc, 0x00, 0x1f, 0x80, 0x03, 0xfc, 0x0f, 0x83, 0xff, 0x00, 0x7f, 0x83, 0xf0, 0x7f, 0xf8, 0x3f,
  0xf0, 0x3f, 0xe0, 0x00, 0x00, 0x00, 0x00, 0x00, 0x00, 0x03, 0xf0, 0x00, 0x0f, 0xc1, 0xfc, 0x00,
  0x1f, 0x80, 0x03, 0xfc, 0x0f, 0x83, 0xff, 0x00, 0x7f, 0x83, 0xf0, 0x7f, 0xf8, 0x3f, 0xf0, 0x3f,
  0xe0, 0x00, 0x00, 0x00, 0x00, 0x00, 0x00, 0x03, 0xf0, 0x00, 0x0f, 0xc1, 0xfc, 0x00, 0x1f, 0x80,
  0x03, 0xfc, 0x0f, 0x83, 0xff, 0x00, 0x7f, 0x83, 0xf0, 0x7f, 0xf8, 0x3f, 0xf0, 0x3f, 0xe0, 0x00,
  0x00, 0x00, 0x00, 0x00, 0x00, 0x03, 0xf0, 0x00, 0x0f, 0xc1, 0xfc, 0x00, 0x1f, 0x80, 0x03, 0xfc,
  0x0f, 0x83, 0xfe, 0x00, 0x7f, 0x83, 0xf0, 0x7f, 0xf8, 0x3f, 0xf0, 0x3f, 0xe0, 0x00, 0x00, 0x00,
  0x00, 0x00, 0x00, 0x03, 0xf0, 0x00, 0x0f, 0xc1, 0xfc, 0x00, 0x1f, 0x80, 0x03, 0xfc, 0x0f, 0x83,
  0xfe, 0x08, 0x7f, 0x83, 0xf0, 0x7f, 0xf8, 0x3f, 0xf0, 0x3f, 0xe0, 0x00, 0x00, 0x00, 0x00, 0x00,
  0x00, 0x03, 0xf0, 0x00, 0x0f, 0xc1, 0xfc, 0x00, 0x1f, 0x80, 0x03, 0xfc, 0x0f, 0x83, 0xfe, 0x08,
  0x7f, 0x83, 0xf0, 0x7f, 0xf8, 0x3f, 0xf0, 0x3f, 0xe0, 0x00, 0x00, 0x00, 0x00, 0x00, 0x00, 0x03,
  0xf0, 0x00, 0x0f, 0xc1, 0xfc, 0x00, 0x1f, 0x80, 0x03, 0xfc, 0x0f, 0x83, 0xfe, 0x08, 0x3f, 0x83,
  0xf0, 0x7f, 0xf8, 0x3f, 0xf0, 0x3f, 0xe0, 0x00, 0x00, 0x00, 0x00, 0x00, 0x00, 0x03, 0xf0, 0x00,
  0x0f, 0xc1, 0xfc, 0x00, 0x1f, 0x80, 0x03, 0xfc, 0x0f, 0x83, 0xfe, 0x18, 0x3f, 0x83, 0xe0, 0x7f,
  0xf8, 0x3f, 0xf0, 0x3f, 0xe0, 0x00, 0x00, 0x00, 0x00, 0x00, 0x00, 0x03, 0xf0, 0x00, 0x0f, 0xc1,
  0xfc, 0x00, 0x1f, 0x80, 0x03, 0xfc, 0x0e, 0x03, 0xfe, 0x18, 0x3f, 0x83, 0xc0, 0x7f, 0xf8, 0x3f,
  0xf0, 0x3f, 0xe0, 0x00, 0x00, 0x00, 0x00, 0x00, 0x00, 0x03, 0xf0, 0x00, 0x0f, 0xc1, 0xfc, 0x00,
  0x1f, 0x80, 0x03, 0xfc, 0x0c, 0x03, 0xfc, 0x18, 0x3f, 0x83, 0x80, 0xff, 0xf8, 0x3f, 0xf0, 0x3f,
  0xe0, 0x00, 0x00, 0x00, 0x00, 0x00, 0x00, 0x03, 0xf0, 0x00, 0x0f, 0xc1, 0xfc, 0x00, 0x1f, 0x80,
  0x03, 0xfc, 0x08, 0x03, 0xfc, 0x18, 0x3f, 0x83, 0x01, 0xff, 0xf8, 0x3f, 0xf0, 0x3f, 0xe0, 0x00,
  0x00, 0x00, 0x00, 0x00, 0x00, 0x03, 0xf0, 0x00, 0x0f, 0xc1, 0xfc, 0x00, 0x1f, 0x80, 0x03, 0xfc,
  0x00, 0x0f, 0xfc, 0x1c, 0x3f, 0x80, 0x03, 0xff, 0xf8, 0x3f, 0xf0, 0x3f, 0xe0, 0x00, 0x00, 0x00,
  0x00, 0x00, 0x00, 0x03, 0xf0, 0x00, 0x0f, 0xc1, 0xfc, 0x00, 0x1f, 0x80, 0x03, 0xfc, 0x00, 0x1f,
  0xfc, 0x1c, 0x1f, 0x80, 0x07, 0xff, 0xf8, 0x3f, 0xf0, 0x3f, 0xe0, 0x00, 0x00, 0x00, 0x00, 0x00,
  0x00, 0x03, 0xf0, 0x00, 0x0f, 0xc1, 0xfc, 0x00, 0x1f, 0x80, 0x03, 0xfc, 0x00, 0x3f, 0xfc, 0x1c,
  0x1f, 0x80, 0x07, 0xff, 0xf8, 0x3f, 0xf0, 0x3f, 0xe0, 0x00, 0x00, 0x00, 0x00, 0x00, 0x00, 0x03,
  0xf0, 0x00, 0x0f, 0xc1, 0xfc, 0x00, 0x1f, 0x80, 0x03, 0xfc, 0x00, 0xff, 0xfc, 0x3c, 0x1f, 0x80,
  0x07, 0xff, 0xf8, 0x3f, 0xf0, 0x3f, 0xe0, 0x00, 0x00, 0x00, 0x00, 0x00, 0x00, 0x03, 0xf0, 0x00,
  0x0f, 0xc1, 0xfc, 0x00, 0x1f, 0x80, 0x03, 0xfc, 0x01, 0xff, 0xf8, 0x3c, 0x1f, 0x80, 0x07, 0xff,
  0xf8, 0x3f, 0xf0, 0x3f, 0xe0, 0x00, 0x00, 0x00, 0x00, 0x00, 0x00, 0x03, 0xf0, 0x00, 0x0f, 0xc1,
  0xfc, 0x00, 0x1f, 0x80, 0x03, 0xfc, 0x03, 0xff, 0xf8, 0x3c, 0x1f, 0x81, 0x03, 0xff, 0xf8, 0x3f,
  0xf0, 0x3f, 0xe0, 0x00, 0x00, 0x00, 0x00, 0x00, 0x00, 0x03, 0xf0, 0x00, 0x0f, 0xc1, 0xfc, 0x00,
  0x1f, 0x80, 0x03, 0xfc, 0x0f, 0xff, 0xf8, 0x3c, 0x1f, 0x83, 0x83, 0xff, 0xf8, 0x3f, 0xf0, 0x3f,
  0xe0, 0x00, 0x00, 0x00, 0x00, 0x00, 0x00, 0x03, 0xf0, 0x00, 0x0f, 0xc1, 0xfc, 0x00, 0x1f, 0x80,
  0x03, 0xfc, 0x0f, 0xff, 0xf8, 0x3c, 0x0f, 0x83, 0x83, 0xff, 0xf8, 0x3f, 0xf0, 0x3f, 0xe0, 0x00,
  0x00, 0x00, 0x00, 0x00, 0x00, 0x03, 0xf0, 0x00, 0x0f, 0xc1, 0xfc, 0x00, 0x1f, 0x80, 0x03, 0xfc,
  0x0f, 0xff, 0xf8, 0x00, 0x0f, 0x83, 0x81, 0xff, 0xf8, 0x3f, 0xf0, 0x3f, 0xe0, 0x00, 0x00, 0x00,
  0x00, 0x00, 0x00, 0x03, 0xf0, 0x00, 0x0f, 0xc1, 0xfc, 0x00, 0x1f, 0x80, 0x03, 0xfc, 0x0f, 0xff,
  0xf8, 0x00, 0x0f, 0x83, 0xc1, 0xff, 0xf8, 0x3f, 0xf0, 0x3f, 0xe0, 0x00, 0x00, 0x00, 0x00, 0x00,
  0x00, 0x03, 0xf0, 0x00, 0x0f, 0xc1, 0xfc, 0x00, 0x1f, 0x80, 0x03, 0xfc, 0x0f, 0xff, 0xf0, 0x00,
  0x0f, 0x83, 0xc1, 0xff, 0xf8, 0x3f, 0xf0, 0x3f, 0xe0, 0x00, 0x00, 0x00, 0x00, 0x00, 0x00, 0x03,
  0xf0, 0x00, 0x0f, 0xc0, 0xfc, 0x00, 0x3f, 0x80, 0x03, 0xfc, 0x0f, 0xff, 0xf0, 0x00, 0x0f, 0x83,
  0xc0, 0xff, 0xf8, 0x3f, 0xf0, 0x3f, 0xe0, 0x00, 0x00, 0x00, 0x00, 0x00, 0x00, 0x03, 0xf0, 0x00,
  0x0f, 0xc0, 0xfe, 0x00, 0x7f, 0x80, 0x03, 0xfc, 0x0f, 0xff, 0xf0, 0x00, 0x0f, 0x83, 0xc0, 0xff,
  0xf8, 0x3f, 0xf0, 0x3f, 0xe0, 0x00, 0x00, 0x00, 0x00, 0x00, 0x00, 0x03, 0xff, 0xfe, 0x0f, 0xc0,
  0xff, 0xff, 0xff, 0x00, 0x03, 0xfc, 0x0f, 0xff, 0xf0, 0x00, 0x07, 0x83, 0xe0, 0xff, 0xf8, 0x3f,
  0xf0, 0x3f, 0xe0, 0x00, 0x00, 0x00, 0x00, 0x00, 0x00, 0x03, 0xff, 0xfe, 0x0f, 0xc0, 0x7f, 0xff,
  0xff, 0x00, 0x03, 0xfc, 0x0f, 0xff, 0xf0, 0x7f, 0x07, 0x83, 0xe0, 0xff, 0xf8, 0x3f, 0xf0, 0x3f,
  0xe0, 0x00, 0x00, 0x00, 0x00, 0x00, 0x00, 0x03, 0xff, 0xfe, 0x0f, 0xc0, 0x7f, 0xff, 0xfe, 0x00,
  0x03, 0xfc, 0x0f, 0xff, 0xe0, 0x7f, 0x07, 0x83, 0xe0, 0x7f, 0xf8, 0x3f, 0xf0, 0x3f, 0xe0, 0x00,
  0x00, 0x00, 0x00, 0x00, 0x00, 0x03, 0xff, 0xfe, 0x0f, 0xc0, 0x3f, 0xff, 0xfc, 0x00, 0x03, 0xfc,
  0x0f, 0xff, 0xe0, 0xff, 0x07, 0x83, 0xe0, 0x7f, 0xf8, 0x3f, 0xf0, 0x3f, 0xe0, 0x00, 0x00, 0x00,
  0x00, 0x00, 0x00, 0x03, 0xff, 0xfe, 0x0f, 0xc0, 0x1f, 0xff, 0xf8, 0x00, 0x03, 0xfc, 0x0f, 0xff,
  0xe0, 0xff, 0x07, 0x83, 0xf0, 0x7f, 0xf8, 0x3f, 0xf0, 0x3f, 0xe0, 0x00, 0x00, 0x00, 0x00, 0x00,
  0x00, 0x03, 0xff, 0xfe, 0x0f, 0xc0, 0x03, 0xff, 0xe0, 0x00, 0x03, 0xfc, 0x0f, 0xff, 0xe0, 0xff,
  0x07, 0x83, 0xf0, 0x3f, 0xf8, 0x3f, 0xf0, 0x3f, 0xe0, 0x00, 0x00, 0x00, 0x00, 0x00, 0x00, 0x00,
  0x00, 0x00, 0x00, 0x00, 0x00, 0x3f, 0xe0, 0x00, 0x03, 0xfc, 0x0f, 0xff, 0xe0, 0xff, 0x03, 0x83,
  0xf0, 0x3f, 0xf8, 0x3f, 0xf0, 0x3f, 0xe0, 0x00, 0x00, 0x00, 0x00, 0x00, 0x00, 0x00, 0x00, 0x00,
  0x00, 0x00, 0x00, 0x1f, 0xf0, 0x00, 0x03, 0xff, 0xff, 0xff, 0xff, 0xff, 0xff, 0xff, 0xff, 0xff,
  0xff, 0xff, 0xff, 0xff, 0xe0, 0x00, 0x00, 0x00, 0x00, 0x00, 0x00, 0x00, 0x00, 0x00, 0x00, 0x00,
  0x00, 0x0f, 0xf0, 0x00, 0x03, 0xff, 0xff, 0xff, 0xff, 0xff, 0xff, 0xff, 0xff, 0xff, 0xff, 0xff,
  0xff, 0xff, 0xe0, 0x00, 0x00, 0x00, 0x00, 0x00, 0x00, 0x00, 0x00, 0x00, 0x00, 0x00, 0x00, 0x07,
  0xe0, 0x00, 0x03, 0xff, 0xff, 0xff, 0xff, 0xff, 0xff, 0xff, 0xff, 0xff, 0xff, 0xff, 0xff, 0xff,
  0xe0, 0x00, 0x00, 0x00, 0x00, 0x00, 0x00, 0x00, 0x00, 0x00, 0x00, 0x00, 0x00, 0x03, 0xc0, 0x00,
  0x03, 0xff, 0xff, 0xff, 0xff, 0xff, 0xff, 0xff, 0xff, 0xff, 0xff, 0xff, 0xff, 0xff, 0xe0, 0x00,
  0x00, 0x00, 0x00, 0x00, 0x00, 0x00, 0x00, 0x00, 0x00, 0x00, 0x00, 0x01, 0x80, 0x00, 0x03, 0xff,
  0xff, 0xff, 0xff, 0xff, 0xff, 0xff, 0xff, 0xff, 0xff, 0xff, 0xff, 0xff, 0xe0, 0x00, 0x00, 0x00,
  0x00, 0x00, 0x00, 0x00, 0x00, 0x00, 0x00, 0x00, 0x00, 0x00, 0x00, 0x00, 0x03, 0xff, 0xff, 0xff,
  0xff, 0xff, 0xff, 0xff, 0xff, 0xff, 0xff, 0xff, 0xff, 0xff, 0xe0, 0x00, 0x00, 0x00, 0x00, 0x00,
  0x00, 0x00, 0x00, 0x00, 0x00, 0x00, 0x00, 0x00, 0x00, 0x00, 0x03, 0xff, 0xff, 0xff, 0xff, 0xff,
  0xff, 0xff, 0xff, 0xff, 0xff, 0xff, 0xff, 0xff, 0xe0, 0x00, 0x00, 0x00, 0x00, 0x00, 0x00, 0x00,
  0x00, 0x00, 0x00, 0x00, 0x00, 0x00, 0x00, 0x00, 0x00, 0x00, 0x00, 0x00, 0x00, 0x00, 0x00, 0x00,
  0x00, 0x00, 0x00, 0x00, 0x00, 0x00, 0x00, 0x00, 0x00, 0x00, 0x00, 0x00, 0x00, 0x00, 0x00, 0x00,
  0x00, 0x00, 0x00, 0x00, 0x00, 0x00, 0x00, 0x00, 0x00, 0x00, 0x00, 0x00, 0x00, 0x00, 0x00, 0x00,
  0x00, 0x00, 0x00, 0x00, 0x00, 0x00, 0x00, 0x00
};
```
