## Extended Data 1 for "A novel mouse home cage lickometer system reveals sex- and housing-based influences on alcohol drinking": SD_RTC_functions.ino

```
//==========================================================================================

void start_rtc() {

  // connect to rtc
  Wire.begin();
  Wire.setClock(3400000);
  Wire.setWireTimeout();

  if (!rtc.begin()) {
    logfile.println(F("rtc failed"));
    DEBUG_System_PRINTLN(F("rtc failed"));
  }

  if (!rtc.initialized() || rtc.lostPower()) {
    DEBUG_System_PRINTLN("RTC syncing to computer clock.");
    // following line sets the RTC to the date & time this sketch was compiled
    rtc.adjust(DateTime(__DATE__, __TIME__));
  }
}

//==========================================================================================

bool start_rfid() {

  if (!myRfid1.begin()) {
    error("RFID Reader #1 not found, check wiring?");
    refresh_page = true;
    while (!myRfid1.begin()) {
      resetWire();
      if (!rtc.begin()) {
        logfile.println(F("rtc failed"));
        DEBUG_System_PRINTLN(F("rtc failed"));
      }
      if (ts.touched()) {
        TS_Point p = ts.getPoint();
        wait(100);
        p.x = map(p.x, 0, 240, 0, 240);
        p.y = map(p.y, 0, 320, 0, 320);
        if (p.x > 10 && p.x < 230 && p.y > 220 && p.y < 280) {
          p = {};
          display_page = "main";
          refresh_page = true;
          break;
        }
      }
    }
    start_display();
    tft.fillScreen(ILI9341_RED);
    return true;
  }

  if (!myRfid2.begin()) {
    error("RFID Reader #2 not found, check wiring?");
    refresh_page = true;
    while (!myRfid2.begin()) {
      resetWire();
      if (!rtc.begin()) {
        logfile.println(F("rtc failed"));
        DEBUG_System_PRINTLN(F("rtc failed"));
      }
      if (ts.touched()) {
        TS_Point p = ts.getPoint();
        wait(100);
        p.x = map(p.x, 0, 240, 0, 240);
        p.y = map(p.y, 0, 320, 0, 320);
        if (p.x > 10 && p.x < 230 && p.y > 220 && p.y < 280) {
          p = {};
          display_page = "main";
          refresh_page = true;
          break;
        }
      }
    }
    start_display();
    tft.fillScreen(ILI9341_RED);
    return true;
  }
}

//======================================================================================

void read_tag_int() {
  if (digitalRead(intPin1) == LOW) {
    tag1 = myRfid1.getTag();
    DEBUG_RFID_PRINT("LEFT Tag: ");
    DEBUG_RFID_PRINTLN(tag1);
    myRfid1.clearTags();
  }
  if (digitalRead(intPin2) == LOW) {
    tag2 = myRfid2.getTag();
    DEBUG_RFID_PRINT("RIGHT Tag: ");
    DEBUG_RFID_PRINTLN(tag2);
    myRfid2.clearTags();
  }
}

//======================================================================================

void resetWire() {
  TWCR = 0;
  Wire.end();
  Wire.begin();
  Wire.setClock(3400000);
  Wire.setWireTimeout();
}

//==========================================================================================

void make_file() {
  // fetch the time
  DateTime now;
  now = rtc.now();

  SdFile::dateTimeCallback(dateTime_);

  filename[0] = (now.year() / 10) % 10 + '0';  //To get 3rd digit from year()
  filename[1] = now.year() % 10 + '0';         //To get 4th digit from year()
  filename[2] = now.month() / 10 + '0';        //To get 1st digit from month()
  filename[3] = now.month() % 10 + '0';        //To get 2nd digit from month()
  filename[4] = now.day() / 10 + '0';          //To get 1st digit from day()
  filename[5] = now.day() % 10 + '0';          //To get 2nd digit from day()

  for (uint8_t i = 0; i < 100; i++) {
    filename[6] = i / 10 + '0';
    filename[7] = i % 10 + '0';
    if (!SD.exists(filename)) {
      // only open a new file if it doesn't exist
      logfile = SD.open(filename, FILE_WRITE);
      break;  // leave the loop!
    }
  }
  if (!logfile) {
    error(F("Could not create file"));
  }

  DEBUG_LIQ_PRINT(F("Logging to: "));
  DEBUG_LIQ_PRINTLN(filename);
  nowloggingto();
  logfile.flush();

  // Write headers to file
  if (!logfile.println(F("Experimental_side,millis_since_start,datetime,MouseRFID,BottleLicked,BoutDuration,BoutLickNumber,BoutLickDuration,LickFrequency,Inter-Lick-Interval"))) {
    error(F("Could not write file"));
  }

  //echo to serial monitor
  DEBUG_LIQ_PRINTLN(F("Experimental_side,millis_since_start,datetime,MouseRFID,BottleLicked,BoutDuration,BoutLickNumber,BoutLickDuration,LickFrequency,Inter-Lick-Interval"));

  syncTime = now.minute();
  SYNC_COUNTER = 0;
}

//==========================================================================================

void nowloggingto() {
  tft.fillRect(0, 20, 240, 320, ILI9341_RED);
  tft.setFont(&FreeSansBold12pt7b);
  tft.setTextColor(ILI9341_WHITE);
  NickText_center("Logging to: ", 120, 120, 1);
  NickText_center(filename, 120, 150, 1);  //show what file is being recorded to (filename)
  tft.setFont(&FreeSans12pt7b);
  NickText_center("Calibrating Sensors", 120, 200, 1);
  NickText_center("Stand Back!", 120, 230, 1);
  wait(2000);  //wait 2000ms
}

//==========================================================================================

void logging_same_file() {
  tft.fillRect(0, 20, 240, 320, ILI9341_RED);
  tft.setFont(&FreeSans12pt7b);
  tft.setTextColor(ILI9341_WHITE);
  NickText_center("Logging to same file: ", 120, 120, 1);
  NickText_center(filename, 120, 150, 1);  //show what file is being recorded to (filename)
  NickText_center("Calibrating Sensors", 120, 200, 1);
  NickText_center("Stand Back!", 120, 230, 1);
  wait(2000);  //wait 2000ms
}

//======================================================================================

void write_to_file(int p) {
  // fetch the time
  now = rtc.now();
  uint32_t m = millis() - startmillis;

  logfile.print(E_side);  //log Experimental Side
  logfile.print(F(", "));

  logfile.print(m);  // log milliseconds since starting
  logfile.print(F(", "));

  logfile.print('"');  // log time
  logfile.print(now.year(), DEC);
  logfile.print(F("/"));
  logfile.print(now.month(), DEC);
  logfile.print(F("/"));
  logfile.print(now.day(), DEC);
  logfile.print(F(" "));
  logfile.print(now.hour(), DEC);
  logfile.print(F(":"));
  if (now.minute() < 10) {
    logfile.print("0");
  }
  logfile.print(now.minute(), DEC);
  logfile.print(F(":"));
  if (now.second() < 10) {
    logfile.print("0");
  }
  logfile.print(now.second(), DEC);
  logfile.print('"');
  logfile.print(F(", "));

  if (p == 0) {
    licked_bottle = "LEFT";
    if (tag1 != "") {
      if (!RFIDerrorLEFT && !RFID_lasterrorLEFT) {
        tag1_count += 1;
      } else {
        tag1_count = 1;
      }
      logfile.print(tag1);
      logfile.print(F(", "));
      RFIDerrorLEFT = false;
      RFID_lasterrorLEFT = false;
    } else if (tag1 == "" && lasttag1 == "") {  //no tags have been read!
      RFIDerrorLEFT = true;
      RFID_lasterrorLEFT = false;
      notag1_count += 1;
      start_rfid();
    } else if (tag1 == "" && lasttag1 != "") {  //most recent tag was not read
      if (RFID_lasterrorLEFT) {
        lasttag1_count += 1;
      } else {
        lasttag1_count = 1;
      }
      logfile.print(lasttag1);
      logfile.print(F(", "));
      start_rfid();
      RFID_lasterrorLEFT = true;
      RFIDerrorLEFT = false;
    }
  }

  if (p == 1) {
    licked_bottle = "RIGHT";
    if (tag2 != "") {
      if (!RFIDerrorRIGHT && !RFID_lasterrorRIGHT) {
        tag2_count += 1;
      } else {
        tag2_count = 1;
      }
      logfile.print(tag2);
      logfile.print(F(", "));
      RFIDerrorRIGHT = false;
      RFID_lasterrorRIGHT = false;
    } else if (tag2 == "" && lasttag2 == "") {  //no tags have been read!
      RFIDerrorRIGHT = true;
      RFID_lasterrorRIGHT = false;
      notag2_count += 1;
      start_rfid();
    } else if (tag2 == "" && lasttag2 != "") {  //most recent tag was not read
      if (RFID_lasterrorRIGHT) {
        lasttag2_count += 1;
      } else {
        lasttag2_count = 1;
      }
      logfile.print(lasttag2);
      logfile.print(F(", "));
      start_rfid();
      RFID_lasterrorRIGHT = true;
      RFIDerrorRIGHT = false;
    }
  }

  logfile.print(licked_bottle);
  logfile.print(F(", "));

  logfile.print(BoutDuration[p]);
  logfile.print(F(", "));

  logfile.print(BoutLickNumber[p]);
  logfile.print(F(", "));

  logfile.print(BoutLickDuration[p]);
  logfile.print(F(", "));

  logfile.print(LickFrequency[p]);
  logfile.print(F(", "));

  logfile.print(ILI[p]);
  logfile.println();  // this is going to next line

  DEBUG_LIQ_PRINT(E_side);
  DEBUG_LIQ_PRINT(F(", "));

  DEBUG_LIQ_PRINT(m);
  DEBUG_LIQ_PRINT(F(", "));

  DEBUG_LIQ_PRINT('"');
  DEBUG_LIQ_PRINTDEC(now.year());
  DEBUG_LIQ_PRINT(F("/"));
  DEBUG_LIQ_PRINTDEC(now.month());
  DEBUG_LIQ_PRINT(F("/"));
  DEBUG_LIQ_PRINTDEC(now.day());
  DEBUG_LIQ_PRINT(F(" "));
  DEBUG_LIQ_PRINTDEC(now.hour());
  DEBUG_LIQ_PRINT(F(":"));
  if (now.minute() < 10) {
    DEBUG_LIQ_PRINT("0");
  }
  DEBUG_LIQ_PRINTDEC(now.minute());
  DEBUG_LIQ_PRINT(F(":"));
  if (now.second() < 10) {
    DEBUG_LIQ_PRINT("0");
  }
  DEBUG_LIQ_PRINTDEC(now.second());
  DEBUG_LIQ_PRINT('"');
  DEBUG_LIQ_PRINT(F(", "));

  if (p == 0) {
    DEBUG_LIQ_PRINT(tag1);
    DEBUG_LIQ_PRINT(F(", "));
  }
  if (p == 1) {
    DEBUG_LIQ_PRINT(tag2);
    DEBUG_LIQ_PRINT(F(", "));
  }
  DEBUG_LIQ_PRINT(licked_bottle);
  DEBUG_LIQ_PRINT(F(", "));

  DEBUG_LIQ_PRINT(BoutDuration[p]);
  DEBUG_LIQ_PRINT(F(", "));

  DEBUG_LIQ_PRINT(BoutLickNumber[p]);
  DEBUG_LIQ_PRINT(F(", "));

  DEBUG_LIQ_PRINT(BoutLickDuration[p]);
  DEBUG_LIQ_PRINT(F(", "));

  DEBUG_LIQ_PRINT(LickFrequency[p]);
  DEBUG_LIQ_PRINT(F(", "));

  DEBUG_LIQ_PRINT(ILI[p]);
  DEBUG_LIQ_PRINTLN();

  if (p == 0 && tag1 != "") {
    lasttag1 = tag1;
    tag1 = "";
  }
  if (p == 1 && tag2 != "") {
    lasttag2 = tag2;
    tag2 = "";
  }

  if (RFIDerrorLEFT) {
    DEBUG_LIQ_PRINT("LEFT RFID sensor did not detect a tag!")
  }
  if (RFIDerrorRIGHT) {
    DEBUG_LIQ_PRINT("RIGHT RFID sensor did not detect a tag!")
  }

  if (RFIDerrorRIGHT || RFIDerrorLEFT) {
    display_licks();
  }

  //
  //  Serial.print("Time it took to write to file: ");
  //  Serial.print(millis() - m);
  //  Serial.println("ms");
}

//======================================================================================

void write_data() {
  if (SYNC_COUNTER > SYNC_INTERVAL) {
    uint32_t m = millis();
    now = rtc.now();
    syncTime = now.minute();
    SYNC_COUNTER = 0;
    DEBUG_System_PRINT("Writing data to SD card!");
    digitalWrite(redLEDpin, HIGH);  // blink LED to show we are syncing data to the card & updating FAT!
    logfile.flush();
    digitalWrite(redLEDpin, LOW);
    //    Serial.print("Time it took to sync to SD card: ");
    //    Serial.print(millis() - m);
    //    Serial.println("ms");
  }
}

//==========================================================================================

void dateTime_(uint16_t* date, uint16_t* Time) {
  *date = FAT_DATE(now.year(), now.month(), now.day());
  *Time = FAT_TIME(now.hour(), now.minute(), now.second());
}
```
